# Venomics and antivenomics of Indian spectacled cobra (*Naja naja*) from the Western Ghats

**DOI:** 10.1101/2021.02.15.431263

**Authors:** Muralidharan Vanuopadath, Dileepkumar Raveendran, Bipin Gopalakrishnan Nair, Sudarslal Sadasivan Nair

## Abstract

Venom proteome profiling of *Naja naja* from the Western Ghats region in Kerala was achieved through SDS-PAGE and RP-HPLC followed by Q-TOF LC-MS/MS analysis, incorporating PEAKS and Novor assisted *de novo* sequencing methodologies. A total of 115 proteins distributed across 17 different enzymatic and non-enzymatic venom protein families were identified through conventional and 39 peptides through homology-driven proteomics approaches. Fourteen peptides derived through *de novo* complements the Mascot data indicating the importance of homology-driven approaches in improving protein sequence information. Among the protein families identified, glutathione peroxidase and endonuclease were reported for the first time in the Indian cobra venom. Immunological cross-reactivity assessed using Indian polyvalent antivenoms suggested that VINS showed better EC_50_ (2.48 μg/mL) value than that of PSAV (6.04 μg/mL) and Virchow (6.03 μg/mL) antivenoms. Western blotting experiments indicated that all the antivenoms elicited poor binding specificities, especially towards low molecular mass proteins. Second-generation antivenomics studies revealed that VINS antivenom was less efficient to detect many low molecular mass proteins such as three-finger toxins and Kunitz-type serine protease Inhibitors. Taken together, the present study enabled a large-scale characterization of the venom proteome of *Naja naja* from the Western Ghats and emphasized the need for developing more efficient antivenoms.

**Highlights:**

- Proteomics of cobra venom resulted in the identification of 115 proteins representing 17 snake venom protein families.
- *De novo* approaches exclusively yielded 39 peptides harbouring multiple amino acid mutations.
- Glutathione peroxidase and endonuclease were identified for the first time in Indian cobra venom.
- Indian polyvalent antivenoms showed varying cross-reactivity towards cobra venom.
- VINS antivenom was less efficient to detect many low molecular mass proteins (< 20 kDa).

## 1. Introduction

Snakebite envenoming is a major threat to the people living in the tropical regions of the world. It is noted that approximately half of global snakebite deaths are reported from India. Studies also showed that around 1.2 million people died in India due to snakebites, that too within a short period of about two decades (2000-2019), resulting in approximately 58,000 deaths per year [1]. It is equally disturbing to note that apart from the massive mortality rate, a three-fold increase in morbidity cases accounts for several limb amputations and other permanent disabilities [2]. Due to the degree of severity, chances of occurrence, lack of proper treatment, high costs involved in treating envenomed victims and increased mortality and morbidity rates, snakebite envenoming has been listed under the neglected tropical disease category by the World Health Organization (WHO) in June 2017 [3]. In India, spectacled cobra (*Naja naja*), common krait (*Bungarus caeruleus*), saw-scaled viper (*Echis carinatus*) and Russell’s viper (*Daboia russelii*), commonly known as the ‘Big Four’ venomous snakes, account for majority of the morbidity and mortality cases [4].

The Indian cobra belongs to the genus Naja and family Elapidae is a highly venomous snake species. This species is native to countries of the Indian subcontinent including India, Sri Lanka, Bangladesh and Nepal [5]. They are seen in wide habitats like plains, dense or open forests, rocky terrains, wetlands, agricultural lands, outskirts of villages and even in highly populated urban areas. The species is also known under different names such as spectacled cobra, Asian cobra and binocellate cobra due to the presence of hoods behind the neck [6]. Snake venoms belonging to Elapidae family are rich in proteins that include three-finger toxins (3FTxs) and phospholipase A_2_s (PLA_2_s). Though these proteins are smaller in size, they are highly toxic and are responsible for most of the morbidity and mortality cases associated with the envenomation. The major contributory proteins in the three-finger toxins family include post-synaptic neurotoxins, cardiotoxins/cytotoxins and muscarinic toxins [7]. Since *Naja naja* is distributed throughout India, studies were conducted to understand constituent proteins of *Naja naja* venom collected from different geographical locations and the data clearly indicate intra-species variation in the venom composition [5, 8–14]. The observed difference can be correlated to multiple factors including the geographical and prey-specific variations [15, 16]. Irrespective of all these, it is interesting to note that cobra venom from all the geographical areas are predominant with proteins belonging to three-finger toxin class.

Currently, mass spectrometry follows two principal approaches, top-down and bottom-up, for the structural characterization of venom components. The former deals with the in-depth analysis of proteins through gas phase fragmentation to establish their identity, identify post-translational modifications and characterise various isoforms [17, 18]. Whereas the latter involves proteolytic digestion of proteins into peptide fragments before mass spectrometric analysis. Since the bottom-up proteomics approaches use the information stored in public database repositories including NCBI and Uniprot, the success rate of protein identification through this strategy totally relies on the protein sequence information present in the database. However, due to the lack of inadequate protein sequence information in public repositories, in-depth characterisation of snake venom proteome remains a challenging one. Previous studies from our laboratory on venom proteome of various snake species had clearly demonstrated the importance of homology-assisted sequencing strategies in elucidating the venom profiles of snake species, whose sequence information is under-represented in the databases [19, 20]. In addition to this, the advantages of *de novo* sequencing followed by MS-BLAST and PEAKS analysis over conventional proteomics approaches have been substantiated by multiple reports [21–23].

Snake venomics enable characterising the venom arsenal of various snake species and antivenomics on the other hand utilise proteome information for extending the possibilities in improving antivenom preparations. Antivenomics could be also used for assessing the immunological cross-reactivity between homologous toxins present within various snake species [24]. Since antivenoms are the only choice available for treating envenomed victims, there is a need in improving the efficacy of the available antivenoms [25, 26]. Polyvalent antivenoms in India are manufactured using the pooled venom collected from the ‘big four’ snake species which includes *Naja naja*. Still the commercial antivenoms from the major Indian manufacturers showed differences in immunological reactivity profiles towards *Naja naja* venom collected from different geographical locations in India [8, 10, 11, 13, 14]. For example, though Indian polyvalent antivenoms contain enough antibodies against antigenic epitopes present in the cobra venom, they often fail to neutralize the low-molecular mass toxins such as 3FTxs and PLA_2_s [8, 10]. Similarly, cobra venom from different geographical locations exhibited varied immunological cross-reactivity profiles to the polyvalent antivenoms used [8, 11].

We report here, new observations on the venom proteome of *Naja naja* venom from the Western Ghats region in Kerala through orthogonal separation and identification strategies. Also, we have evaluated the specificity and binding potential of commercially available antivenoms towards *Naja naja* venom through WHO approved in vitro preclinical protocols.

## 2. Materials and methods

### 2.1 Reagents and chemicals

Formic acid (FA), methanol, acetonitrile (ACN), trifluoroacetic acid (TFA), acrylamide, bisacrylamide, ammonium persulphate, coomassie brilliant blue R-250 (CBB R-250), beta-mercaptoethanol (β-ME), bromophenol blue, glycerol, glycine, iodoacetamide (IAm), ammonium bicarbonate, Tetramethyl benzidine/Hydrogen Peroxide (TMB/H_2_O_2),_ HRP-conjugated secondary antibody, tetramethylethylenediamine, dithiothreitol (DTT), trypsin, chymotrypsin and V8 proteases were purchased from Sigma-Aldrich. ECL substrate, Tris, SDS and PVDF membrane were procured from Bio-Rad. NHS-activated Sepharose 4 Fast Flow was obtained from GE Healthcare and protein ladder from Thermo Fisher Scientific.

### 2.2 Venom collection

Venom from two snakes belong to *Naja naja* species from the Western Ghats region in Kerala, one of the states in India, (NNwg) was collected in captive. Venom milking was done in-house (as per the Government order, GO (Rt) No. 94/2009/F & WLD dated 25/02/2009) in the Government serpentarium (Thiruvananthapuram, Kerala, India) by an experienced herpetologist. Followed by lyophilization, venom was stored at −80 °C until analysis.

### 2.3 Antivenoms

Lyophilized equine antivenoms raised against the ‘Big Four’ snakes of India (*Echis carinatus*, *Naja naja*, *Bungarus caeruleus*, and *Daboia russelli*) obtained from VINS (batch number: 01AS16040, expiry date: 07/2020), Virchow (batch number: PAS00116, expiry date: 01/2020) and Premium Serums and Vaccines (PSAV; batch number: ASVS (I) Ly-010, expiry date: 08/2022) were used for the entire immunological cross-reactivity assays.

### 2.4. Protein estimation using BCA assay

Protein estimation of crude venom and antivenom samples was performed using standard BCA assay [27]. For this, BSA with known concentrations was serially diluted from 2 mg/mL stock to get working concentrations of 1.5, 1.0, 0.75, 0.5, 0.25, 0.125, 0.025, and 0.03125 mg/mL. 10 μL from each of these standards was then mixed with 200 μL of BCA reagent (A and B solutions in a 1:50 ratio) and kept for incubation (30 min at 37 °C). The values were recorded at 562 nm using a plate reader.

### 2.5 SDS-PAGE and In-gel digestions of crude NNwg venom

12% SDS gel was used to resolve the crude NNwg venom (100 μg) proteins. The bands obtained were stained and visualized using CBB R-250 stain followed by scanning using a gel doc system (Bio-Rad). The gels bands separated under reducing conditions were then processed for in-gel digestion procedures. For this, the gel bands were cut into small pieces and excess stain was removed through 25 mM ammonium bicarbonate buffer and 50% ACN washing. This step was repeated until the gel bands were clear. Subsequently, the gel pieces were subjected to reduction using 100 mM DTT. The reaction was performed at 56 °C for 45 min. After incubation, the tube was allowed to cool at room temperature and excess of DTT was aspirated. Further, alkylation was performed using IAm (250 mM in ammonium bicarbonate buffer, pH 8.0). This was kept for incubation in the dark at room temperature for 30 min. After this, 25 mM ammonium bicarbonate buffer solution was used to wash the gel pieces for removing any leftover IAm. Subsequently, the gel pieces were dehydrated using 100% ACN for removing water content. Followed by this, in-gel digestion was performed using trypsin, chymotrypsin and V8 protease on three different gel preparations. For trypsin digestion, 25 μL of enzyme (from 12.5 ng/μL stock) was added to the gel pieces after dehydration (using 100% ACN). Approximately, 200 ng of chymotrypsin or V8 protease were added on to the individual gel pieces for the respective digestion processes. All the three enzymatic digestions were allowed to proceed at 37 °C for 16 h. The extraction of the digested peptides was carried-out using ACN and water containing 0.1 % TFA. The extraction step was repeated twice, and the resulting supernatants were pooled and were completely dried using a vacuum concentrator. The samples resuspended in 5% ACN solution were subjected to mass spectrometric analysis or stored at −80 °C until analysis.

### 2.6 Reversed-phase HPLC of NNwg venom

3.2 mg of crude venom was dissolved in 80 μL of Milli-Q water and was centrifuged at 12000 rpm for 15 min at 4 °C. From this, 5 μL was loaded on to a reversed-phase column (Agilent Zorbax SB C8, 150 x 2.1 mm, 3.5 μM) through an Agilent 1260 HPLC system for the separation of various venom components. Solvent (A: water + 0.1% FA; B: ACN + 0.1% FA) flowrate was maintained at 200 μL/min. The following time program was used for the fraction collection: 5% B for 5 min, 5-15 % for 10 min, 15-45 % B for 60 min, 45-70 % B for 12 min, 70-95% B for 5 min and equilibration for 13 min. Absorbance was monitored at 210 nm and the peaks were manually collected and stored.

### 2.7 In-gel and In-solution enzymatic digestions of HPLC separated protein fractions

In-gel digestion of RP-HPLC separated fractions were performed as mentioned above (section 2.4). The fractions collected after reversed-phase HPLC was subjected to in-solution enzymatic digestions also. pH of the samples was adjusted to 8.0 using 50 mM ammonium bicarbonate buffer prior to in-solution digestion. Initially the samples were subjected to reduction using 100 mM DTT. This was then kept for incubation at 56 °C for 45 min. Alkylation was performed on the resulting solution using IAm (250 mM in ammonium bicarbonate buffer). The tubes were incubated at room temperature in the dark for 30 min. Followed by this, trypsin, chymotrypsin or V8 protease were added and kept for overnight incubation at 37 °C. The resultant solutions were acidified and dried completely using a vacuum concentrator.

### 2.8 Tandem mass spectrometry

The digested samples were reconstituted in 5% ACN containing 0.1% FA. Sample analysis was performed using an Agilent 6540 Q-TOF LC-MS/MS equipped with a chip cube source. The samples were infused to the MS through an HPLC system containing capillary and nano pumps that deliver solvents to the enrichment (40 nL) and analytical columns (Zorbax 300 SB-C18, 150 mm x 75 μm, 5 μ) present in the chip, respectively. The solvent (A: water with 0.1% formic aid; B: 90% acetonitrile with 0.1% formic acid) flowrate was maintained at 3 μL/min for the capillary and 0.3μL/min for the nano pump. The gradient conditions used for the separation of the peptide samples were as mentioned: initial equilibration using 5% B for 2 min, 5-45 % B in 30 min, 45-90% B in 45 min, 90% B for 2 min. All the MS data were collected in a data-dependent and positive ionisation mode using nitrogen as the collision gas. The MS parameters used for the data acquisition were as follows: MS range-250-3000 m/z; MS/MS-50-3000 m/z; Capillary voltage: 2000 V; Dry gas temperature: 325 °C; Dry gas flow: 4 L/min; Fragmentation voltage: 150 V; MS and MS/MS scan speeds were set at 4 spectra/sec and 3 spectra/sec, respectively. A maximum of five precursors were selected for the fragmentation. The charge states for tryptic digested samples were selected as 2, 3, >3 and for chymotrypsin and V8 digests: 1, 2, 3, >3. Active exclusion was enabled after 3 spectra with a releasing time of 0.5 min. MassHunter software (Version 6.01) from Agilent was used for the data acquisition and analysis.

### 2.9 Identification and validation of proteins using Mascot and Scaffold

The mgf files generated from the raw mass spectrometric data were analysed using Mascot (version 2.5.1, Matrix Science) through Mascot Daemon (version 2.5.0) interface with the following parameters: Database: Serpentes (Taxonomy ID:8570); peptide tolerance: 10 ppm, MS/MS tolerance: 0.5 Da; amino acid modifications: cysteine (carbamidomethylation as fixed modification), glutamine, asparagine (deamidation) and methionine (oxidation) as variable modifications.; number of missed cleavages: 2; Enzymes (selected charge states), trypsin (2^+^, 3^+^, 4^+^), chymotrypsin and V8 protease (1^+^, 2^+^ and 3^+^); the database searches were decoy enabled. The obtained data were further validated using Scaffold software (version 4.5.0, Proteomes Software Inc.) by setting the protein and peptide identification probabilities to 99% and 95%, respectively [28, 29]. The data were filtered using the inbuilt peptide and protein prophet algorithms and those proteins which were represented by a minimum of 2 peptides were included for the protein identification and classification process.

### 2.10 Protein identification through PEAKS

The obtained MS data was also subjected to protein identification using PEAKS Studio (version 8.5, Bioinformatics Solutions Inc., Canada) software. For the analysis, the precursor mass tolerance was set at 10 ppm and the fragment mass tolerance was set at 0.5 Da. Carbamidomethylation of cysteine was set as the fixed modification and deamidation of asparagine and glutamine along with the oxidation of methionine as variable modifications. Trypsin, chymotrypsin or V8 protease digestion options were selected according to the enzymes used. The confidently identified sequence tags (achieved by fixing the average local confidence to ≥30%) were then searched against Serpentes database. The peptides derived through *de novo* analysis were further validated through BLAST searches and were then manually verified [19].

### 2.11 *De novo* sequencing through Novor and MS-BLAST

In order to identify peptides with possible amino acid mutations, Novor followed by MS-BLAST sequence similarity searches were performed. For this, the raw data converted to mgf format was subjected to Novor analysis using the DeNovoGUI (Version 1.15.11) user interface [30]. Parameters used for the identification of proteins and peptides are: 10 ppm and 0.5 Da error tolerance for precursor and fragment masses, respectively; amino acid modifications: cysteine (carbamidomethylation as fixed modification), glutamine, asparagine (deamidation) and methionine (oxidation) as variable modifications. The obtained data after Novor analysis were pre-processed manually and subjected to MS-BLAST [31] analysis using Serpentes database as described previously [20].

### 2.12 BLAST search

Peptides with mutations obtained through *de novo* approaches using PEAKS and Novor followed by MS-BLAST were validated through BLAST search. For this, the obtained sequences were searched against the non-redundant (nr) database from NCBI (blastp suite at www.ncbi.nlm.nih.gov/BLAST) using the taxonomy, Serpentes (taxid: 8570) [32]. Following BLAST, the identified peptide (s) representing various snake venom proteins were listed using the score and expectation (E) values obtained. Ideally, peptides with the lowest E-values were taken further. For matches with similar E-values and hit scores, proteins belonging to the same family (though they are having different accession numbers) and different snake species, the top hit was considered for the representation purpose. Also, if a *de novo* derived peptide represent a protein that was already identified through conventional database assisted proteomic analysis, the peptide was assigned to that particular protein for improving the sequence coverage. All the obtained sequences were then manually verified from the tandem mass spectra.

### 2.13 ClustalW

For confirming the sequences obtained for newly reported proteins from NNwg venom, the obtained peptide sequences were aligned from N-C terminus and subjected to BLAST search as mentioned above. Multiple sequence alignment of the top four protein hits along with the peptide sequences obtained through our proteomic characterization were represented through ClustalW.

### 2.14 Data availability

The raw mass spectrometry data and processed files have been deposited to the ProteomeXchange Consortium (http://proteomecentral.proteomexchange.org) [33] via the PRIDE partner repository [34] with the dataset identifier PXD018936.

### 2.15 End point titration ELISA

ELISA was performed as described previously [19]. For this, 10 ng of NNwg venom added on to high bind ELISA plate (Costar) was kept for overnight incubation at 4□. The unbound venom was removed from the wells by inverting the plates and flick drying. Followed by this, any non-reactive sites in the wells were blocked by adding 2.5% BSA in phosphate buffered saline (PBS) and incubated for 2 h at room temperature. Primary antibodies were serially diluted using 2.5% BSA by a factor of three (1:100, 1:300, 1:900, 1:2700, 1:8100, 1:24300, 1:72900, 1:218700, 1:656100, 1:1968300, 1: 5004900) and was incubated for 2 h at room temperature. The unbound antibodies were removed through phosphate buffered saline with 0.1% tween-20 (PBST) wash and this was repeated thrice. Followed by this, secondary antibody (HRP conjugated) at a dilution of 1:32,000 in 2.5 % BSA was added on to each well and kept for incubation at room temperature for 2 h. Excess of secondary antibodies were removed through 0.1% PBST wash as mentioned above. To the wells, 100 μL of HRP substrate solution, TMB/H_2_O_2,_ was added and incubated in the dark for 30 min at room temperature. The end point readings were taken by terminating the reaction by adding 100 μL of 0.5 M sulphuric acid to each well. The optical density values were recorded at 450 nm using a plate reader (Bio-Rad). The EC_50_ values were computed from the obtained OD values through non-linear regression analysis using Prism (version 6.01) software [19].

### 2.16 Western blotting

For immunoblotting experiments, 50 μg of crude NNwg venom resolved on a 15% SDS gel was transferred (25 V for 1 hr) to a PVDF membrane. To minimize non-specific binding, the membrane was blocked using 5% BSA at room temperature for 1 hr. Antivenoms obtained from Virchow, PSAV and VINS were used as primary antibodies (dilution used, 1:500) and was kept for overnight incubation at 4□. The membrane was washed thrice for 10 min using 0.1% PBST solution. To this, secondary antibody (HRP conjugated) at 1:5000 dilution in 5% BSA was added and kept at room temperature for 1 hr. The excess of secondary antibody was then removed through 0.1% PBST wash as mentioned above. Followed by this, ECL substrate was added on to the membrane and signals indicating the venom-antivenom interaction were captured using a plate reader.

### 2.17 Second generation antivenomics

To determine the venom-antivenom interaction, second generation antivenomics protocol was adapted from a previous study [24]. For this, 600 μL of NHS-activated Sepharose 4 fast flow was packed on to a column. The column was washed using 15 column volumes of ice cold 1 mM HCl for removing isopropanol (used for storing column material) and made sure that the pH of the column was maintained between 7.0 and 8.0. After this, the column was pre-equilibrated using coupling solution (0.2 M NaHCO_3_ and 0.5 M NaCl, pH 8.3). To this 30 mg of anti-venom dissolved in the coupling buffer was added and kept for overnight incubation at 4 °C. Subsequently, unbound anti-venom was eluted out and the non-reacted groups were blocked using 0.1M Tris-HCl at room temperature for 4 hr. The column was then alternatively washed using buffers with high (0.1 M Tris-HCl; pH 8.5) and low pH (0.1 M acetate buffer containing 0.5 M NaCl; pH 4.5). After this, the column was equilibrated using 5 column volumes of working buffer solution (1X PBS, pH 7.4). Followed by, 200 μg of venom dissolved in PBS was added. After keeping it for incubation (4 h at room temperature), the unbound venom was washed out using PBS. The bound venom proteins were eluted using 0.1 M glycine-HCl, pH 2.0 and the resultant solution was immediately neutralised using 1M Tris buffer, pH 9.0. The bound and unbound fractions after desalting was subjected to RP-HPLC analysis as mentioned above.

### 2.18 Statistical analysis

Prism (GraphPad software Inc., San Diego, CA; version 5) was used for performing statistical comparisons. The data obtained from triplicate experiments were plotted on to the software and is represented as mean ± standard deviation or as stated, otherwise.

## 3. Results and discussion

### 3.1 Proteomics of *Naja naja* venom from the Western Ghats

The crude NNwg venom components were resolved on a 12% SDS gel. Bands observed after CBB staining indicated that the venom components were distributed across a wider mass range (Fig.1). All the visible bands excised from the samples run under reducing conditions were subjected to various enzymatic digestion protocols followed by tandem mass spectrometric data collection. In parallel, a portion of the crude NNwg venom was fractionated on a reversed-phase column as mentioned in the methodology section. As indicated in Fig. 2a, a total of 12 fractions were collected. Each of these fractions were further separated under reducing conditions on a 12% SDS gel. From the results (Fig. 2b), it is evident that most of the fractions (labelled 1—8) fall around 10 kDa molecular weight region. All these peaks were eluted within 42 min (26% B) of the chromatographic run and rest of the peaks (labelled 9-12) found to contain proteins with molecular weights range between 10-315 kDa.

**Figure 1.**
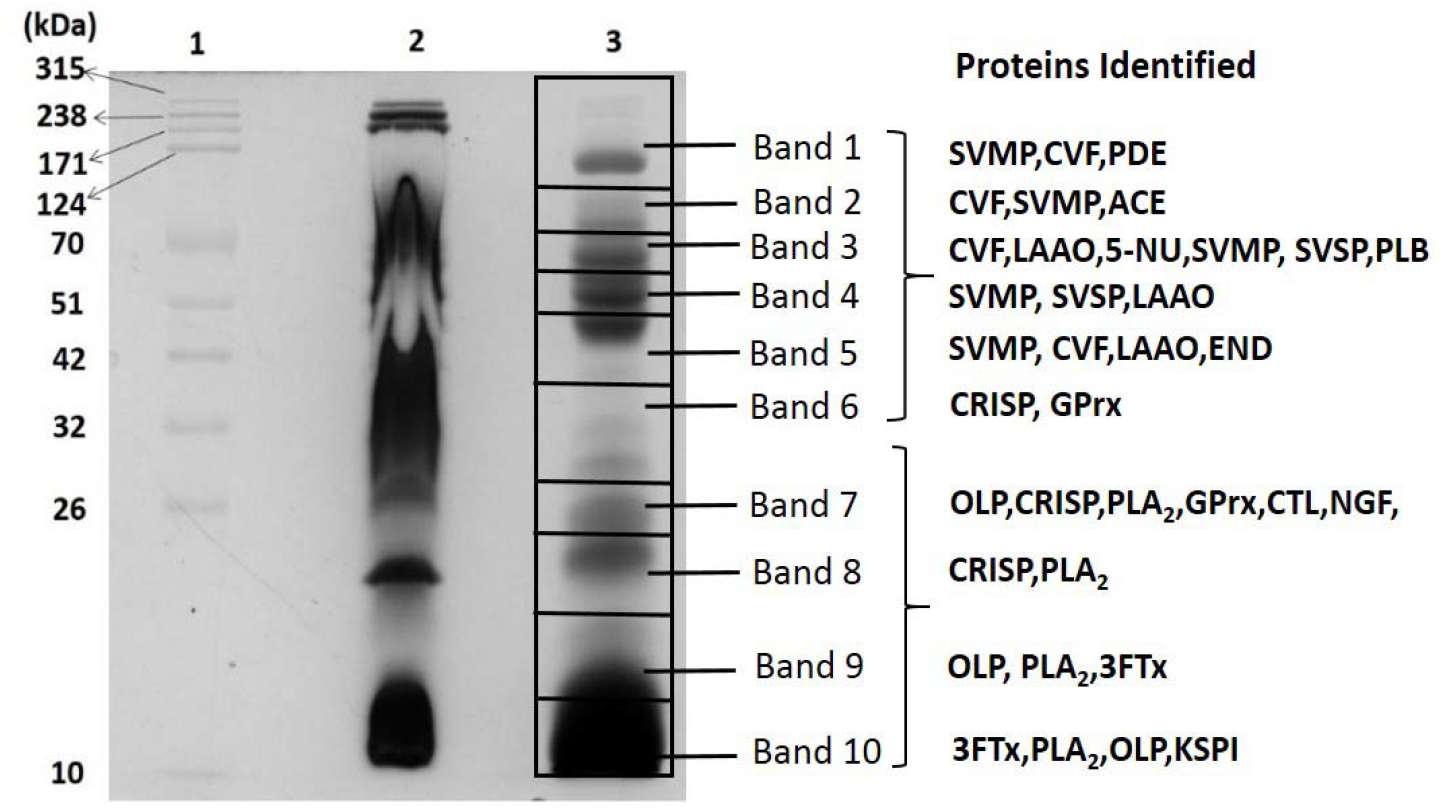
12% SDS-PAGE analysis of crude NNwg venom. Lane 1,molecular weight ladder; Lane 2, Crude NNwg venom resolved without β-ME (non-reducing condition); Lane 3, Crude NNwg venom subjected to β-ME treatment (reducing condition). The bands resolved under reducing conditions were further subjected to various enzymatic digestion (trypsin, chymotrypsin and V8 protease) strategies followed by Q-TOF LC-MS/MS analysis.

**Figure 2a.**
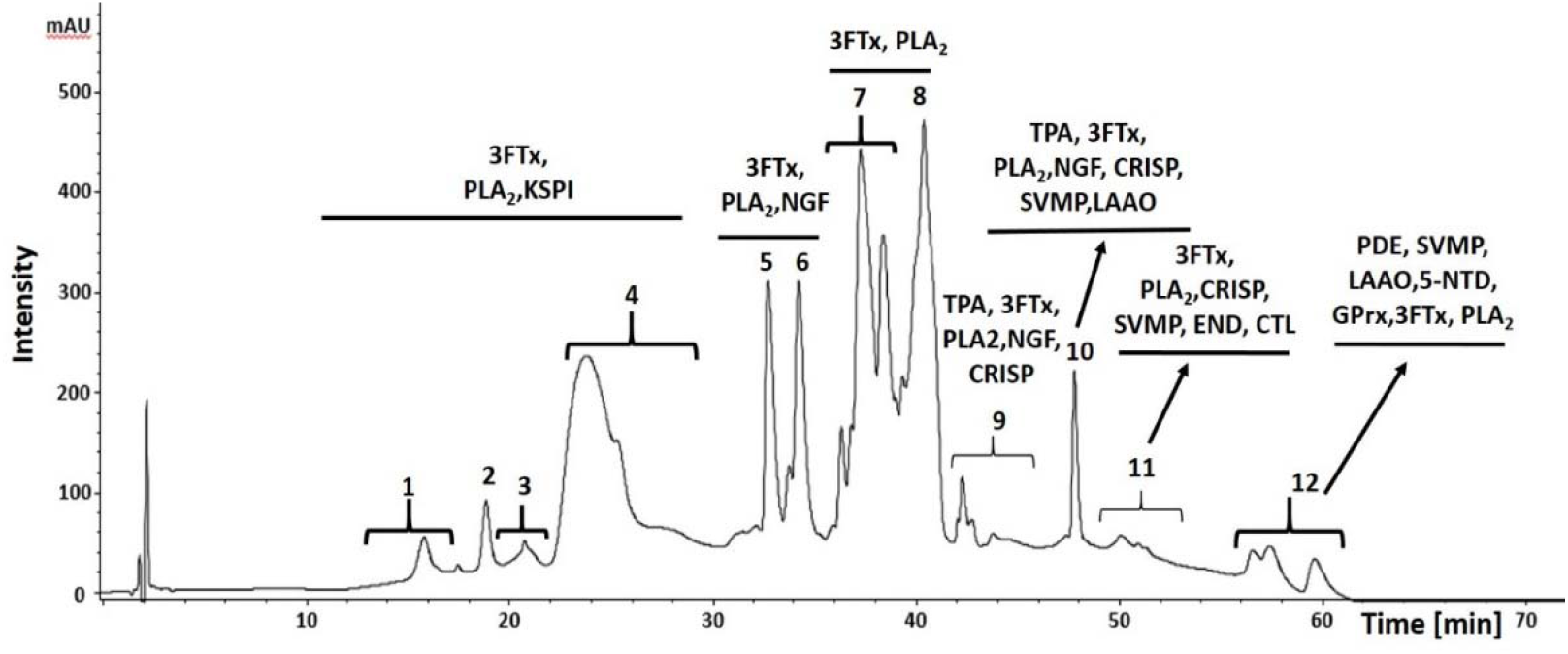
Reversed-phase HPLC profile of crude NNwg venom. Crude *Naja naja* venom (500 μg) was subjected to reversed-phase HPLC separation using an Agilent 1260 HPLC system. X-axis and Y-axis corresponds to the retention time and absorbance observed at 210 nm, respectively.

**Figure 2b.**
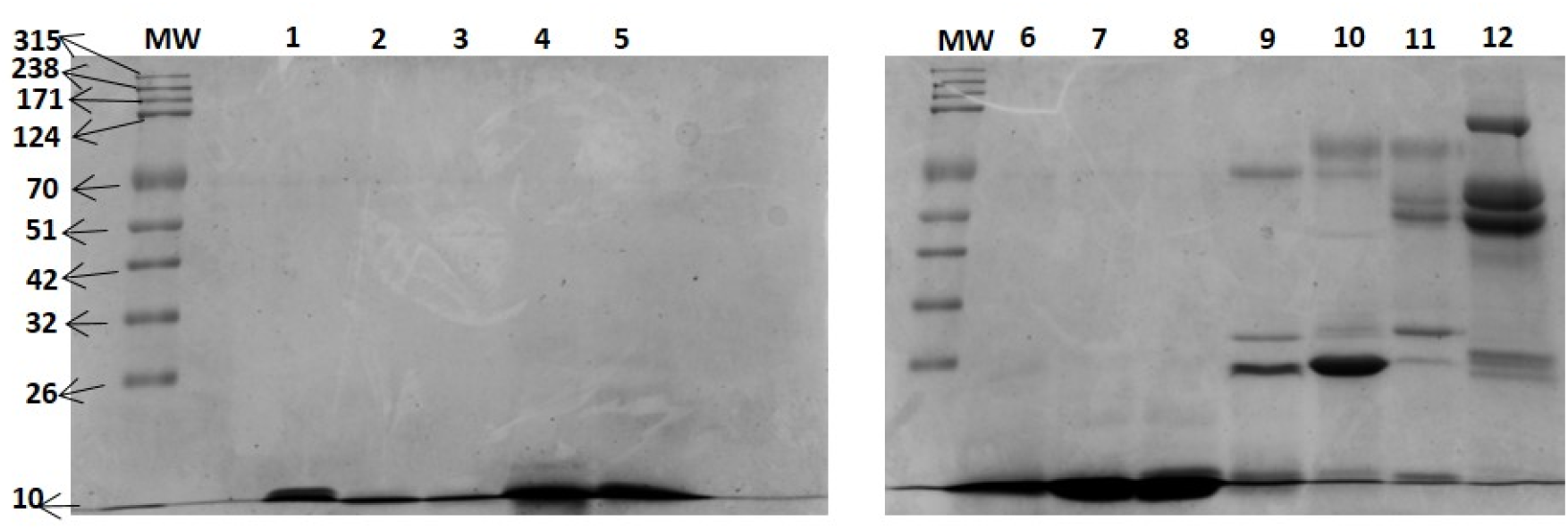
12% SDS-PAGE of RP-HPLC separated fractions of NNwg venom. The 12 fractions collected after HPLC separation were resolved on a 12% SDS gel under reducing conditions. Each of these bands were further diced and subjected to trypsin, chymotrypsin and V8 protease digestion. The resulting peptides were further analysed by using a Q-TOF tandem mass spectrometer. (MW; molecular weight marker)

The proteins and peptides identified through orthogonal separation and multiple enzymatic digestion procedures are shown in Supplementary Table 1. From the Table, it is evident that NNwg venom contains proteins belonging to enzymatic and non-enzymatic families. A total of 115 proteins representing 17 different snake venom protein families were identified. As seen in Fig. 3a, 29% of the identified proteins belongs to 3FTx family followed by PLA_2_ (10%) and SVMP (9%). The identified proteins classified according to their molecular masses indicated that almost 29% of them falls under 10 kDa region (Fig.3b). 3FTxs, the highly abundant proteins commonly found in elapidae snake species are seen in this mass range. Peptides that failed to appear in conventional database searches were subjected to homology-driven proteomic approaches using PEAKS and Novor based *de novo* sequencing followed by MS-BLAST analysis and manual validation. *De novo* analysis of a peptide representing cytotoxin 3a identified from NNwg is shown in Fig. 4. Manual inspection of the doubly charged peptide at m/z 730.4047 indicated that NNwg derived peptide from our study has an arginine instead of lysine (K12R) of the reference peptide. In a similar way, we could identify a total of 39 peptides representing thirteen snake venom protein families through *de novo* analysis. The identified peptides harbouring one or more amino acid mutations are shown in Table 1. All the proteins identified from NNwg venom were then grouped under different snake venom protein families depending on the extent to which they mediate various biological functions and are outlined in detail below [7].

**Figure 3.**
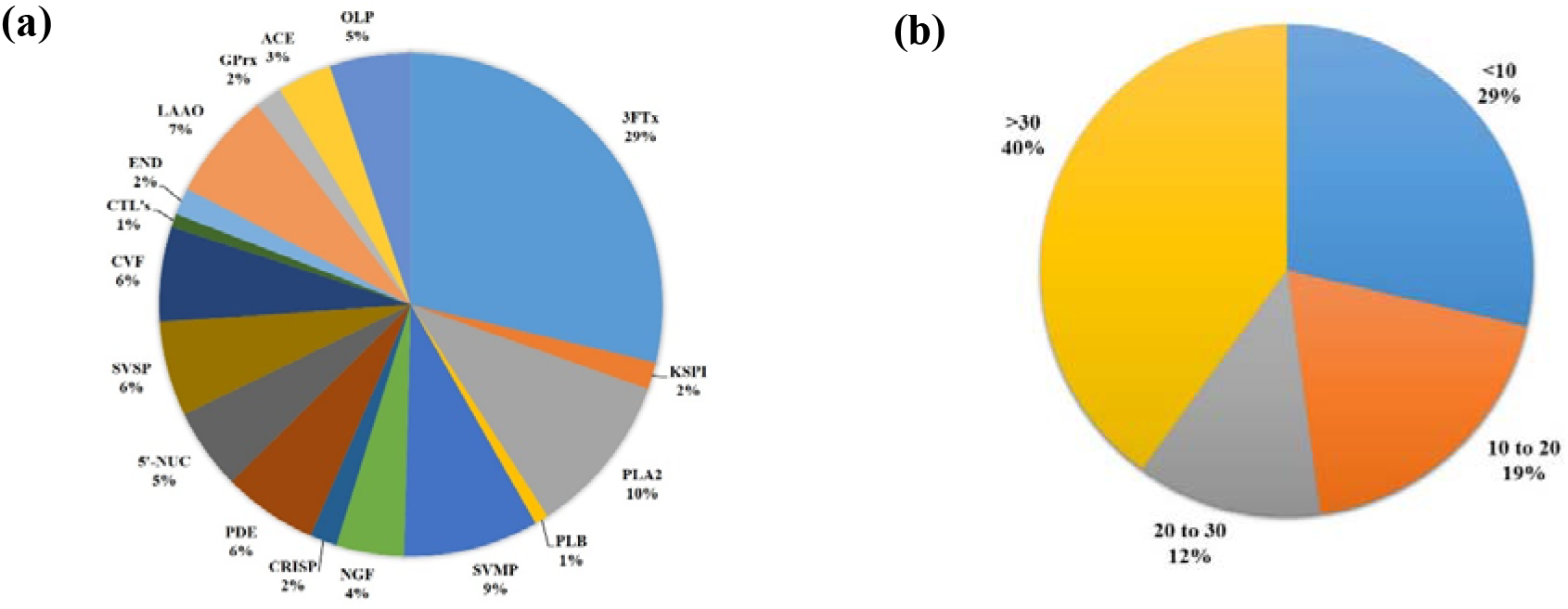
Relative abundance of protein families identified from NNwg venom. The relative abundance of NNwg venom proteins identified (a) after different separation and digestion protocols and (b) according to molecular weight. [Relative abundance = (Number of proteins identified representing a snake venom protein family/ Total number of proteins identified) X 100]. (PLA_2_-Phospholipase A_2_, PLB-Phospholipase B, SVMP- Snake Venom Metalloprotease, NGF- Nerve Growth Factor, CRISP- Cysteine Rich Secretory Protein, PDE- Phosphodiesterase, 5’-NUC- 5 Nucleotidase, SVSP- Snake Venom Serine Protease, 3FTx- 3 Finger Toxin, CVF- Cobra Venom Factor, LAAO- L-Aminoacid Oxidase, GPrx- Glutathione peroxidase, ACE- Acetylcholinesterase, OLP- Ohanin/ Vespryn family proteins, END- Endonuclease, CTL-C Type Lectin, GPrx-Glutathione Peroxidase)

**Figure 4.**
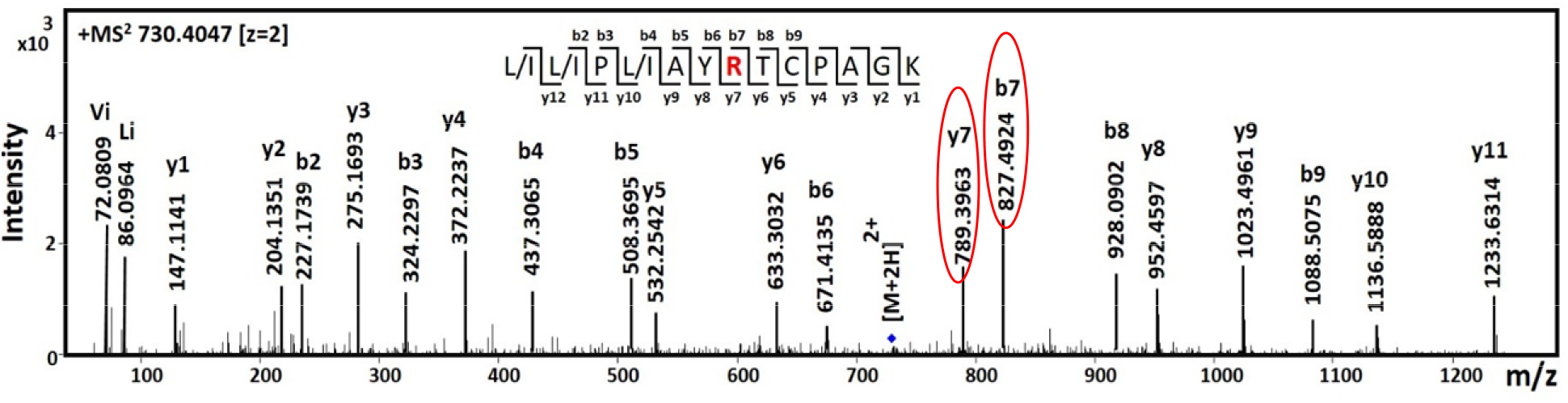
*De novo* analysis of a peptide representing NNwg venom protein. Cobra venom derived peptide (shown in inset) has sequence similarity to Cytotoxin 3a (P86539; *Naja naja*) harbouring one amino acid mutation. The m/z values and corresponding ion intensities are labelled in the x- and y-axis, respectively. MS/MS spectrum representing b and y ions are labelled along with the immonium ions (depicted by single letter amino acid code along with ‘i’ as subscript). The b7 and y7 ions representing the mutated amino acid arginine, R, are circled in the spectrum.

**Table 1:**
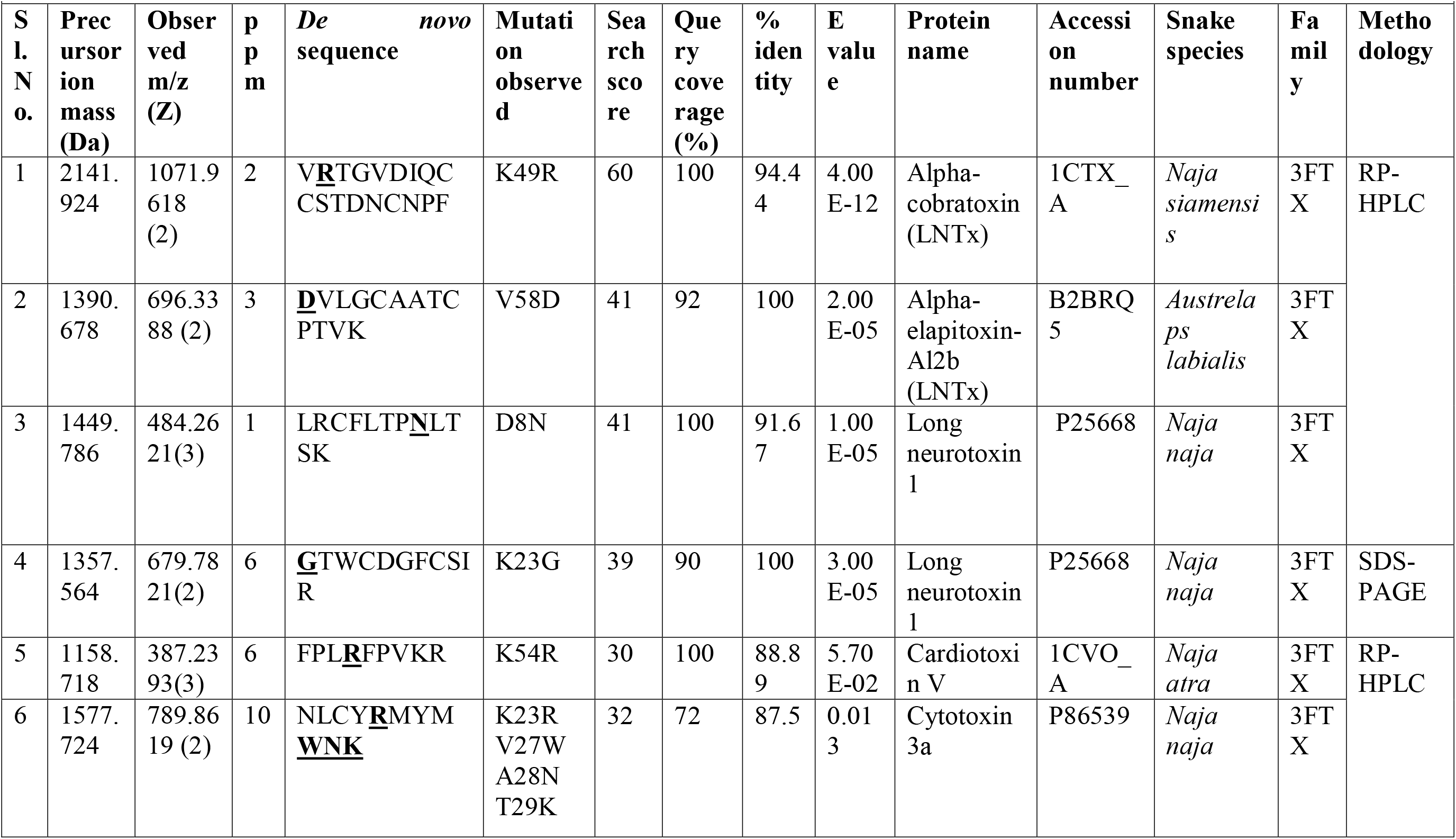
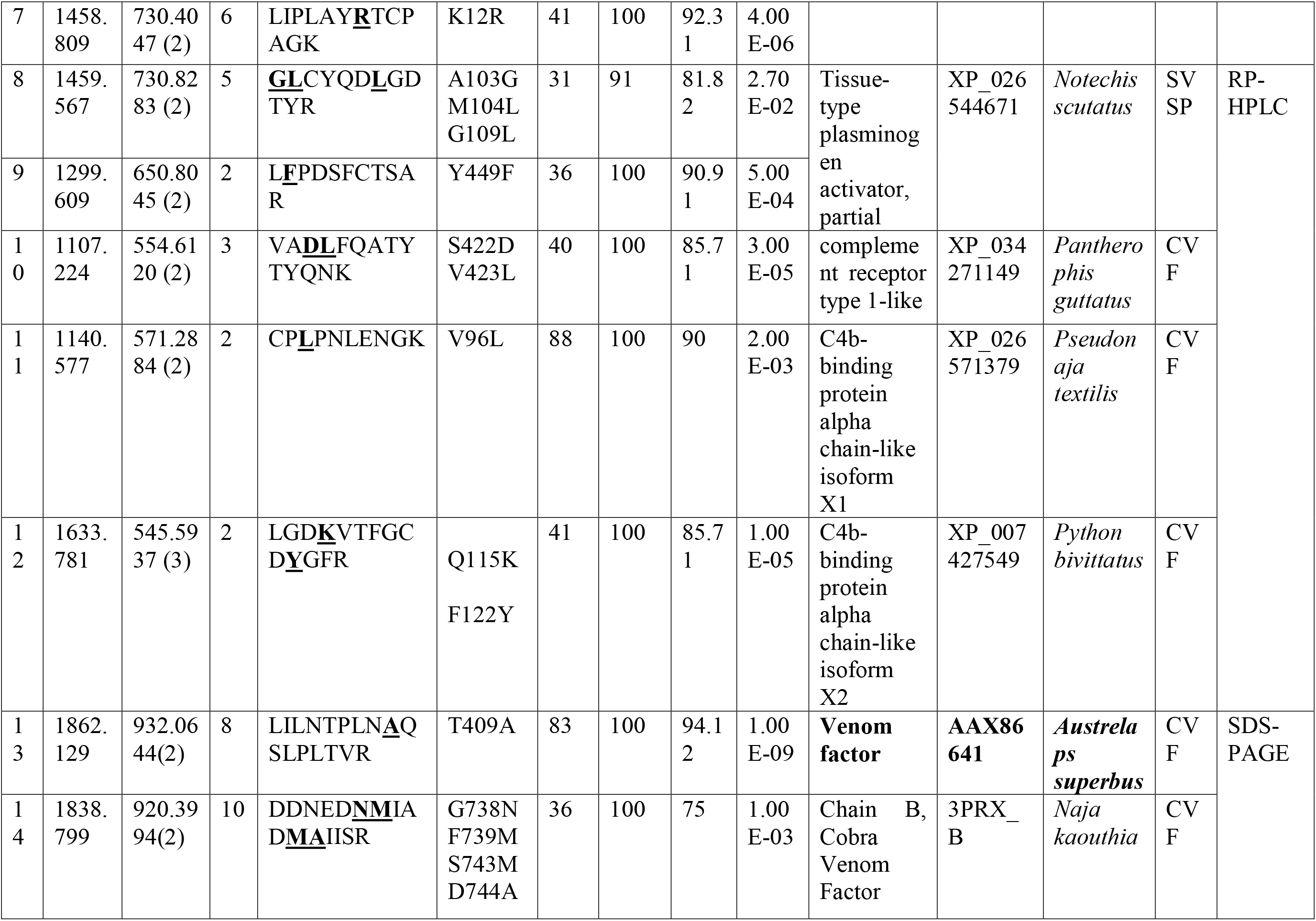
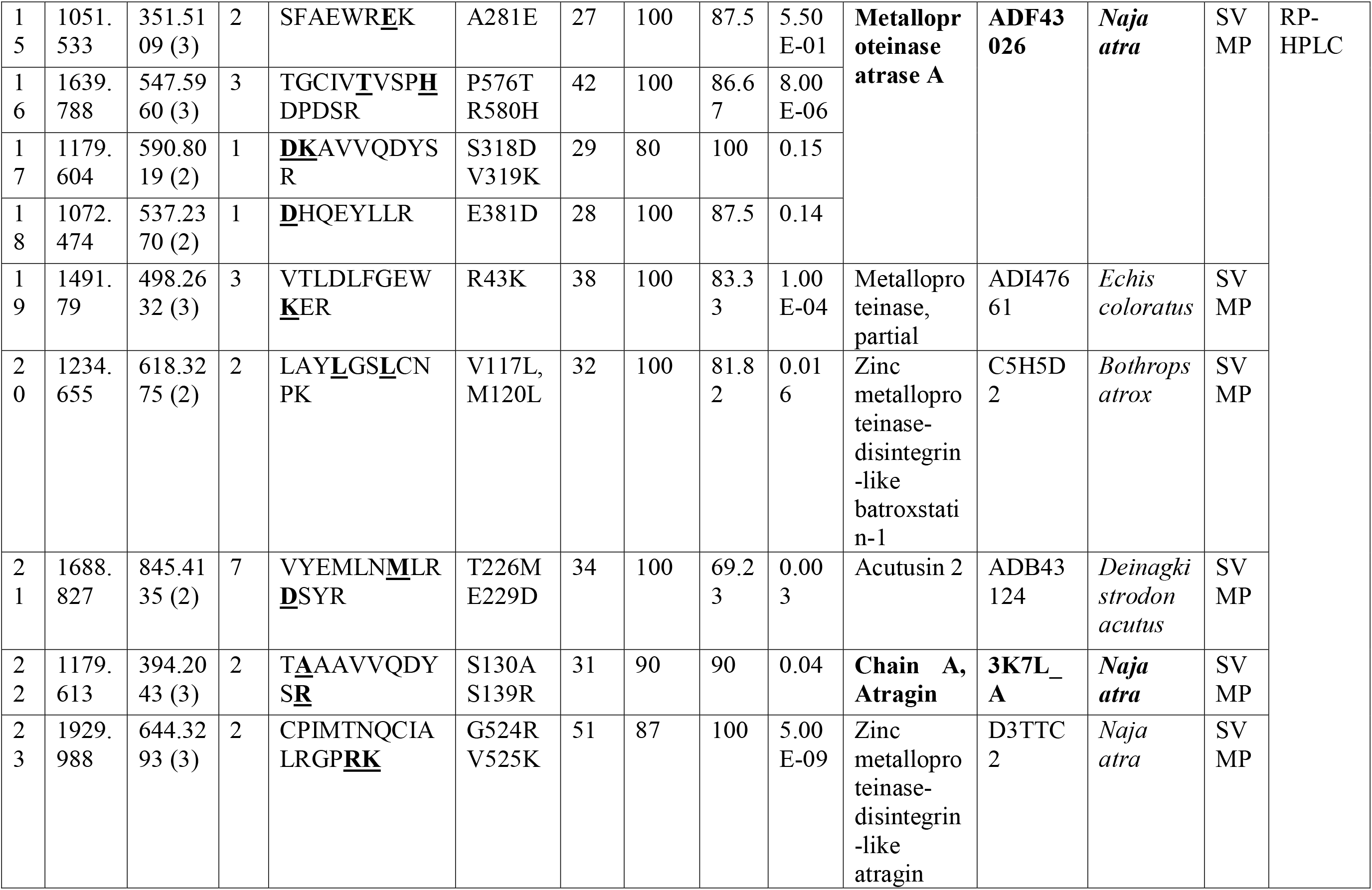
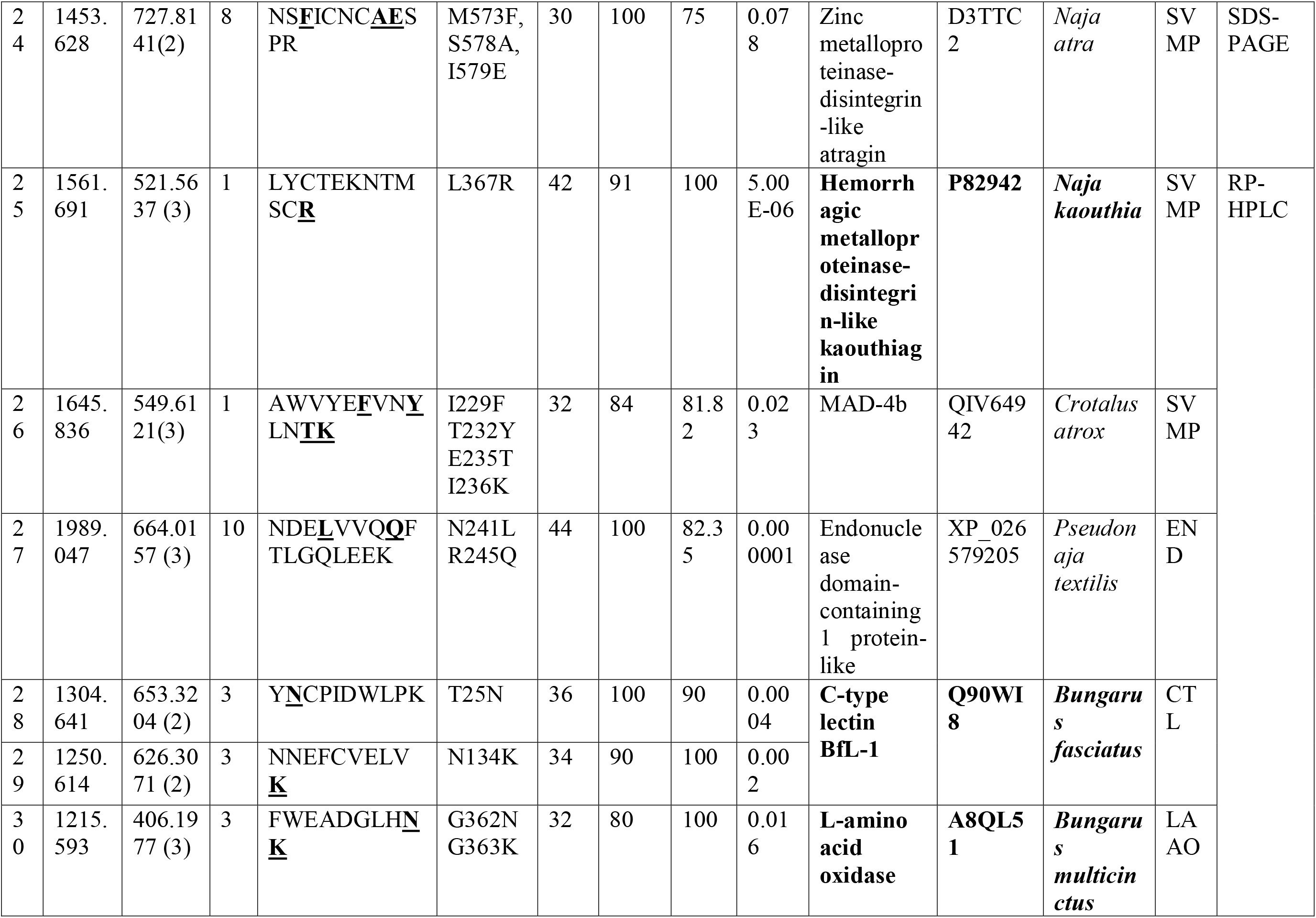
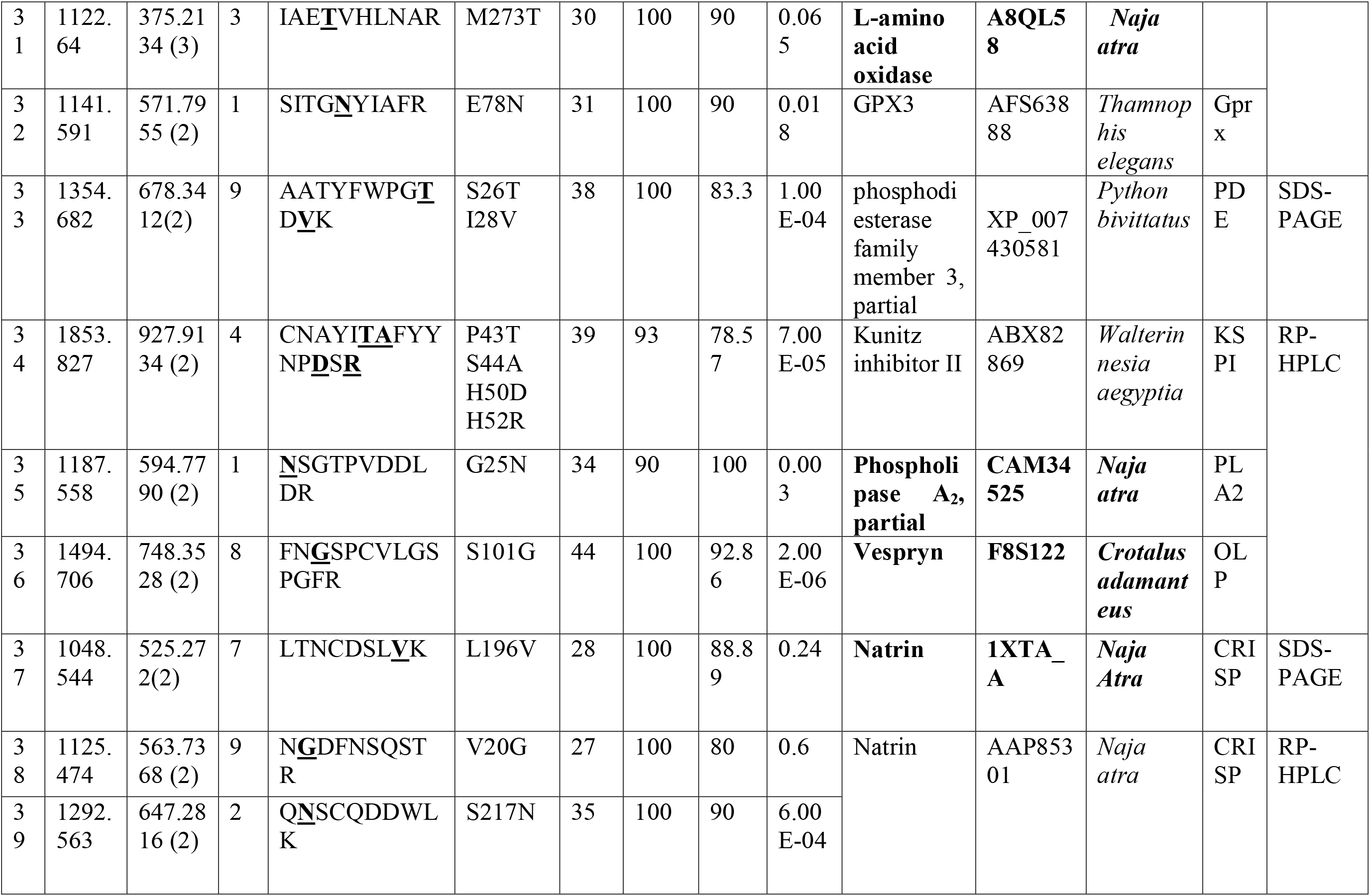
Peptides representing various snake venom proteins identified through de novo analysis. *De novo* sequencing was performed using PEAKS and Novor softwares followed by MS-BLAST. The mutated amino acids observed are highlighted in bold and underlined. Proteins marked in bold were identified through both conventional proteomics and *de novo* methods.

**Table 2:** Comparison of protein families identified in cobra venom collected from different geographical location of India.

#### 3.1.1 Dominant protein families identified from NNwg venom

The major enzymatic and non-enzymatic protein families identified from NNwg venom includes three-finger toxins, phospholipase A_2_ (PLA_2_s), snake venom metalloproteases (SVMP) and snake venom serine proteases (SVSP).

Snake species belonging to the Elapidae family are abundant with proteins representing three-finger toxins [7]. In agreement with these, our analysis resulted in the identification of 33 proteins representing 3FTxs (Supplementary Table 1) and contributed to 29% of the total proteins identified (Fig. 3a). Three finger toxins present in cobra venom belong to low molecular mass proteins (comprise of 57 to 82 amino acids), that include neurotoxins, cytotoxins/cardiotoxins and muscarinic toxins [35, 36]. Neurotoxins and muscarinic toxins bind to muscle nicotinic acetylcholine receptors (nAChRs) and muscarinic acetylcholine receptors, respectively [35, 37, 38]. As seen in Supplementary Table 1, almost all subclasses of 3FTxs including long and short neurotoxins, weak toxins, muscarinic toxin-like protein, cytotoxins/cardiotoxins were identified from NNwg venom. Most of the NNwg venom derived 3FTxs proteins showed sequence similarity with the proteins reported from various Naja species. Though Mascot searches enabled the determination of peptides representing long neurotoxin proteins belonging to 3FTx family, our *de novo* approaches helped in the identification of four additional peptides from the same category (Table 1). The peptide sequence with m/z, 1071.961, was observed to contain one amino acid substitution at the 49^th^ position (K49R) when compared to the sequence present in the database (Suppl.Fig.1a (i)).

The peptide, DVLGCAATCPTVK, representing an Alpha-elapitoxin was also obtained. Comparing to the *Austrelaps labialis* counterpart, doubly charged NNwg peptide with m/z, 696.3388, was having V58D mutation (Suppl.Fig.1a (ii)). Similarly, through tandem mass spectrometric analysis and homology searches, the peptide at m/z, 484.2621 (z=3) was having asparagine in the position 8 instead of aspartate (Suppl.Fig.1a (iii)) and the doubly charged peptide at m/z, 679.7821 was contained to have a K23G substitution in the reference sequence (Suppl.Fig.1a (iv)).

Similarly, as seen in Supplementary Table1, our analysis resulted in the identification of 18 proteins representing different cytotoxins such as cytotoxin-1, 2, 7, 8, 9, 13 and 14. Additionally, three peptides representing cytotoxins were derived through *de novo* approaches (Table 1). The peptide at m/z, 789.8619, was found to have four amino acid mutations (K23R, V27W, A28N, and T29K) (Suppl.Fig.1a (vi)). Similarly the peptides, FPLRFPVKR and LIPLAYRTCPAGK, were having amino acid mutation at the 54^th^ (K54R) and 12^th^ (K12R) positions, respectively (Suppl.Fig.1a (v, vii)).

PLA_2_s, the collection of proteins belonging to enzymatic venom protein family, are mainly involved in releasing lysophosphoglycerides and fatty acids by the catalytic cleavage of phosphoglycerides. They mediate multiple pharmacological effects such as platelet aggregation, anticoagulation, neurotoxic and cardiotoxic activities [39]. Most importantly, the myotoxic, neurotoxic and cytotoxic potential of several purified PLA_2_s from the Indian cobra venom has been studied in detail [40–42]. As seen in Supplementary Table 1, twelve proteins representing several PLA_2_s are identified through the present study. Additionally, a doubly charged tryptic peptide, NSGTPVDDLDR, with m/z 594.779, representing a PLA_2_ protein was determined through *de novo* analysis (Table 1). On manual inspection, it was found that NNwg derived peptide has asparagine instead of glycine in the 25^th^ position (Suppl.Fig.1b). Altogether, the identified PLA_2_s constitutes to 10% of the total proteome of NNwg venom (Fig. 3a).

Snake venom proteases, another major venom proteins belonging to the enzymatic family, are known to cleave fibrinogen and affect haemostasis through diverse anti- and pro-coagulant mechanisms [43]. Unlike viper venoms, both serine and metalloproteases are less abundant in elapidae species (12-19%) [7]. In accordance with this, our analysis indicated that SVMPs (9%) and SVSPs (6%) contribute only 15% of the total NNwg venom proteome identified (Fig.3a). SVMPs are zinc-dependent high molecular mass proteins with multidomain organization, divided into four main classes, PI, PII, PIII and PIV. They possess fibrinolytic, haemorrhagic and platelet aggregation inhibitory activities [44]. For example, kaouthiagin, a metalloprotease that cleaves and binds to von Willebrand factor (vWF) was isolated and characterized from *Naja kaouthia* venom. Cleavage of vWF results in collagen mediated platelet aggregation inhibition [45]. As seen in Supplementary Table 1, several peptides representing two kaouthiagin proteins were identified from the present study. Also, it is reported that NN-PF3, a non-toxic anticoagulant SVMP protein inhibits collagen induced plated aggregation through binding to α2β1 integrin receptors [46]. Like cobra venom SVMPs (NN-PF3 and kaouthiagin), several other PIII-SVMP_S_ from different snake species were characterized for their platelet-aggregation inhibitory activities [47–49]. As seen in Supplementary Table 1, most of the SVMPs identified through our study belong to the PIII class. Though peptides representing various metalloproteases were derived through Mascot-enabled database search, twelve more additional peptides representing the same protein family were obtained through homology-assisted *de novo* searches (Table 1). One of the identified peptides, LYCTEKNTMSCR (L367R), was representing a kaouthiagin protein present in the database (Suppl.Fig.1c (i)). Similarly, the peptide, TAAAVVQDYSR, has Ala-130 and Arg-139 instead of Ser-130 and Ser-139 when compared to the *N.atra* counterpart (Suppl. Fig.1c (ii)) and the peptide CPIMTNQCIALRGPRK (m/z, 644.3293), was found to have two amino acid substitutions at the C-terminus (G524R, V525K) compared to the original sequence present in the database (Suppl.Fig.1c (iii)). Another metalloproteinase peptide sequence, VTLDLFGEWKER, was shown to have an amino acid mutation, R43K (Suppl.Fig.1c (iv)). Apart from this, eight more peptides representing multiple SVMPs were identified through *de novo* approach. The details of all the *de novo* derived peptides are provided in Table 1 and Suppl.Fig.1c (v-xii).

SVSPs, on the other hand, are mainly composed of thrombin-like and fibrinolytic proteins. Thrombin-like proteins are primarily used as defibrinogenating agents due to its ability to remove fibrin (formed during clotting) from the blood circulation [50]. As seen in Supplementary Table 1, all the NNwg venom derived thrombin-like proteins showed sequence similarity to peptides reported from *Naja naja* and *Gloydius ussuriens* species. In addition to this, tissue-type plasminogen activator proteins were also present in NNwg venom. Since plasmin is involved in degrading clots formed by fibrin, tissue type plasminogen activators can be used as thrombolytic agents for treating deep vein thrombosis [51]. Apart from the proteins reported in Supplementary Table 1, our *de novo* analysis resulted in the identification of two additional peptides, GLCYQDLGDTYR, LFPDSFCTSAR, harbouring various amino acid mutations representing SVSP proteins (Table 1, Suppl.Fig.1d). In the former peptide sequence, three mutations were identified (A103G, M104L, G109L) (Suppl.Fig.1d (i)). Whereas in the latter sequence, tyrosine has been replaced by a phenylalanine (Y449F) (Suppl.Fig.1d (ii)).

#### 3.1.2 Secondary protein families identified from NNwg venom

Proteins belonging to L-amino acid oxidase (LAAO), Kunitz-type serine protease inhibitors (KSPI), C-type lectins (CTLs) and Cysteine-rich secretory proteins (CRISP) falls under this category [7].

LAAOs are flavoenzymes with flavin adenine dinucleotide as cofactors. Flavins are accountable for the yellow colour of snake venom and they mediate cell toxicity by inducing oxidative stress through hydrogen peroxide release [52]. They are also involved in arbitrating platelet aggregation activation or inhibition, anticoagulation, myonecrotic and haemorrhagic activities [52, 53]. From the LC-MS characterization, it is evident that 7% of the total proteins identified from NNwg venom represents LAAO family (Fig.3a). All the NNwg venom derived LAAO peptides showed sequence similarity to the peptides reported from *Bungarus multicinctus*, *Oxyuranus scutellatus* and several other Naja species (Supplementary Table 1). On the other hand, *de novo* sequencing yielded two more additional peptides harbouring different amino acid mutations. As seen in Table 1, the peptides, FWEADGLHNK and IAETVHLNAR, representing an LAAO, showed sequence similarity with *Bungarus multicinctus* and *Naja atra* proteins, respectively. Manual inspection of the MS/MS data of FWEADGLHNK confirmed that the NNwg venom derived peptide has asparagine and lysine at the 360^th^ and 361^st^ positions, respectively (G362N and G363K) (Suppl.Fig.1e (i)). Similarly, the triply charged peptide (IAETVHLNAR) at m/z, 375.2134, was found to contain threonine instead on methionine (M273T) (Suppl.Fig.1e (ii)).

Kunitz-type serine protease inhibitors are known to act on several serine proteases and MMP-2s [54]. They inhibit activity of coagulation factors including thrombin, protease C, factors Xa and XIIa and also found to regulate blood pressure [55]. Considerable number of KSPIs are identified through proteomics analysis of *Naja naja* venom from different parts of India [5, 8–12]. Our study also resulted in the identification of eight different peptides representing two KSPI proteins (Supplementary Table 1). The *de novo* sequencing methods on the other hand has resulted in the identification of an additional peptide representing a kunitz-type inhibitor protein reported from *Walterinnesia aegyptia* venom (Table 1 and Suppl.Fig.1f). On manual inspection, the doubly charged peptide with m/z, 927.9134, was contained to have four amino acid mutations (P43T, S44A, H50D and H52R).

Snaclecs, the C-type lectins found in snake venom are known to activate and inhibit platelet aggregation mechanisms [56]. Though venom proteome of Indian cobra has been studied from various geographical locations, only one study has revealed the presence of a CTL that showed sequence similarity to a C-type lectin BfL-1 from *Bungarus fasciatus* [12]. This study also identified peptides representing the same protein and are included in Supplementary Table 1. Through *de novo* approach, two peptides representing the above mentioned protein (Table 1) were identified. The peptides, YNCPLDWLPK and NNEFCVELVK, were contained to have one amino acid mutation (T25N, N134K) each (Suppl.Fig.1g (i & ii)).

Our proteomic analysis resulted in the identification of proteins representing CRISP family too. The identified peptides showed sequence similarity to CRISP proteins reported from two Naja species (Supplementary Table 1). Apart from this, three doubly charged peptides, LTNCDSLVK, NGDFNSQSTR and QNSCQDDWLK exemplifying CRISP proteins harbouring single amino acid mutations were identified from NNwg venom through *de novo* approaches (Table 1). For the peptide with m/z 525.272 (z=2), leucine in the 196^th^ position has been substituted by valine (Suppl.Fig.1h (i)). Whereas the other two doubly charged NNwg venom derived peptides with m/z, 563.7368 and 647.2816 have V20G and S217N mutations, respectively (Suppl.Fig.1h (ii &iii)).

#### 3.1.3 Minor protein families identified from NNwg venom

The minor proteins families present in snake venom includes 5’-nucleotidase (5-NU), Phosphodiesterase (PDE), Pospholipase B (PLB), Acetylcholinesterase (ACE), Nerve growth factor (NGF) and Vespryn/ohanin (OLP) [7].

The nuclease activity seen in snake venom is due to the presence of phosphodiesterases and 5-nucleotidases [57, 58]. Several of these enzymes with potent nuclease activity was isolated from various Naja species. In our study too, peptides representing seven PDEs and six 5-NUC have been identified (Supplementary Table 1). In addition, a peptide representing a PDE harbouring two amino acid mutations were also identified through the *de novo* analysis (Table 1). The peptide, AATYFWPGTDVK was contained to have S26T and I28V mutations with respect to the reference protein present in the database (Suppl.Fig.1I).

Since ohanin like proteins are present only in smaller amounts, they are not isolated or characterized in detail from snake venoms. Nevertheless, ohanin identified from king cobra has shown to induce hyperalgesia and hypolocomotion in mice [59]. In addition to six proteins identified through conventional approaches (Supplementary Table 1), our *de novo* analysis yielded a doubly charged tryptic peptide, FNGSPCVLGSPGFR, having a single amino acid mutation (S101G) representing ohanin protein family (Table 1 & Suppl.Fig.1j). Further, the peptides identified through our analysis showed sequence similarity to cobra venom factor proteins reported from *Ophiophagus hanna*, *Naja kaouthia*, *Pseudonaja textilis* and *Bungarus fasciatus* (Supplementary Table 1). Apart from this, *de novo* characterization resulted in the identification of five additional peptides with various amino acid mutations representing several CVF proteins. As seen in Suppl.Fig.1k (i &ii) and Table 1, two of the *de novo* derived peptides, CPLPNLENGK and LILNTPLNAQSLPLTVR, were having one amino acid mutation. The doubly charged peptides, at m/z 571.2884 and 932.0644, were found to contain V96L and T409A mutations, respectively (Suppl.Fig.1k (i &ii)). On a similar way, three more peptides with multiple amino acid mutations representing CVF proteins were obtained through the study (Suppl.Fig.1k (iii-v)).

Though Mascot searches enabled the determination of peptides representing proteins belonging to endonuclease family (Supplementary Table 1), our *de novo* study helped in the identification of an additional peptide with two amino acid mutations (Table 1). The triply charged peptide sequence with m/z, 664.0157, was observed to contain two amino acid substitutions at the 241^st^ (N241L) and 245^th^ (R245Q) positions when compared to the sequence present in the database (Suppl.Fig.1l).

Peptides representing glutathione peroxidase proteins were also identified from this study (Supplementary Table 1). In addition, a doubly charged peptide, SITGNYIAFR, with m/z 571.7955, representing the same family was identified through *de novo* analysis (Table 1). The obtained peptide showed sequence similarity to a GPX3 protein reported from *Thamnophis elegans* and through manual analysis it was confirmed that Glu-78 in the original sequence has been replaced by an asparagine (Suppl.Fig.1m).

Apart from the above-mentioned protein families, peptides representing acetylcholinesterases, phospholipase B and nerve growth factor were also identified from NNwg venom (Supplementary Table 1).

### 3.2 Newly identified protein families from the Indian Cobra venom

Though a total of 17 different snake venom protein families were identified, it is interesting to note that glutathione peroxidase and endonucleases were reported for the first time in Indian cobra venom. The peptide information representing these families of proteins are included in Supplementary Table 1 and Table 1. For confirming the authenticity of the derived peptides, sequences were aligned from N-C terminus and subjected to pBLAST using the non-redundant database from NCBI. The sequences obtained were further subjected to multiple sequence alignment using ClustalW. Figures 5a and b show the sequence similarity of the obtained sequences with respect to various snake species. As seen in Supplementary Table 1, the peptides representing GPrx 3 identified from NNwg venom showed 77% sequence similarity to a glutathione peroxidase reported in *Ophiophagus hannah*. Similarly, the endonuclease identified in this study shown to have 53% sequence similarity with an *O.hannah* protein (accession no: ETE56723).

**Figure 5.**
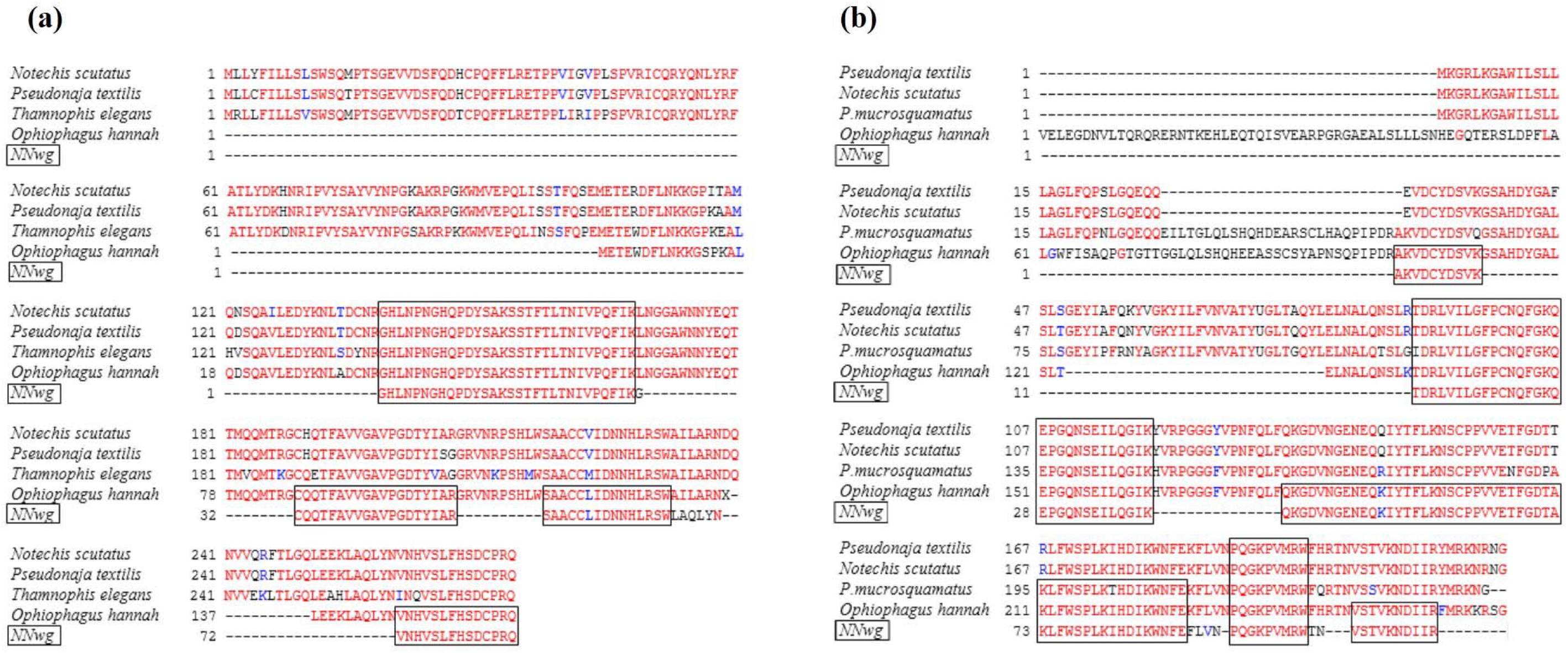
BLASTp and ClustalW analysis of endonuclease and glutathione peroxidase proteins identified from NNwg venom. (a) Endonuclease and (b) Glutathione peroxidase peptide sequences identified from NNwg venom after Mascot search were subjected to BLASTp analysis followed by multiple sequence alignment using known sequences from various snake species. Those highlighted in-boxes are the shared amino acid sequences between NNwg and other snake species.

### 3.3. Importance of *de novo* sequencing strategies in elucidating NNwg venom proteome

Previous studies from our laboratory have shown the importance of *de novo* sequencing workflows in elucidating snake venom proteome profiles [19, 20]. It was observed that the use of complementary identification strategies improves the confidence of protein identification. In agreement with this, our analysis on *Naja naja* venom proteome through *de novo* approaches resulted in improving sequence information of proteins that were determined through conventional proteomic approaches. For example, though a SVMP protein, atrase-A was identified through conventional approaches (Supplementary Table 1, SI No-55), four additional peptides representing the same protein (accession no-ADF43026) were determined through *de novo* analysis. Likewise, a total of 14 peptides (Table1; accession numbers marked in bold) representing ten different snake venom proteins complements the conventional proteome data. Though venom gland transcriptomics has opened up the possibilities in exploring snake venom proteome profiles, due to its limitations, details on protein isoforms might be missed [60, 61]. For example, venom proteomics of *Sistrurus catenatus edwardsii* studied using different ionisation mechanisms using MALDI and ESI mass spectrometry enabled the identification of various protein isoforms that were not detected through the transcriptomics study [60]. Similarly, in the current study, several protein isoforms were derived through homology-driven approaches. As seen in Table 1, a peptide representing 3FTx (accession-1CTX_A) protein was contained to have one amino acid mutation (K49R) in the *de novo* derived sequence and the original sequence without the mentioned mutation, presumed to belong to an isoform, was observed through Mascot analysis. Similarly, a total of five peptides representing various protein isoforms were derived through our analysis (Table 1; SI No-1, 3, 4, 5 &14). This is indicative of the fact that homology approaches helps in the determination of protein isoforms too. In addition to this, 20 different *de novo* derived peptides representing various snake venom protein families were identified through *de novo* analysis (Table 1). These finding highlights the importance of following multiple data analysis approaches in elucidating the venom proteome of snake species whose protein sequence information is limited in the public repositories.

### 3.4 Commercial antivenoms showed varied immunological cross-reactivity profiles towards NNwg venom

From a clinical perspective, detailed information on the toxin components present in venom helps in streamlining antivenom production protocols. As recommended by World Health Organisation, various in vitro preclinical studies including ELISA, western blotting and immunoaffinity chromatography are widely used in determining the specificity and reactivity of commercially available antivenoms towards various snake venoms. In India, polyvalent antivenoms are manufactured by using pooled venom from ‘Big Four’ snake species that include *Naja naja, Bungarus caeruleus, Daboia russelii and Echis carinatus*. Still, commercial antivenoms from the major Indian manufacturers’ present varied immunological reactivity towards cobra venom collected from different geographical locations in India [8, 10, 11]. In agreeing with the above observation, our results obtained through end-point titration ELISA indicate that the immuno-cross-reactivity of VINS antivenom (EC_50_: 2.48 μg/mL) is better compared to Virchow (6.03 μg/mL) and PSAV antivenoms (6.04 μg/mL) (Fig. 6a). All the tested Indian antivenoms showed a dose-dependent increase in reactivity towards NNwg venom components. However, as seen in Fig. 6a, the binding potential seems to be varied across the tested antivenoms. Though the end point titration values of all the antivenoms were same (1:72,900), the strength of binding seen at the same dilution was higher in Virchow antivenom. At the same time, EC_50_ values computed from the graph suggest that VINS (2.48 μg/mL) showed better reactivity towards NNwg venom compared to that of PSAV (6.04 μg/mL) and Virchow (6.03 μg/mL) antivenoms.

**Figure 6.**
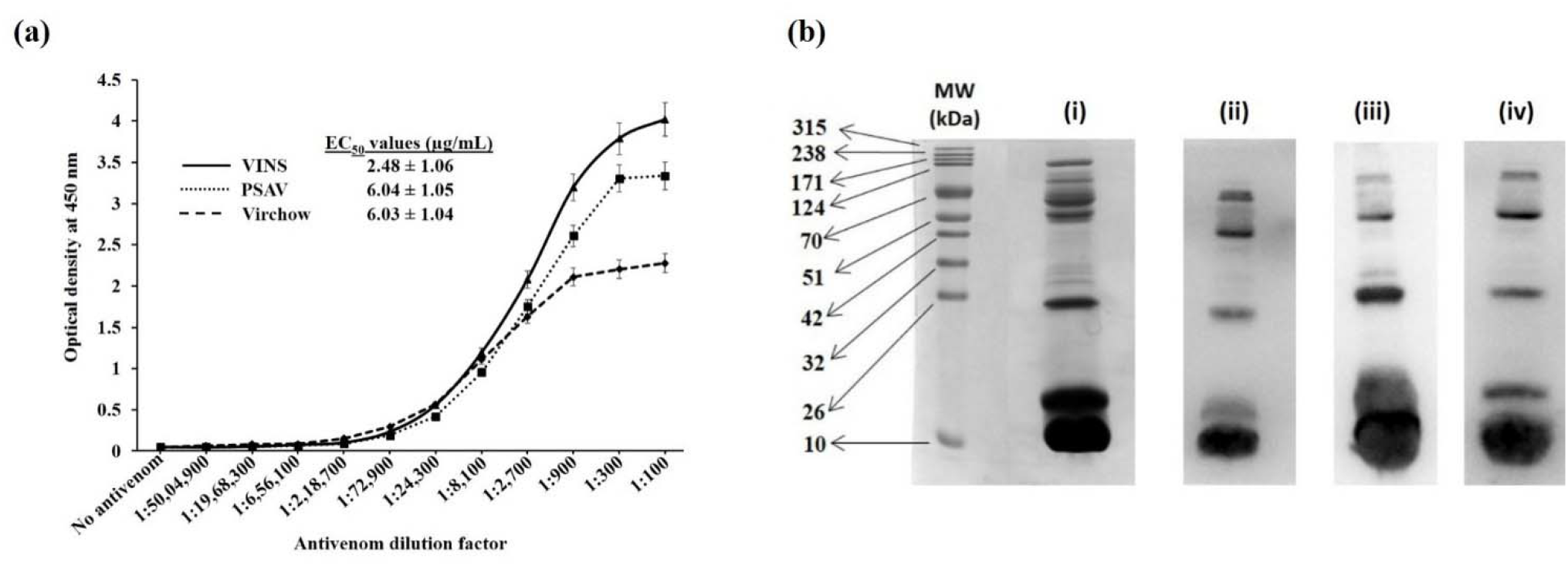
Assessment of the immunological cross-reactivity of different polyvalent antivenoms towards NNwg venom. (a) Titration curves obtained through ELISA using various dilutions of VINS, PSAV, Virchow antivenoms. Results are represented as mean ± SD of triplicates. The EC_50_ values for different antivenoms obtained from this graph are shown in insets. (b) (i) 15% SDS-PAGE gel electrophoretic profile of crude NNwg venom under reducing conditions. Immunoblot of the SDS-PAGE with (ii) VINS (iii) PSAV and (iv) Virchow polyvalent antivenoms.

Similarly, the specificity of antivenom binding towards NNwg venom was assessed through immunoblot analysis. As seen in Fig. 6b (i), the resolved proteins were transferred on to a PVDF membrane and immunoblotting was performed using various polyvalent antivenoms as primary antibodies. From the results (Fig.6b (ii-iv)), it can be inferred that the binding specificities are varied across different antivenom preparations. Hence, there are notable differences in detecting the antigenic epitopes of NNwg venom by various antivenoms. It was evident that low molecular mass proteins were abundant in NNwg venom (Fig. 6b (i)). Irrespective of this, compared to low molecular mass proteins, all the antivenoms recognized high molecular mass proteins with better specificity. Previous reports indicate that high molecular mass proteins are less toxic but highly immunogenic compared to highly toxic low molecular mass proteins that are poorly immunogenic [62], and the same was observed from our findings. Hence, our observations are in agreement with the previous data that reveal the failures associated with Indian polyvalent antivenoms especially in recognizing small molecular mass proteins falling under 20 kDa region [8, 10]. The differences in the antivenom neutralisation efficiencies might also be contributed by the differences in the venom antigens that were used for immunizing horses while generating polyvalent antivenoms [63].

### 3.5 VINS antivenom was less efficient in detecting many low molecular mass proteins falling under 20 kDa

Though ELISA and western blotting can provide an overall immunoreactive profile of venom and antivenom interaction, they are not successful enough to distinguish the antivenom binding and non-binding protein components. To verify this, we have followed a second generation antivenomics approach combining immunoaffinity chromatography and LC-MS based proteomic characterization [24]. Since the immuno-cross-reactivity assays showed a significant difference in binding potential of various antivenoms towards NNwg venom, second generation antivenomics approach was used in determining the identity of antivenom bound and unbound protein fractions. It was observed from the EC_50_ values that VINS antivenom showed better reactivity towards NNwg venom (Fig. 6a). Figure 7 (a-c) shows the UV chromatogram of VINS antivenom bound and unbound venom proteins. Though majority of the proteins are found in the bound fraction (Fig. 7c), some proteins (represented by the peaks, 1-4) are not getting recognized by the antivenom. From the MS analysis it was identified that 3FTxs, PLA_2_s and KSPIs were present in those peaks, suggesting that VINS antivenom failed to detect few proteins falling under 20 kDa region (Fig. 7a). Altogether, our study reveals that the specificity of antivenoms varies across different manufactures. Hence, as recommended by WHO, the production and purification strategies need to be streamlined for improving the efficacy of antivenoms [64].

**Figure 7.**
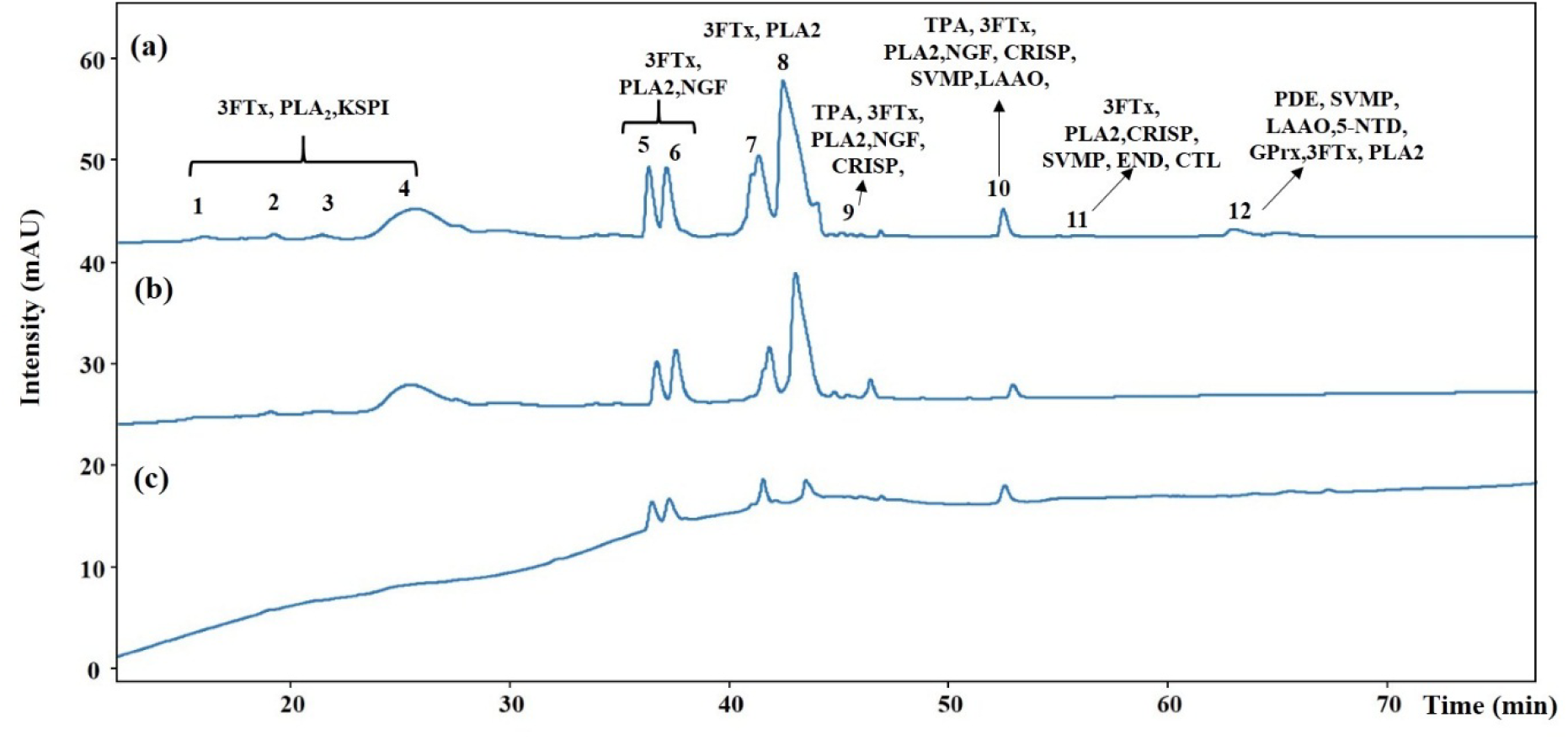
Immunoaffinity chromatography of NNwg using VINS antivenom. (a) The protein peaks observed from crude NNwg venom after reversed-phase HPLC fractionation are labelled from 1-12 (b) the unbound protein fractions observed after PBS buffer wash and (c) bound protein fractions obtained after glycine buffer wash. Proteins present in the bound and unbound fractions were compared using the chromatogram obtained from the crude venom (shown in a). The proteins present in each peaks were then identified through proteomics approaches.

## 4. Conclusion

The study demonstrates the venom proteomics and antivenom immunological cross-reactivity of *Naja naja* venom collected from the Western Ghats region in Kerala. Peptides that failed to appear in conventional database searches were subjected to homology-driven proteomic approaches using PEAKS and Novor based *de novo* sequencing followed by MS-BLAST analysis and manual validation. This finding highlights the importance of following multiple data analysis approaches in elucidating the venom proteome of snake species whose protein sequence information is limited in the public repositories. Like other reports on cobra venom, our analysis also indicated that *Naja naja* venom from the Western Ghats is predominant with proteins belonging to three-finger toxin family. To date, there are a total of eight published reports on cobra venom proteome from different geographical locations of India (Table 3). Though the species remains the same, detailed venom proteome profiling of *Naja naja* venom from different parts of India reports that venom composition varies according to the geographical location. The immunological cross-reactivity studies indicated that there are variations across Indian polyvalent antivenoms in detecting and binding to the antigenic epitopes present in *Naja naja* venom proteins. Altogether, our study reveals that the specificity of antivenoms varies across different manufacturers. This highlights the importance of improving antivenom preparations against *Naja naja* envenomation in India.

**Table 3:** Summary of cobra venom proteomic studies from India.

| Sl. No. | Venom family | *Naja naja* |  |  |  |  |  |  |  |
| --- | --- | --- | --- | --- | --- | --- | --- | --- | --- |
|  |  | West India [8] | West India [14] | East India [9,10] | North India [11] | South India [5] | South India [13] | Kentucky Reptile Zoo [12] | Western Ghats # |
| 1 | **PLA<sub>2</sub>** | + | + | + | + | + | + | + | + |
| 2 | **PLB** | + | - | + | - | - | - | + | + |
| 3 | **SVMP** | + | + | + | + | + | + | + | + |
| 4 | **SVSP** | + | + | - | - | + | + | + | + |
| 5 | **CRISP** | + | + | + | + | + | + | + | + |
| 6 | **Cystatin** | + | - | + | - | - | + | + | - |
| 7 | **KSPI** | + | + | + | + | + | + | + | + |
| 8 | **3FTx** | + | + | + | + | + | + | + | + |
| 9 | **NGF** | + | - | + | + | + | + | + | + |
| 10 | **ACE** | + | - | + | - | - | + | + | + |
| 11 | **PDE** | + | - | + | - | + | - | - | + |
| 12 | **LAAO** | + | + | + | - | + | + | + | + |
| 13 | **CVF** | + | + | + | - | + | + | + | + |
| 14 | **5-NU** | + | + | + | - | + | + | + | + |
| 15 | **GPrx** | - | - | - | - | - | - | - | + |
| 16 | **Natriuretic peptide** | - | + | + | - |  | + | - | - |
| 17 | **AP** | + | - | - | - | + | - | - | - |
| 18 | **OLP** | + | + | + | - | + | + | + | + |
| 19 | **CTL** | - | - | - | - | - | - | + | + |
| 20 | **END** | - | - | - | - | - | - | - | + |
| 21 | **Hyal** | - | - | - | - | - | - | + | - |
| 22 | **VEGF** | - | - | - | - | - | - | + | - |
|  | **\*- cobra venom obtained from Kentucky Reptile Zoo, #-current study** |  |  |  |  |  |  |  |  |

**Table 4:** Summary of cobra venom proteomic studies from India.

| Table 4: Summary of cobra venom proteomic studies from India. |  |  |  |  |  |
| --- | --- | --- | --- | --- | --- |
| Sl. No. | Geographical location/place from venom was collected or procured | Number of snakes used for the study | Total number of proteins identified | Number of venom families identified | Reference |
| 1 | North-Western (Rajasthan, Gujarat) | 17 | 26 | 6 | [11] |
| 2 | Eastern India (Nadia District, West Bengal)- Calcutta Snake park | 4 | 43 | 15 | [9] |
| 3 | Eastern India-(Burdwan District, West Bengal)- Calcutta snake park, Kolkata | 4 | 52 | 14 | [10] |
| 4 | Southern India-Tamil Nadu-Irula snake catcher's society | Pooled venom | 81 | 13 | [5] |
| 5 | Southern India-Tamil Nadu-Irula snake catcher's society | 4 | 45 | 14 | [13] |
| 6 | Western India-Haffkine Institute, Mumbai | 4 | 54 | 16 | [8] |
| 7 | Maharashtra (West India) | 5 | 56 | 11 | [14] |
| 8 | Indian cobra venom obtained from Kentucky Reptile Zoo (exact geographical location not mentioned) | Pooled venom | 48 | 17 | [12] |
| 9 | Western Ghats region in Kerala (India) | 2 | 115 | 17 | Current study |

## Acknowledgements

The authors would like to acknowledge Sri. Mata Amritanandamayi Devi, The Chancellor of Amrita Vishwa Vidyapeetham, for being the inspiration behind this study. We are grateful to the forest department of Kerala for their assistance during the field visits during the time of venom collection. The authors would like to thank Dr. Dileepkumar Raveendran and Indriyam Biologics for providing the venom, various chemicals and reagents including antivenoms, which were used for all the immunological cross-reactivity assays. MV thank Amrita Vishwa Vidyapeetham and Amrita Centre for Research and Development for providing funding in the form of fellowship. The authors also acknowledge Amrita Agilent Analytical Research Centre for the mass spectrometric data collection and analysis. The authors would like to thank Dr. Sanu K. Shaji, Ms.Meera Venugopal and Mr.Vishnu Raj for helping with the Western blotting experiments. Financial assistance in executing this project provided by Amrita Vishwa Vidyapeetham is also duly acknowledged.

## Conflict of Interest

There are no potential conflict of interests.

---

**Supplementary Table 1:**
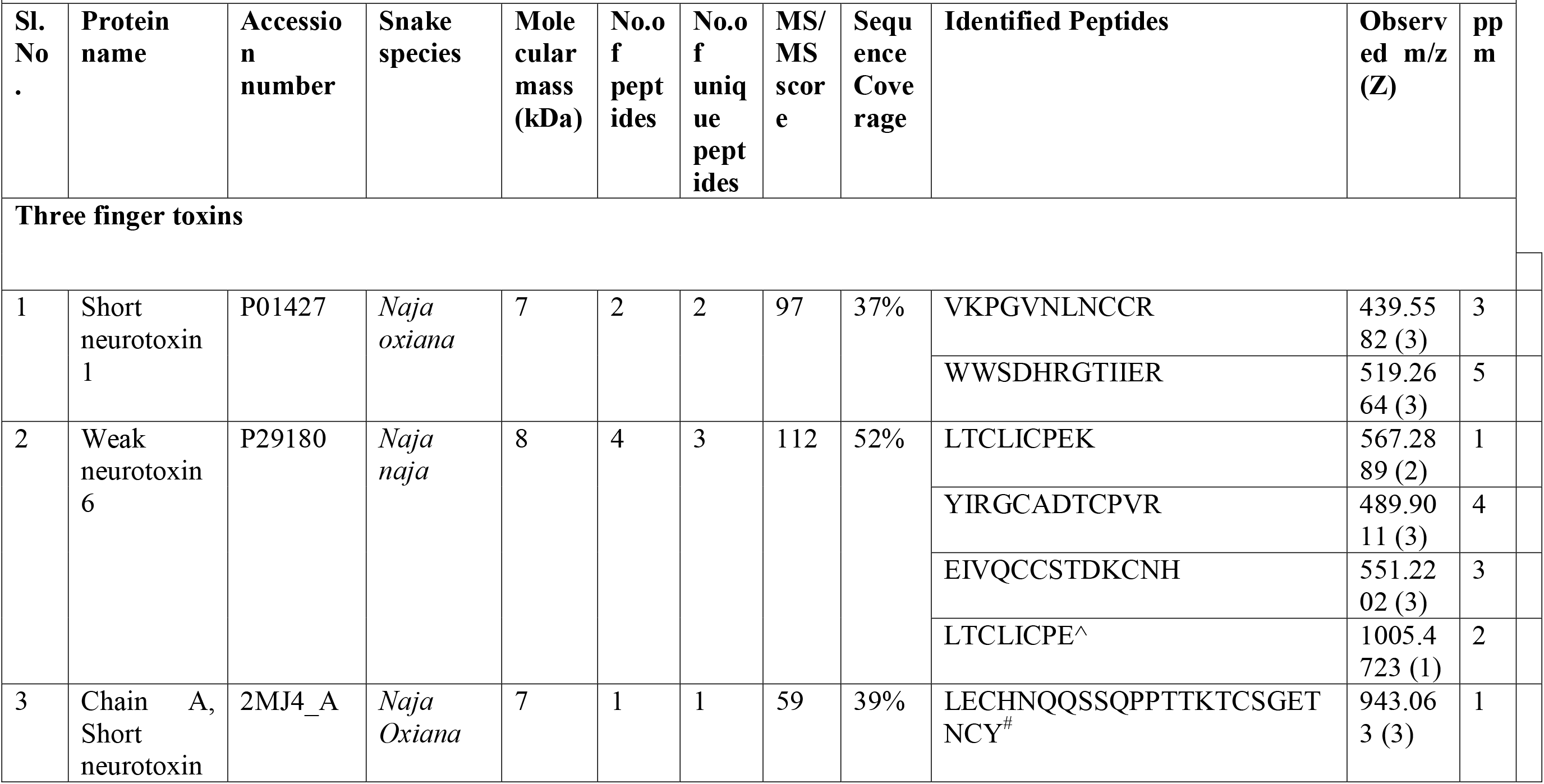
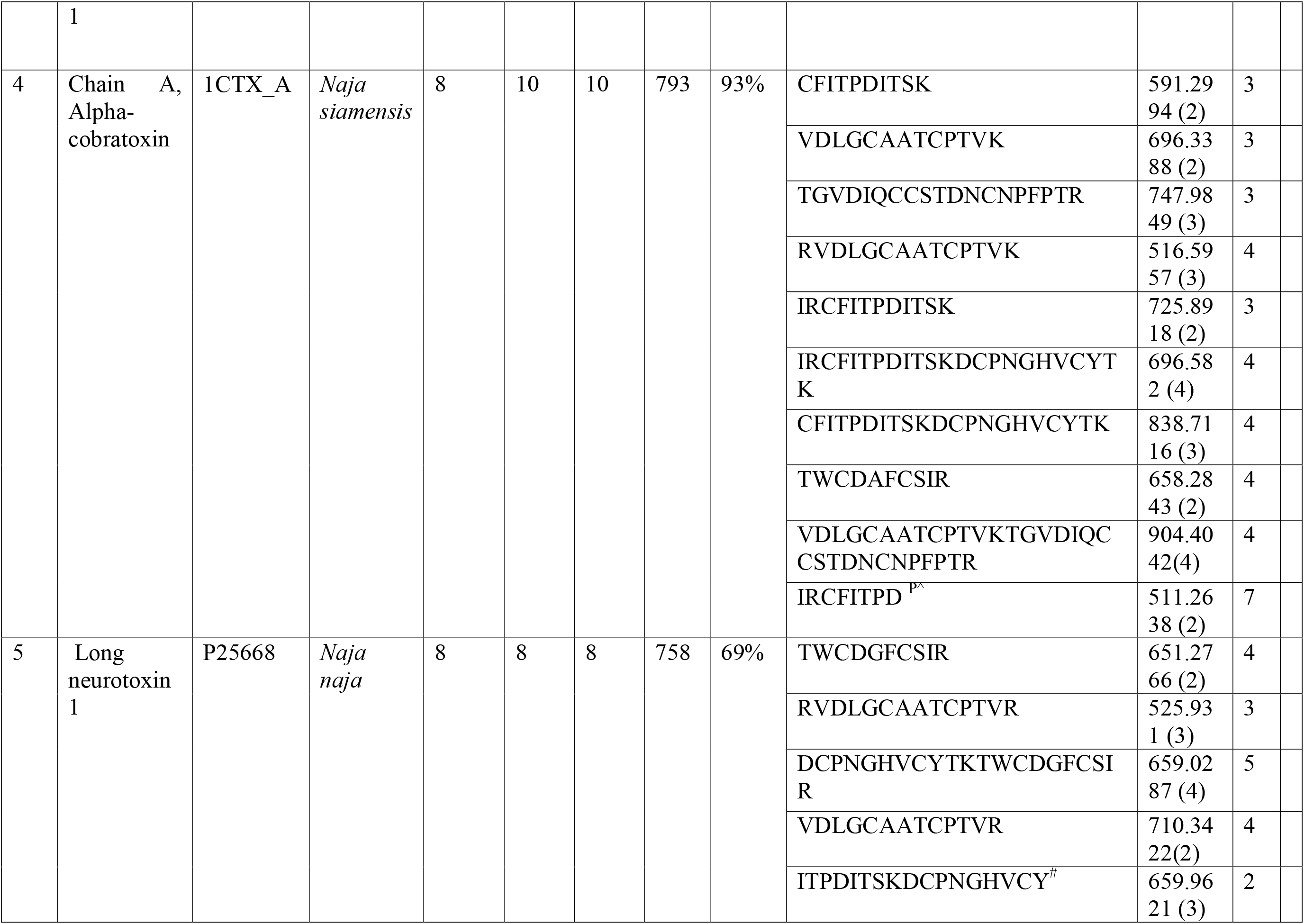
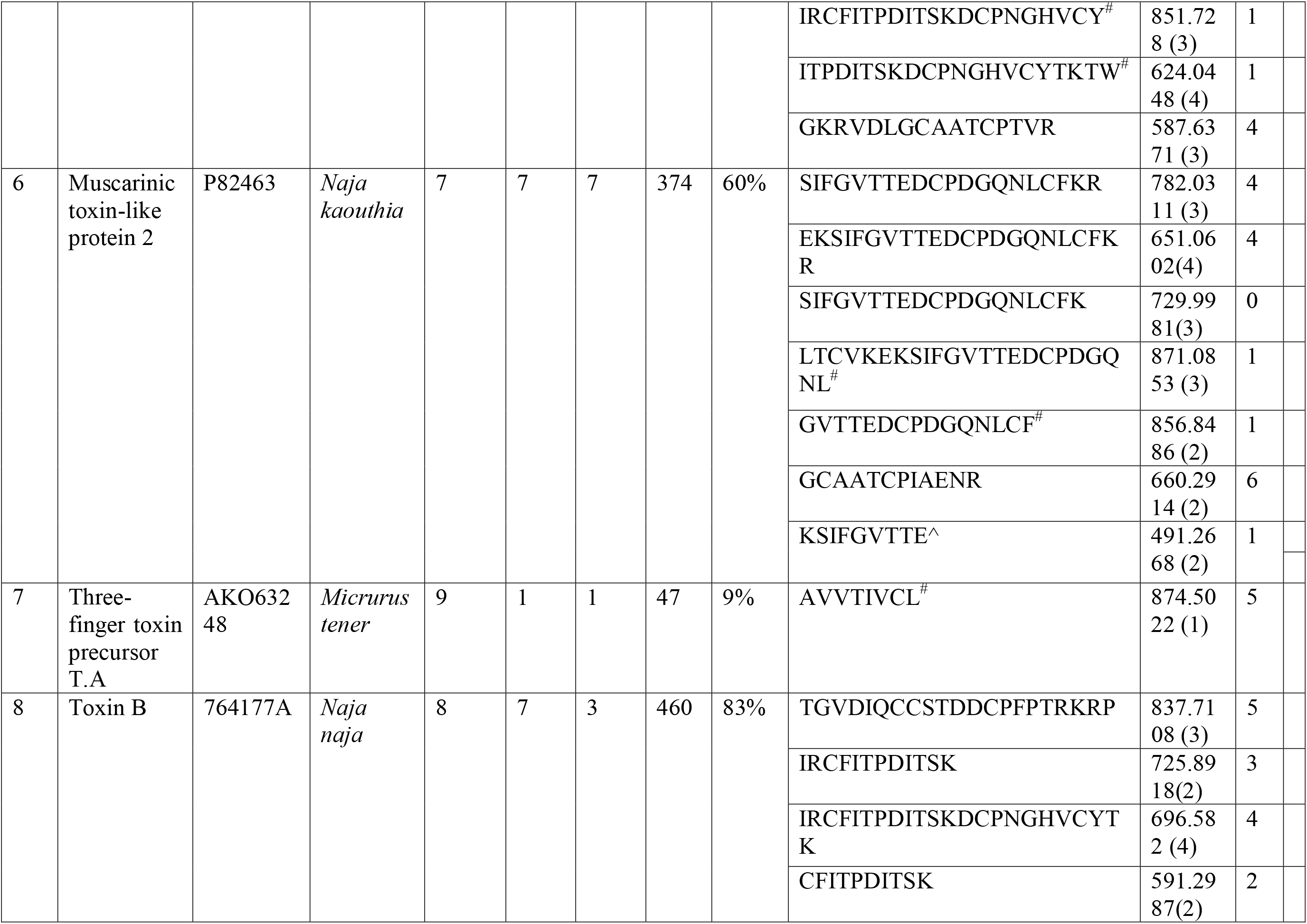
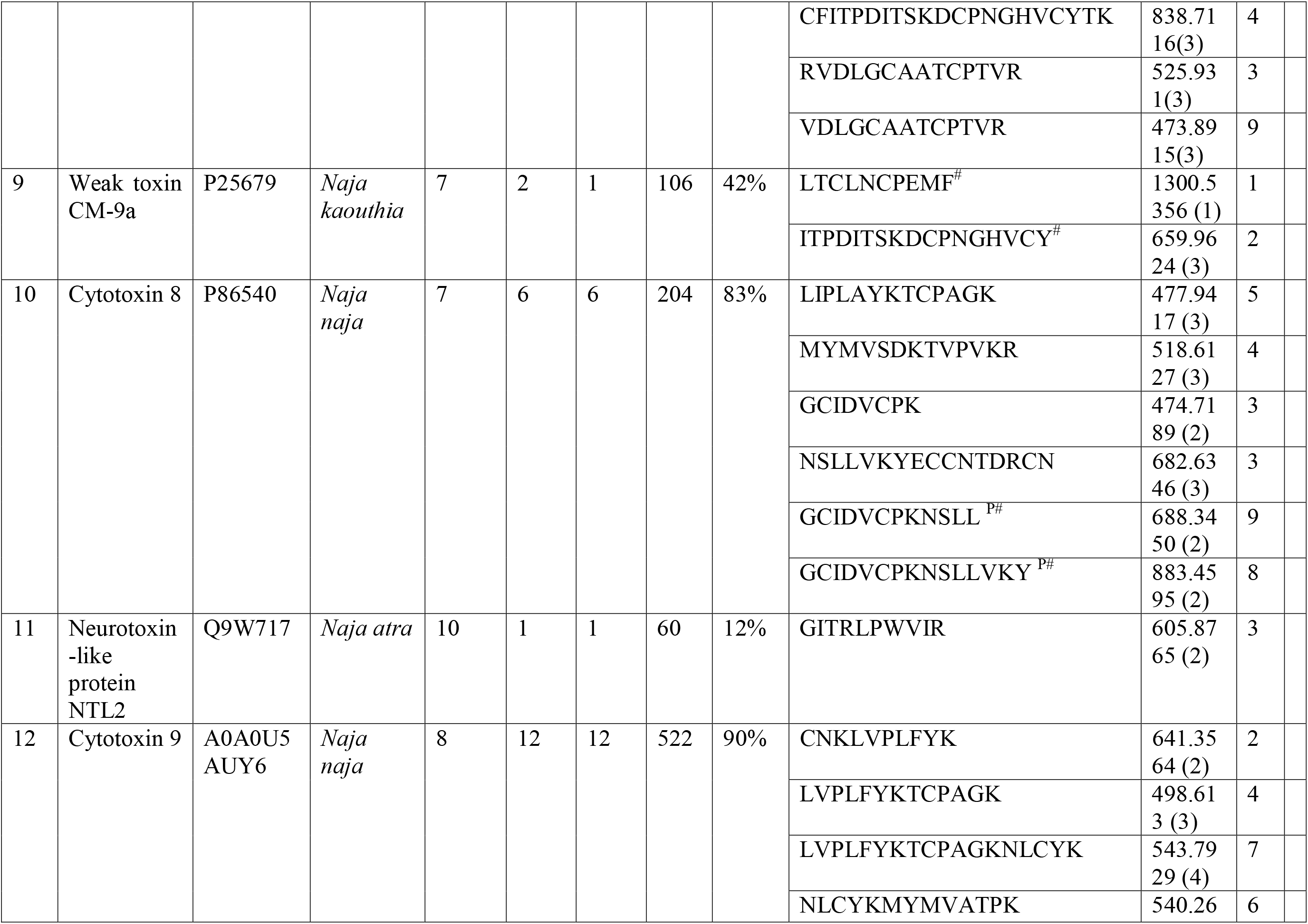
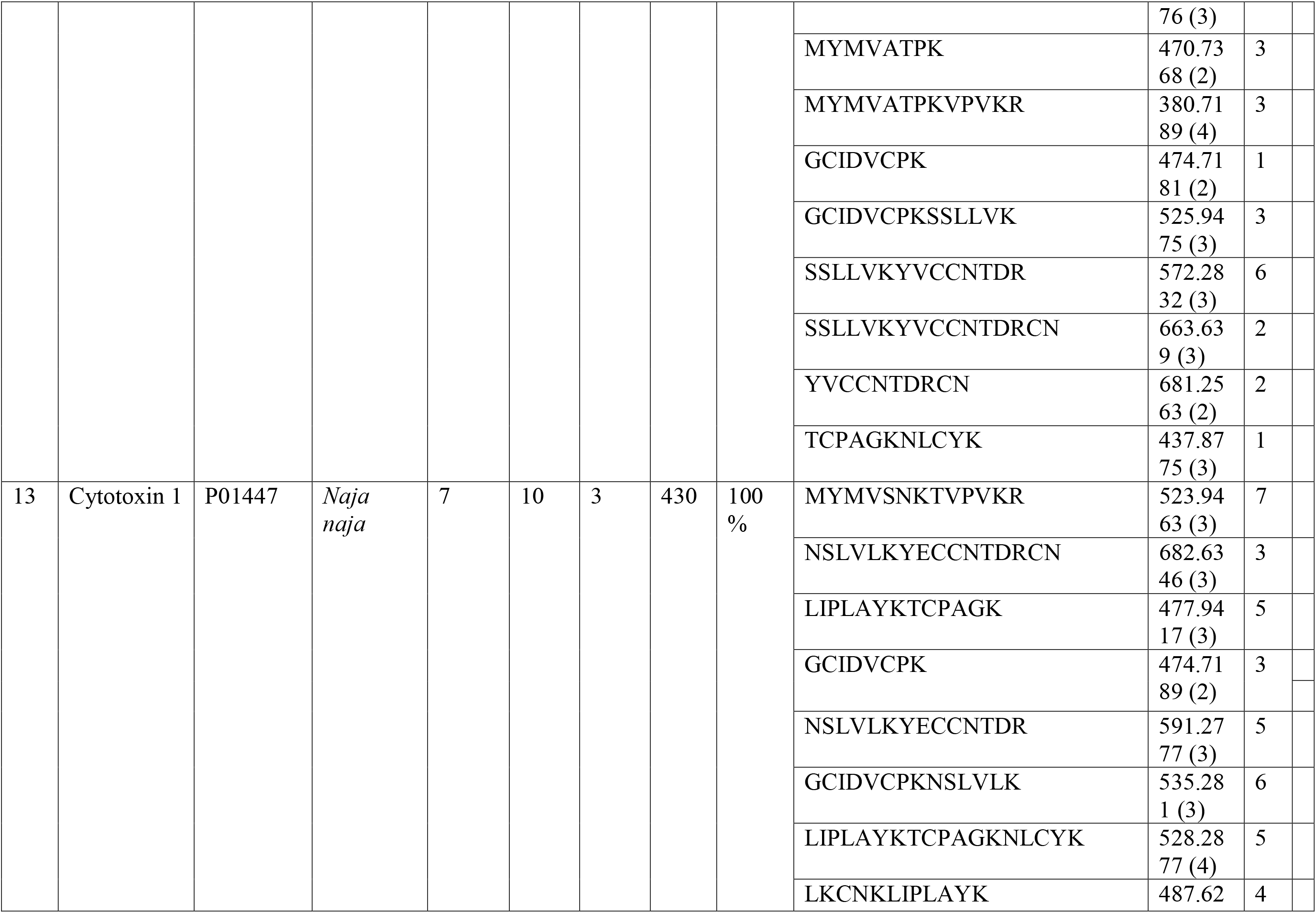
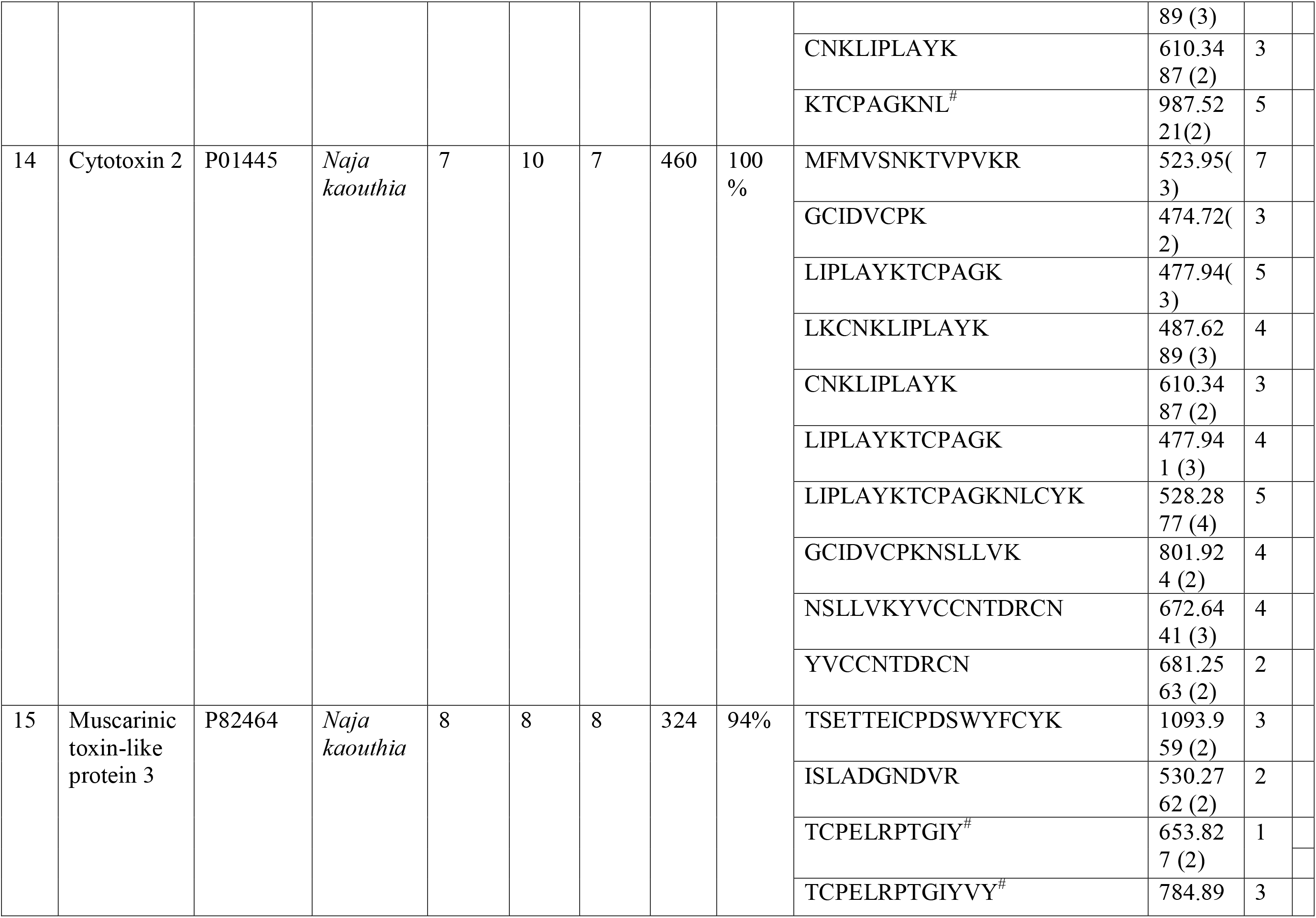
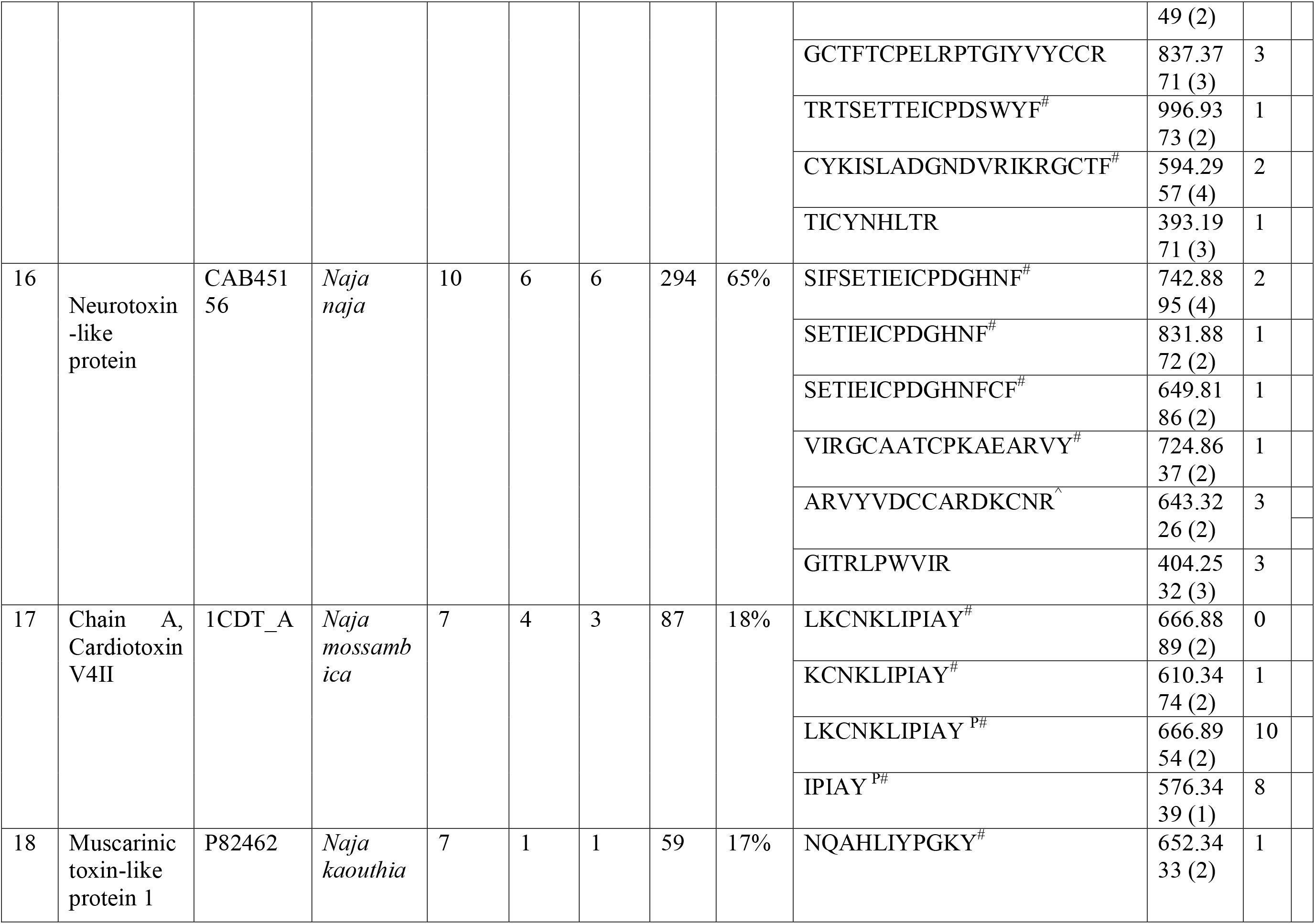
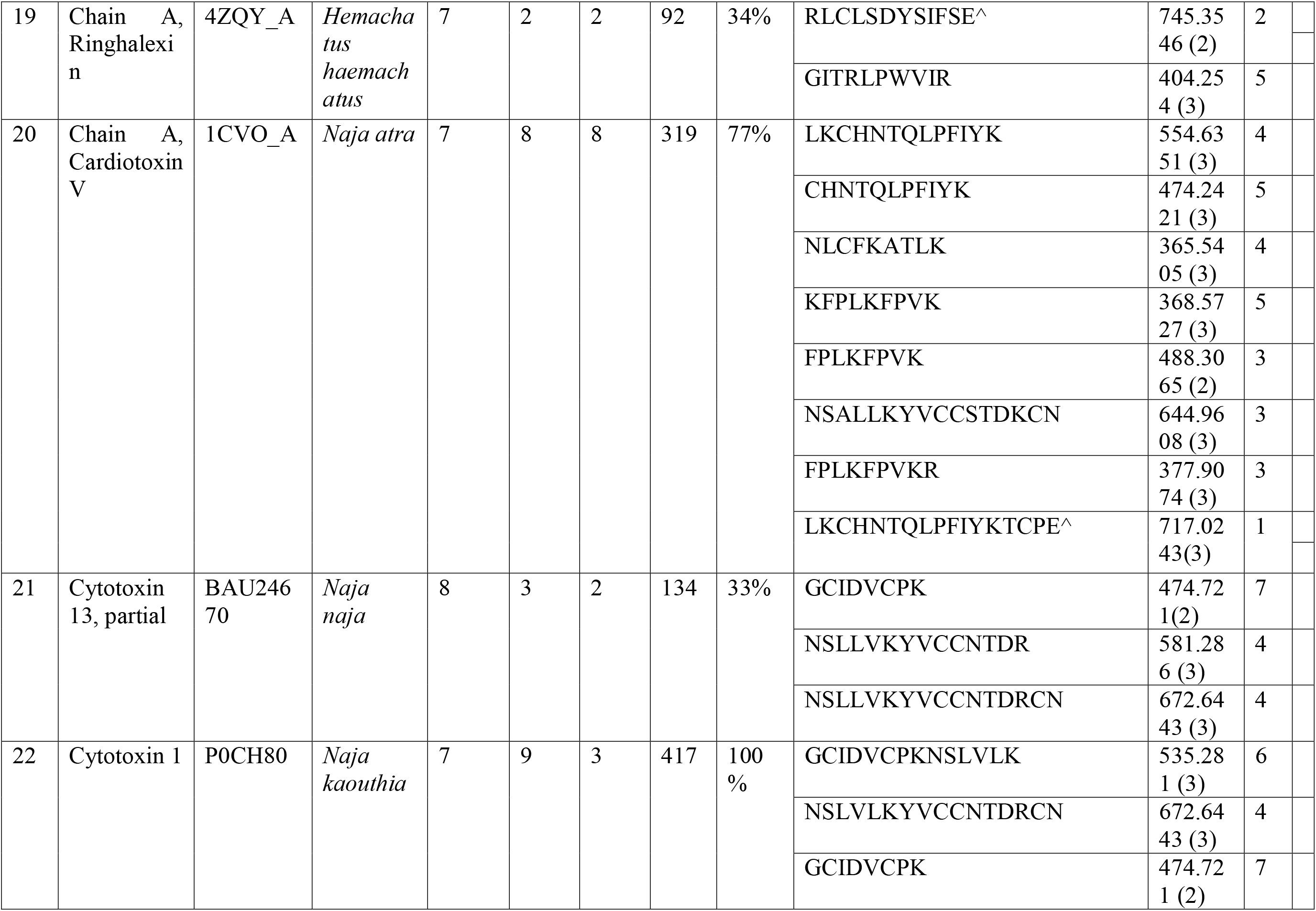
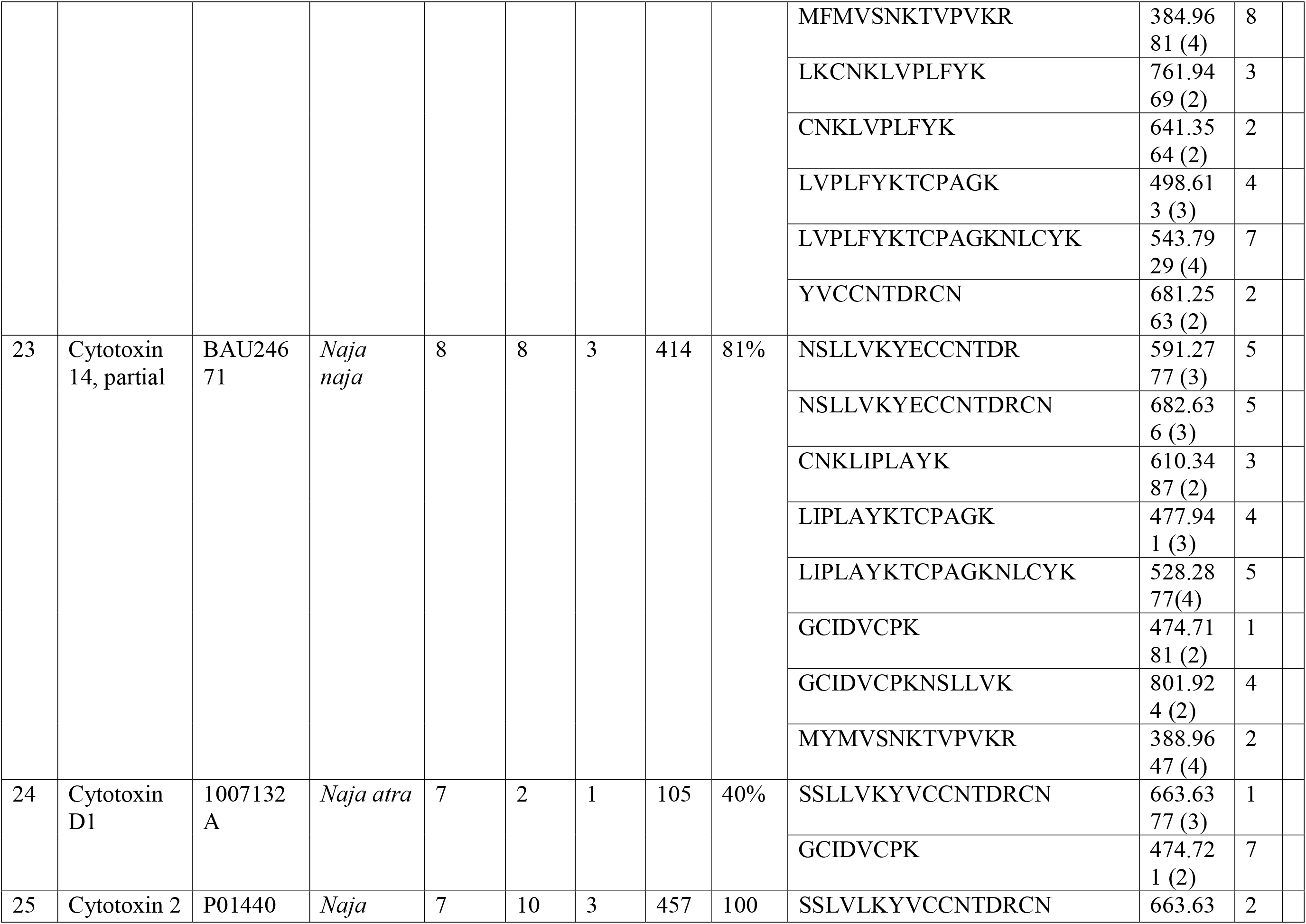
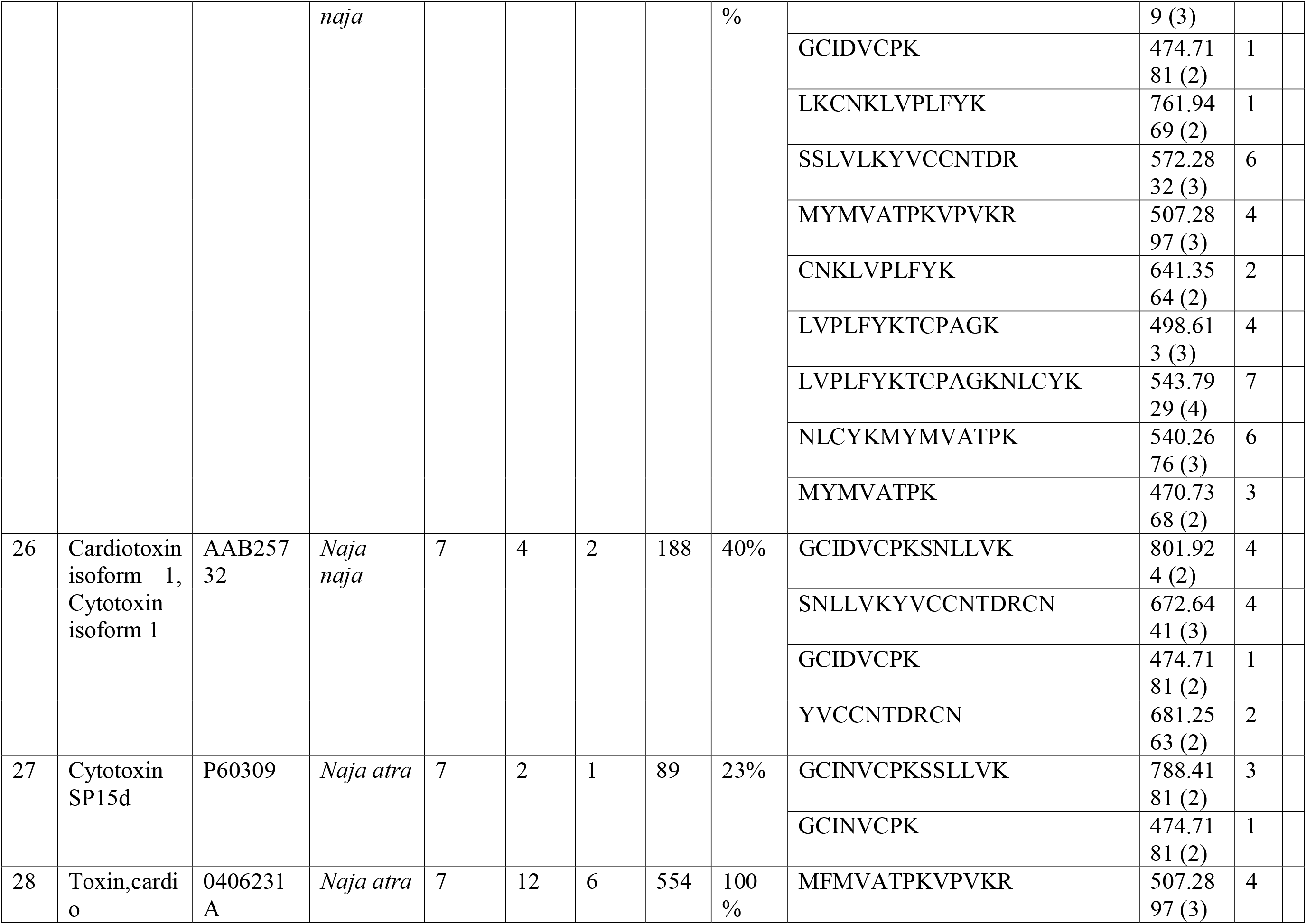
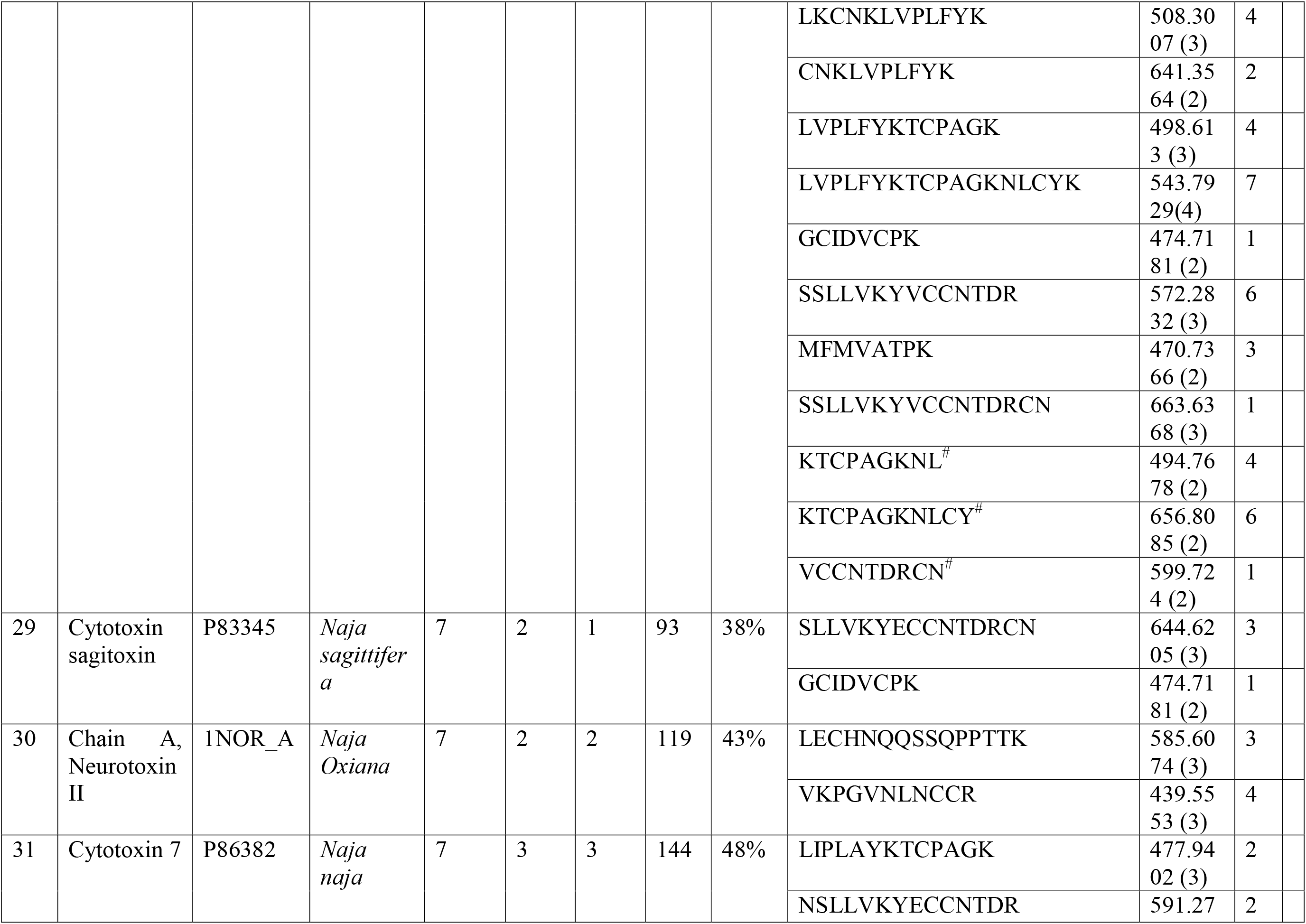
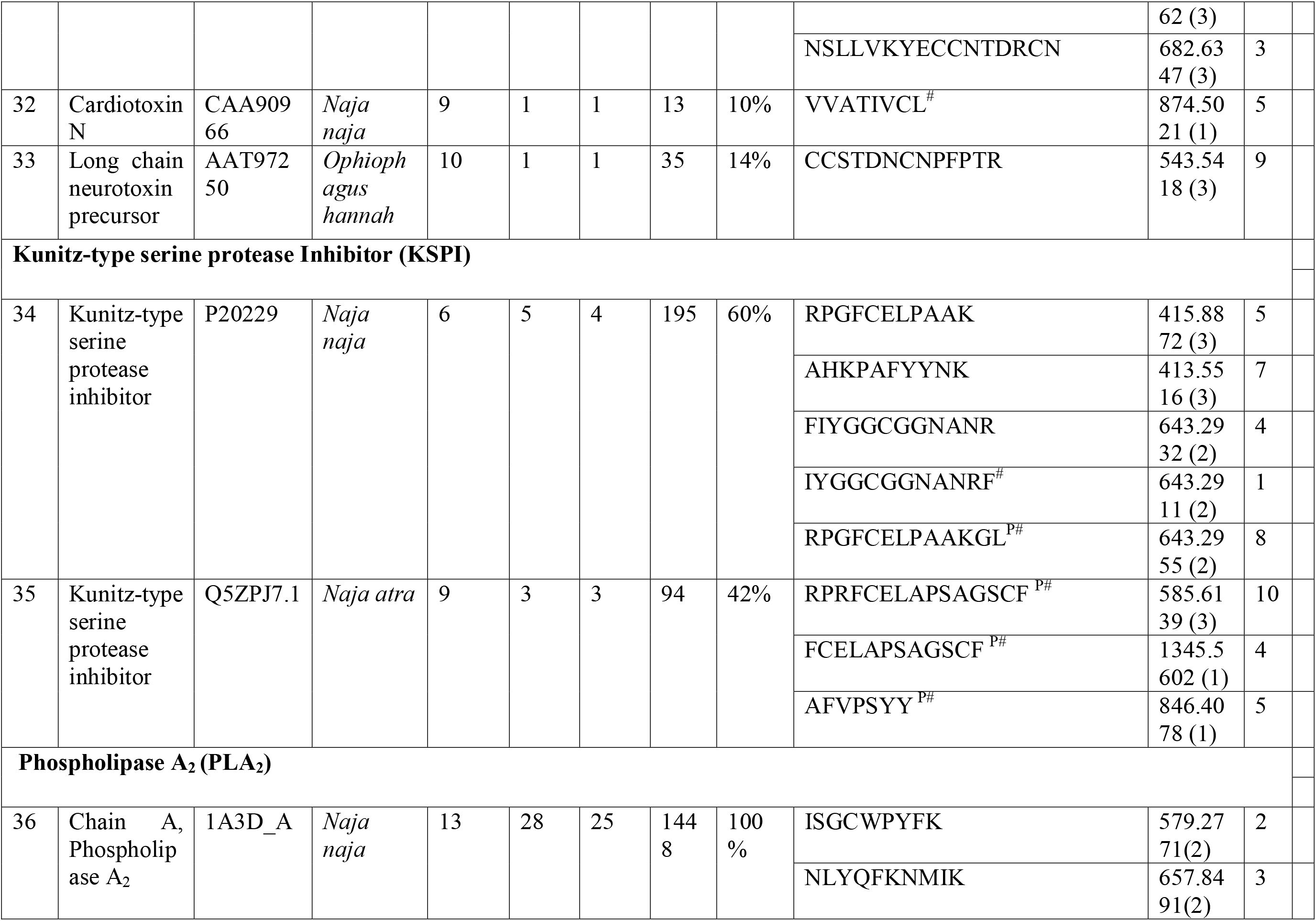
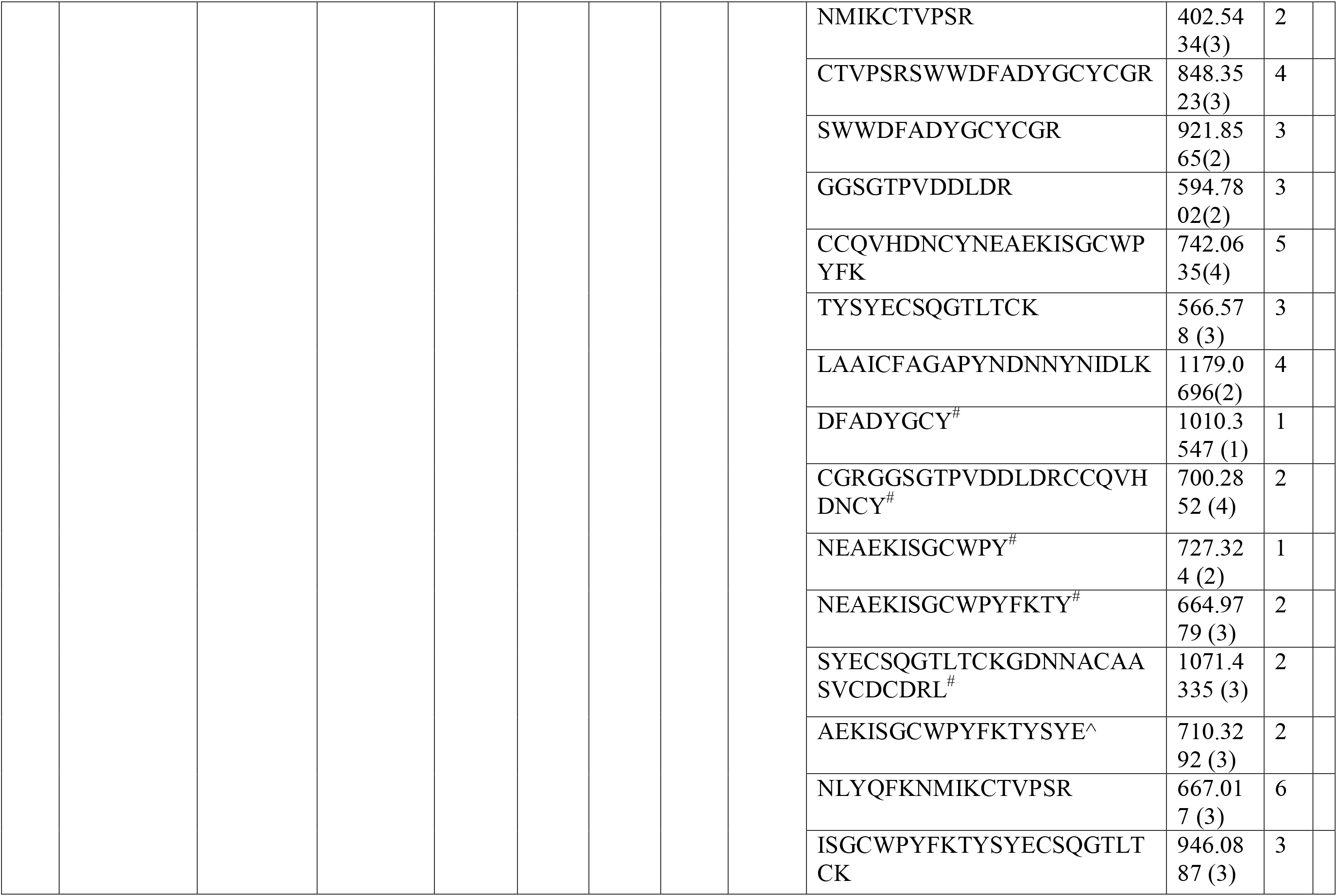
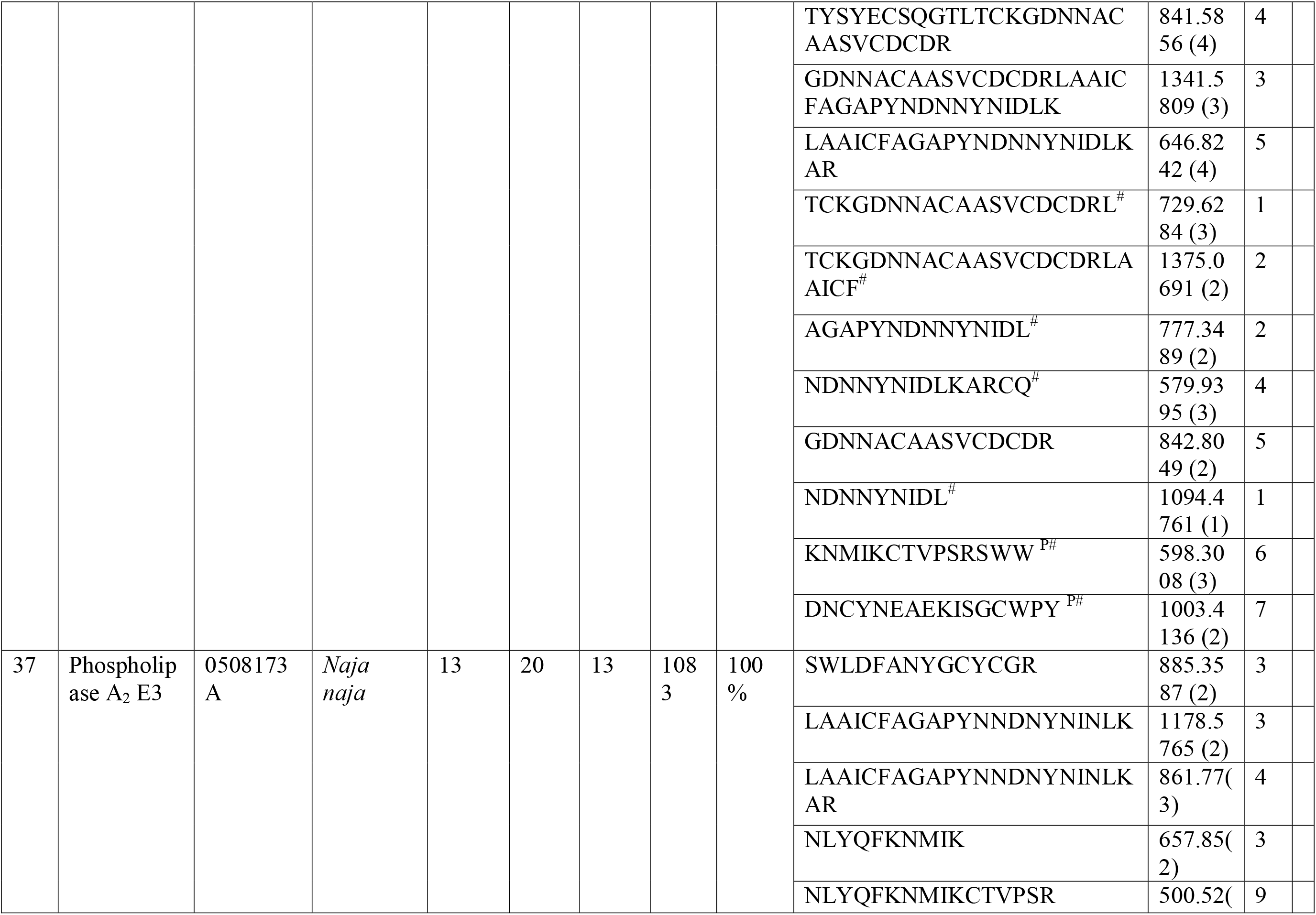
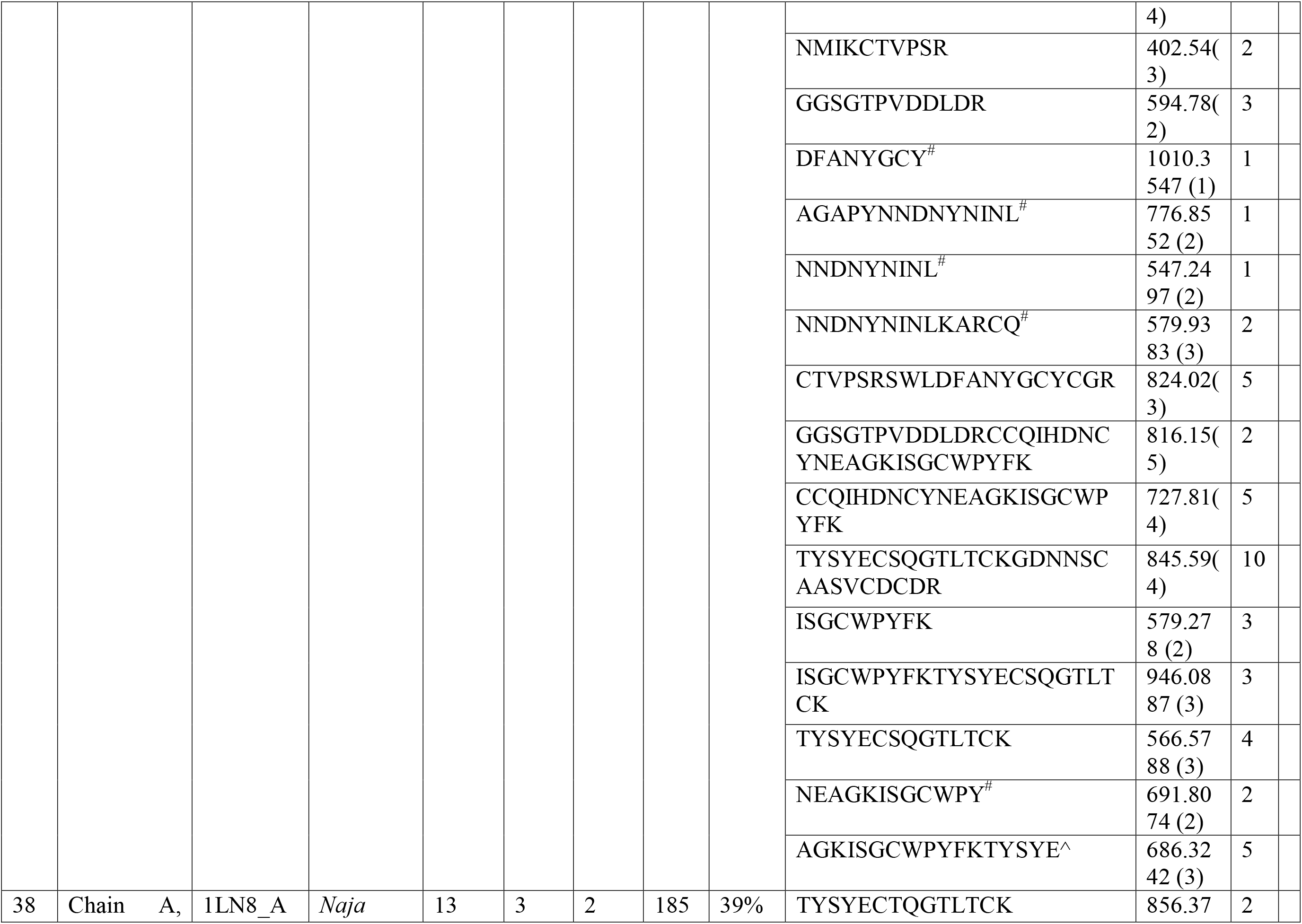
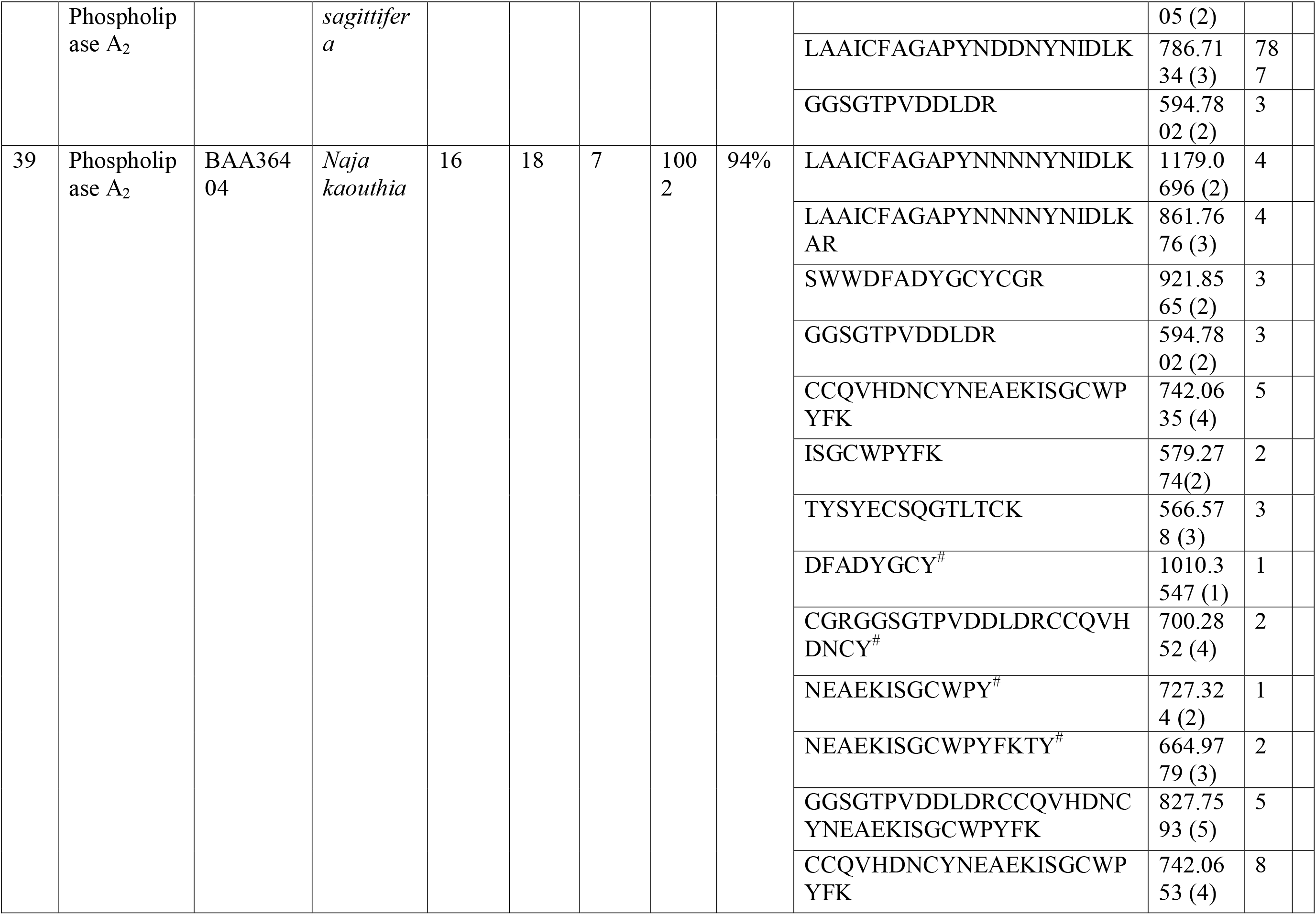
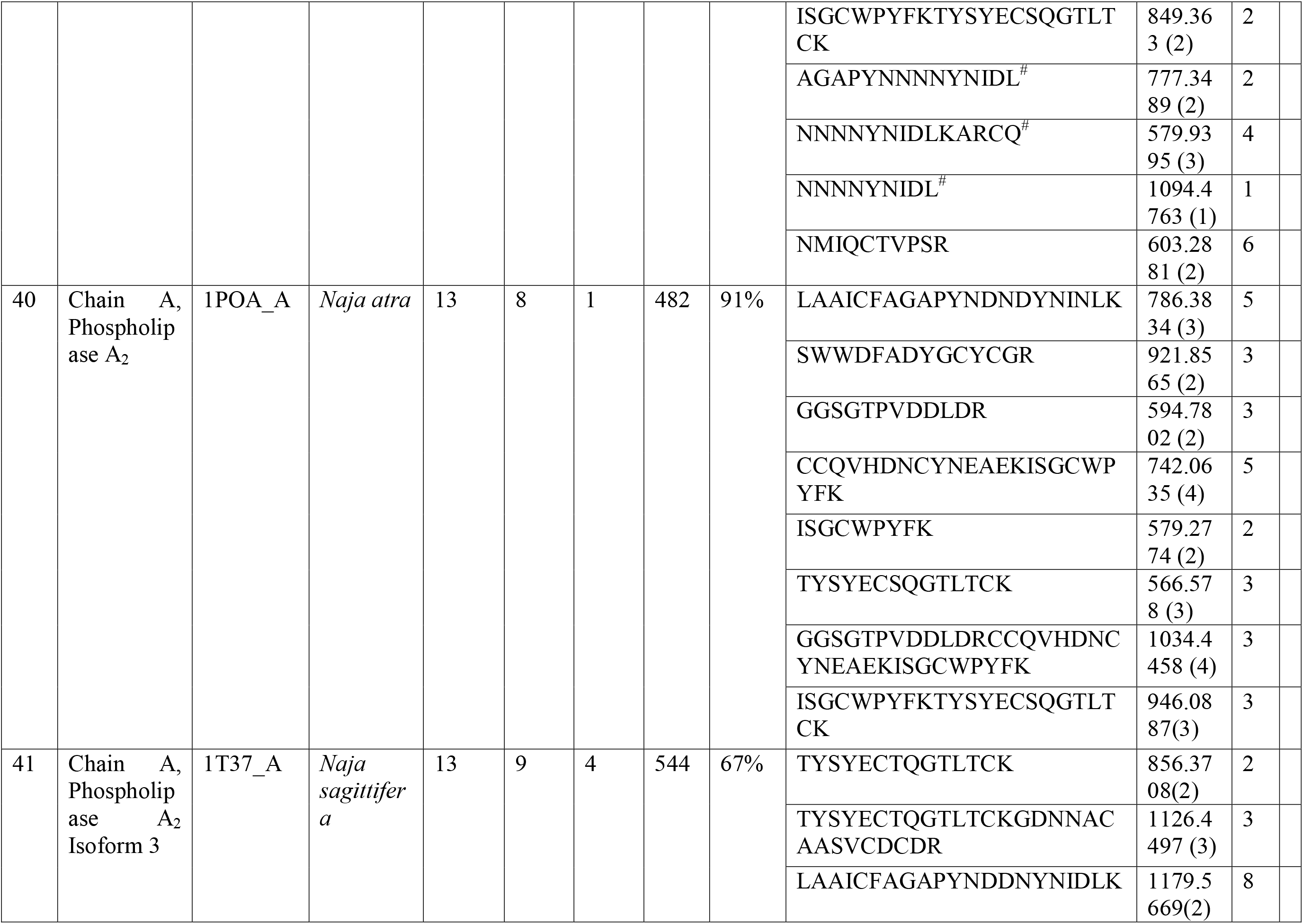
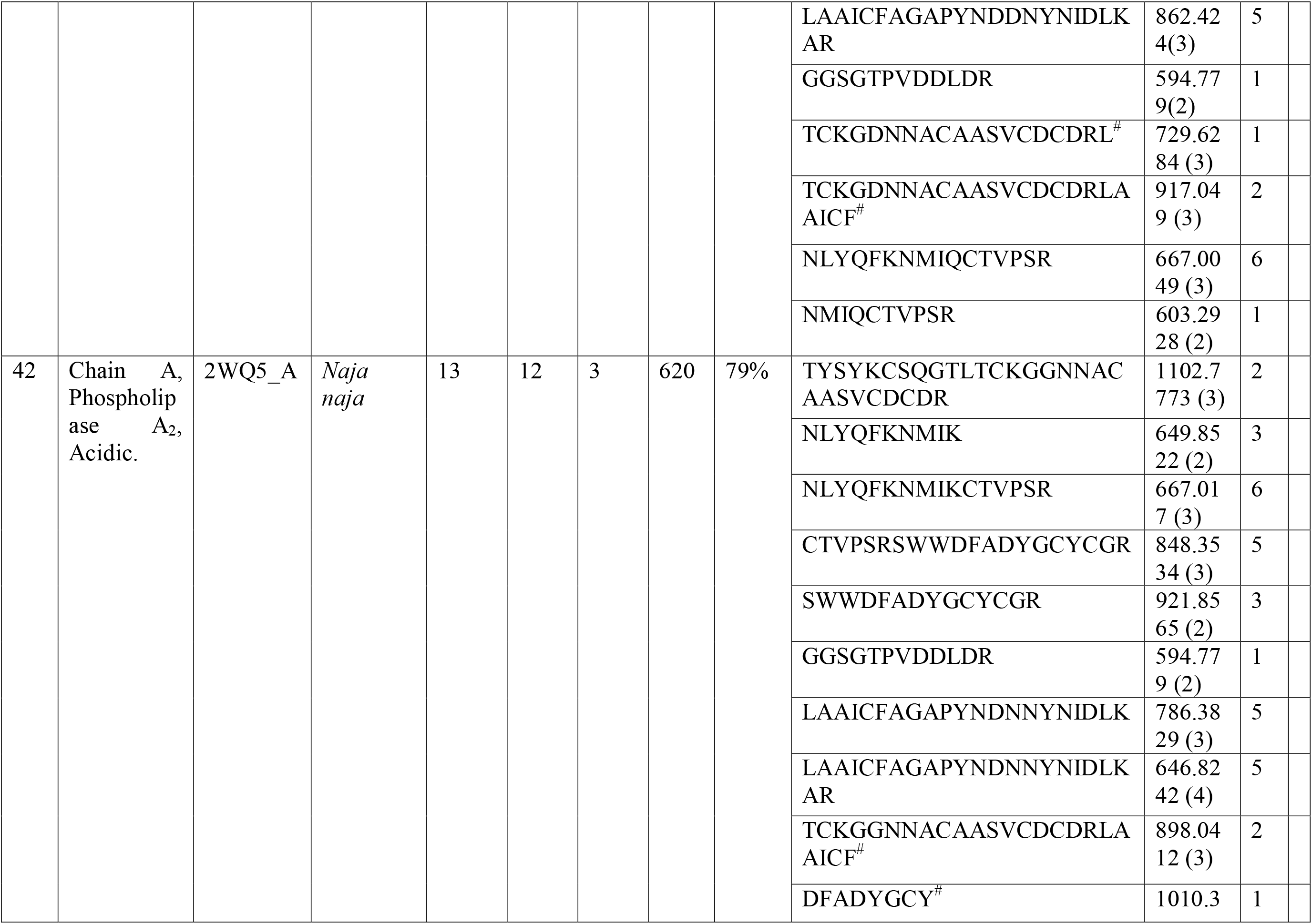
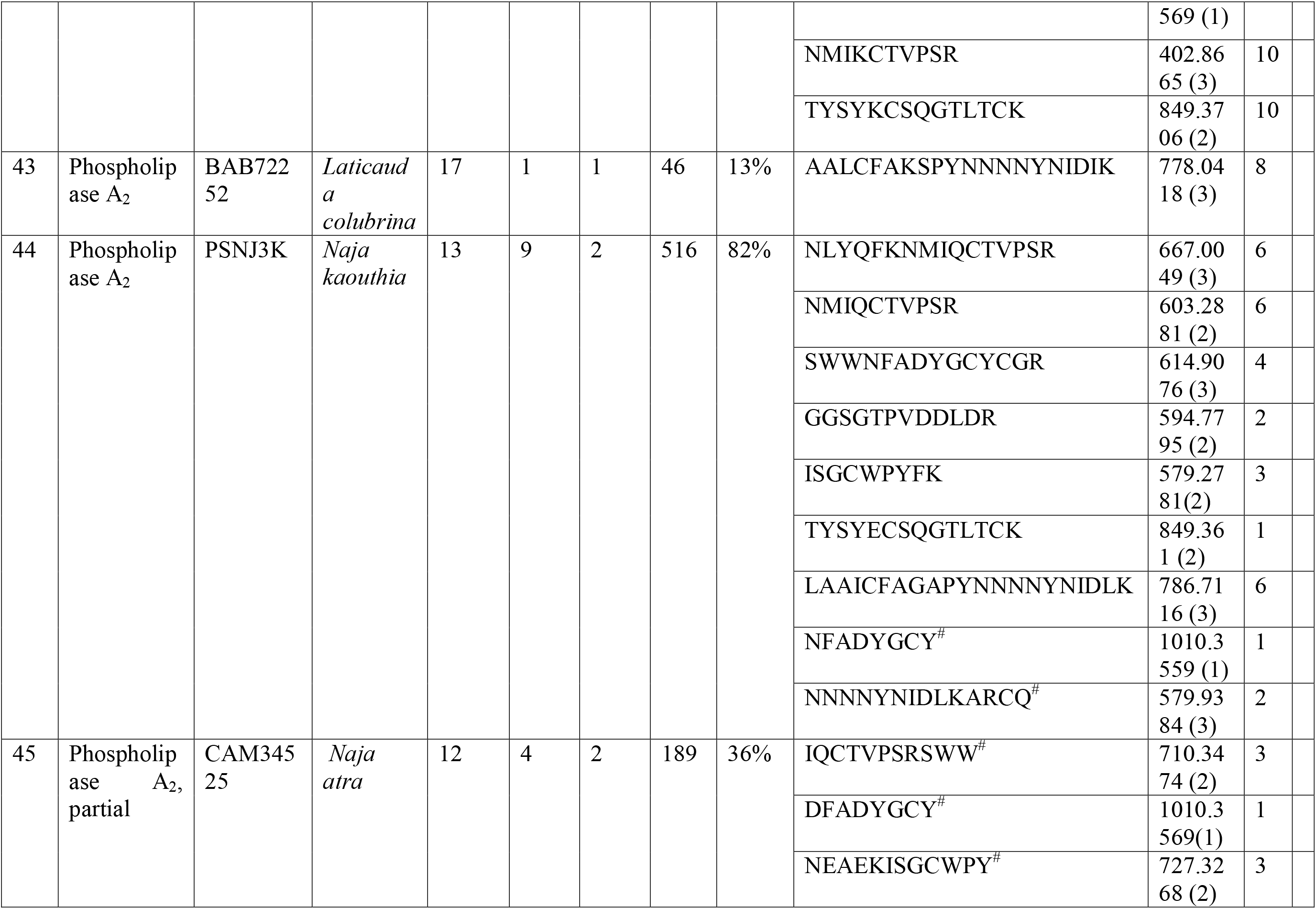
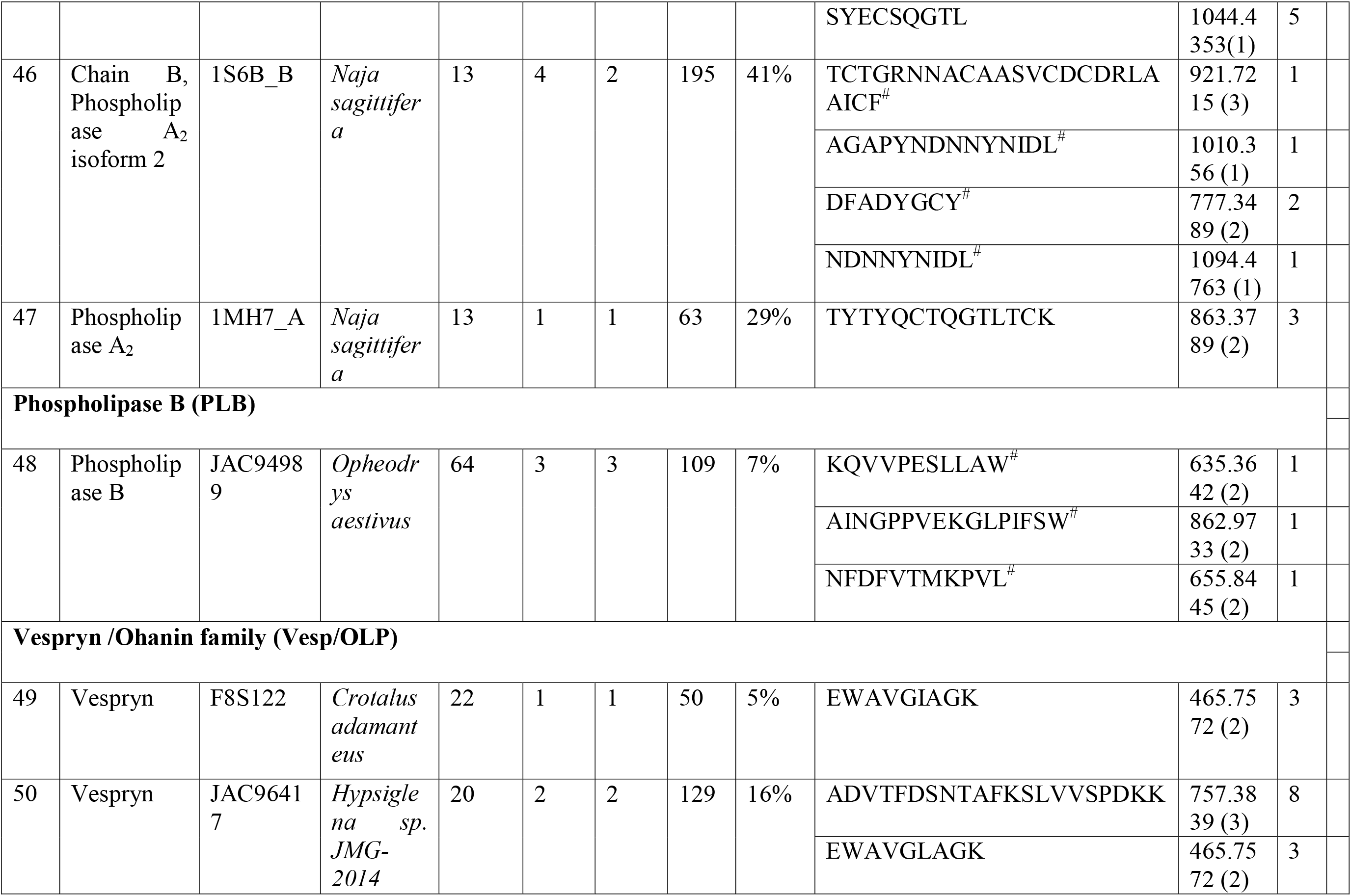
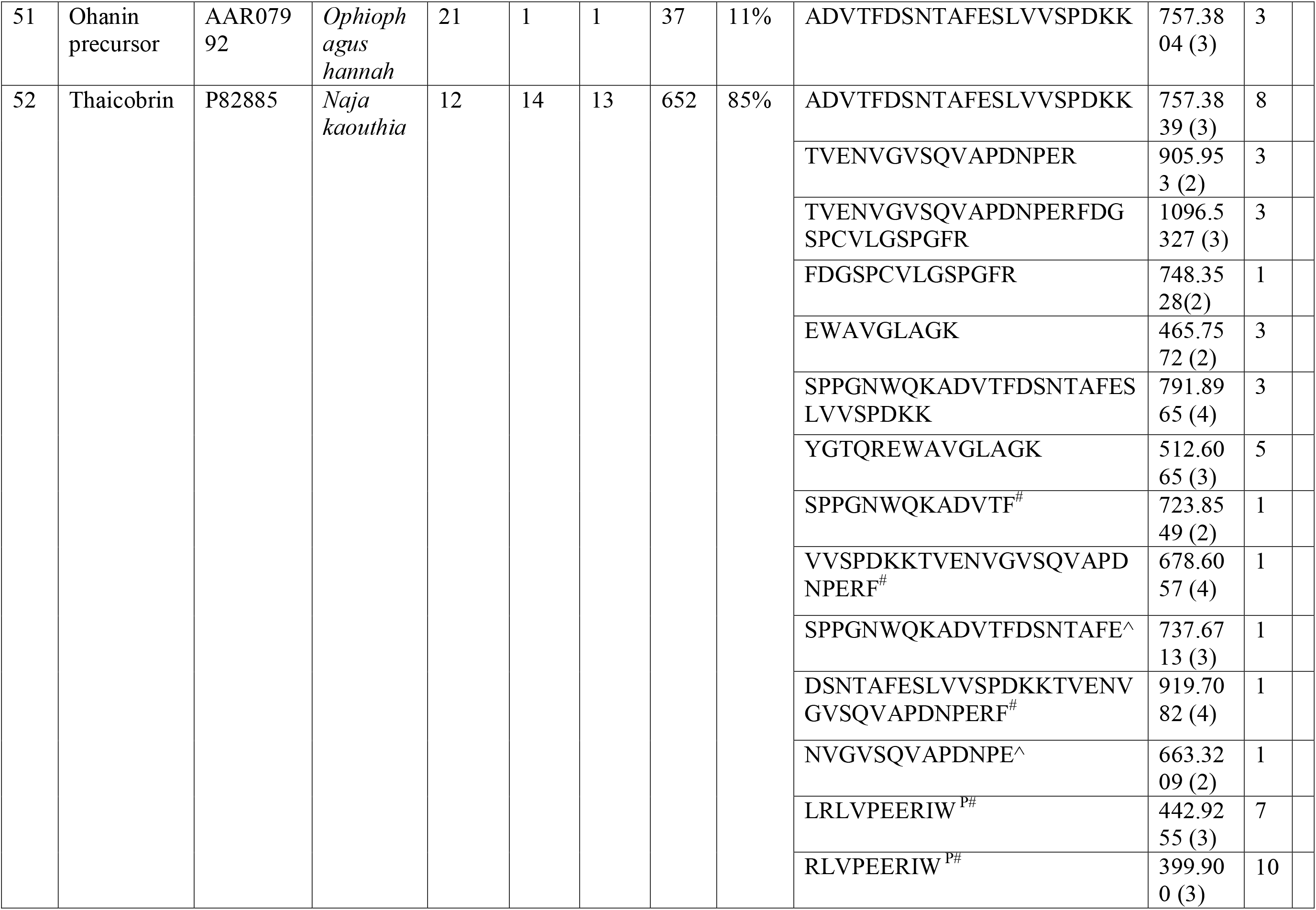
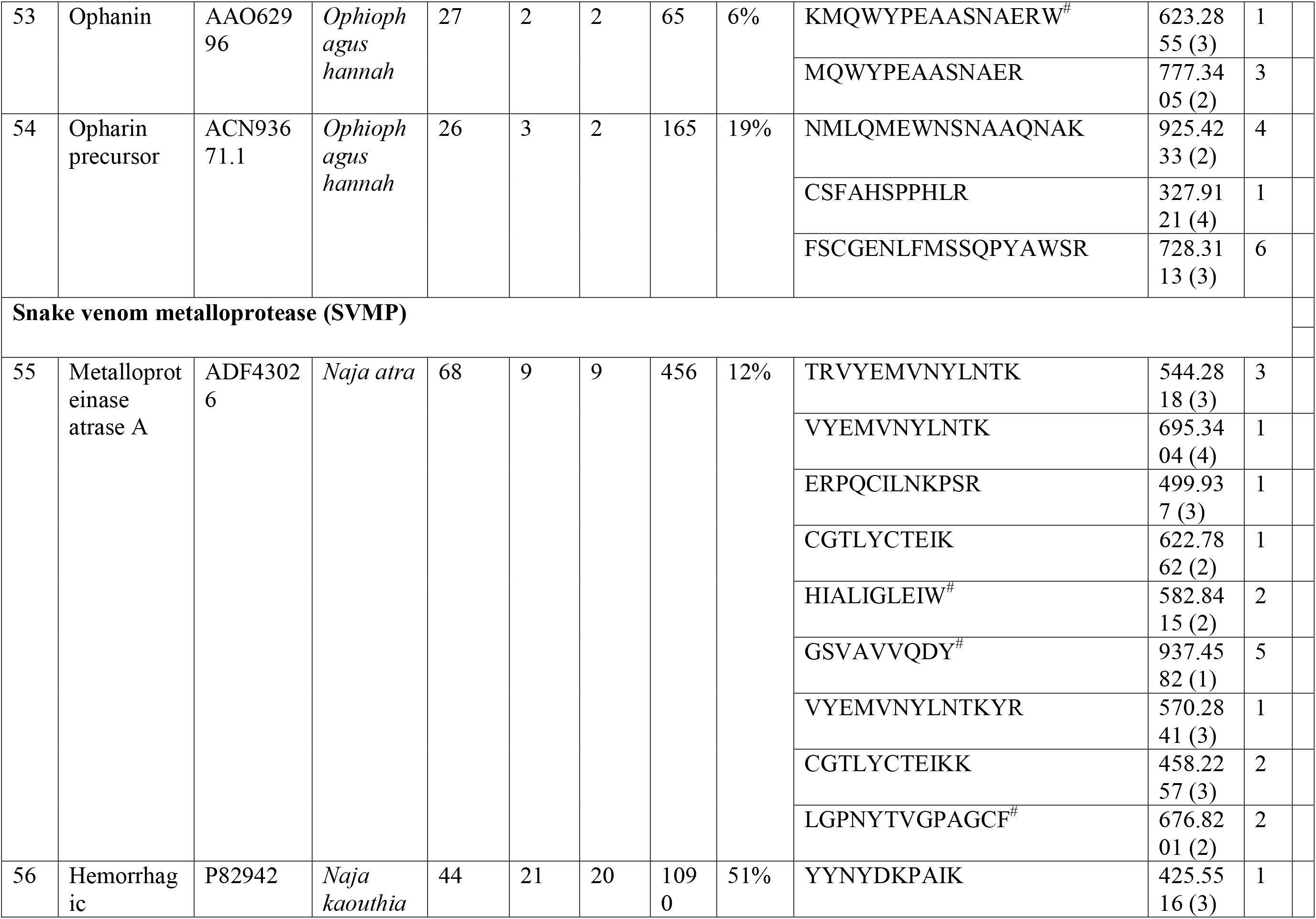
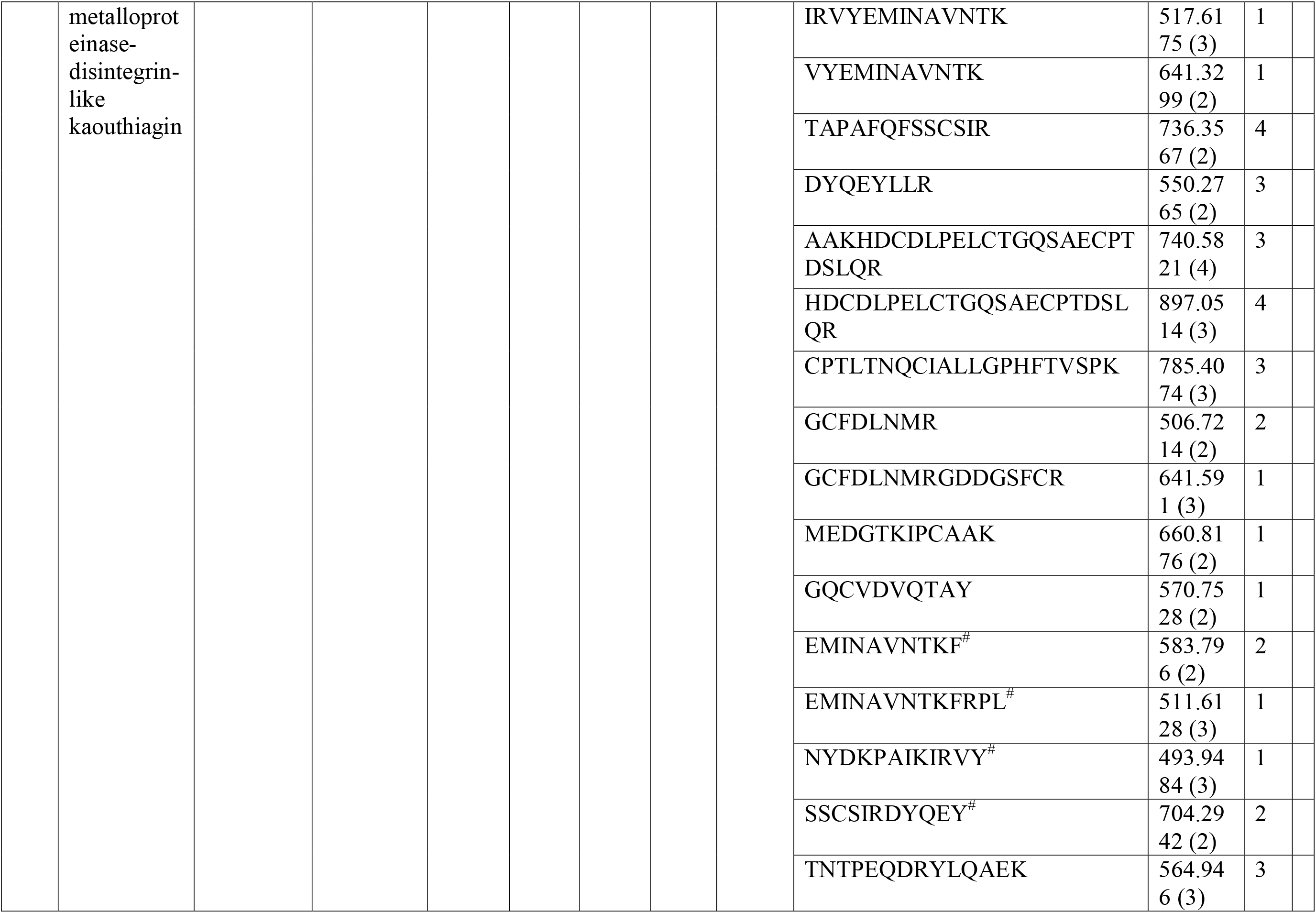
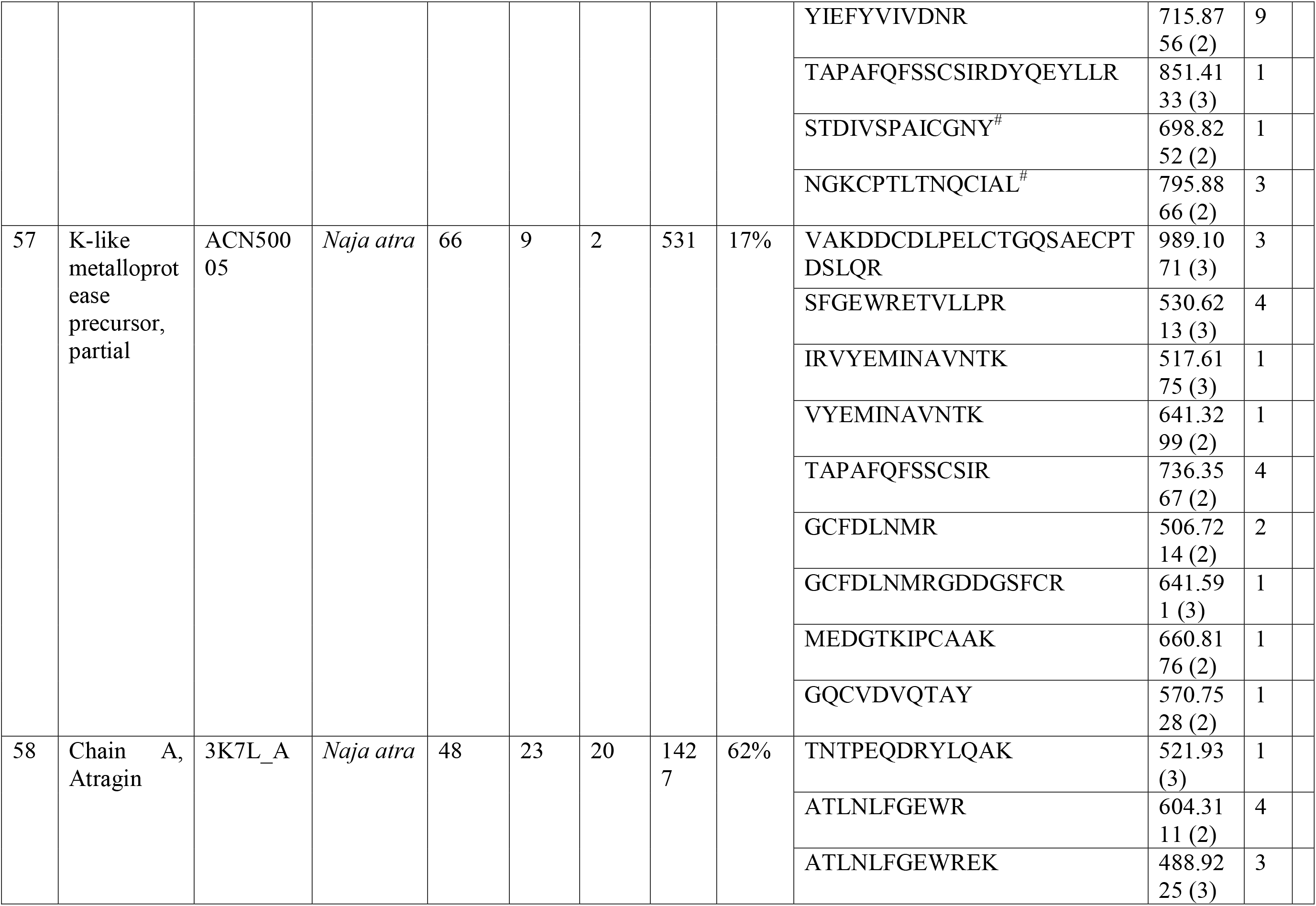
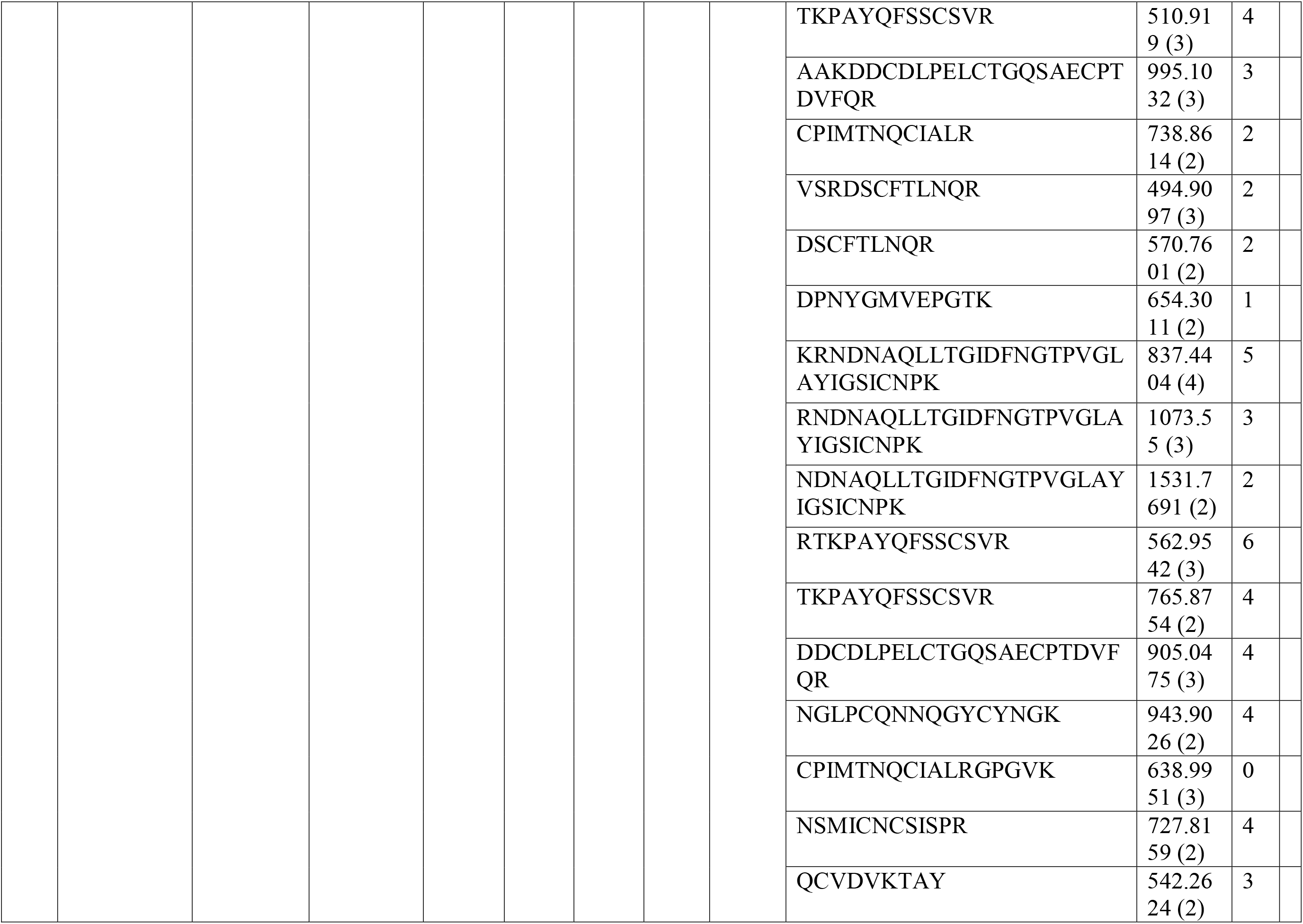
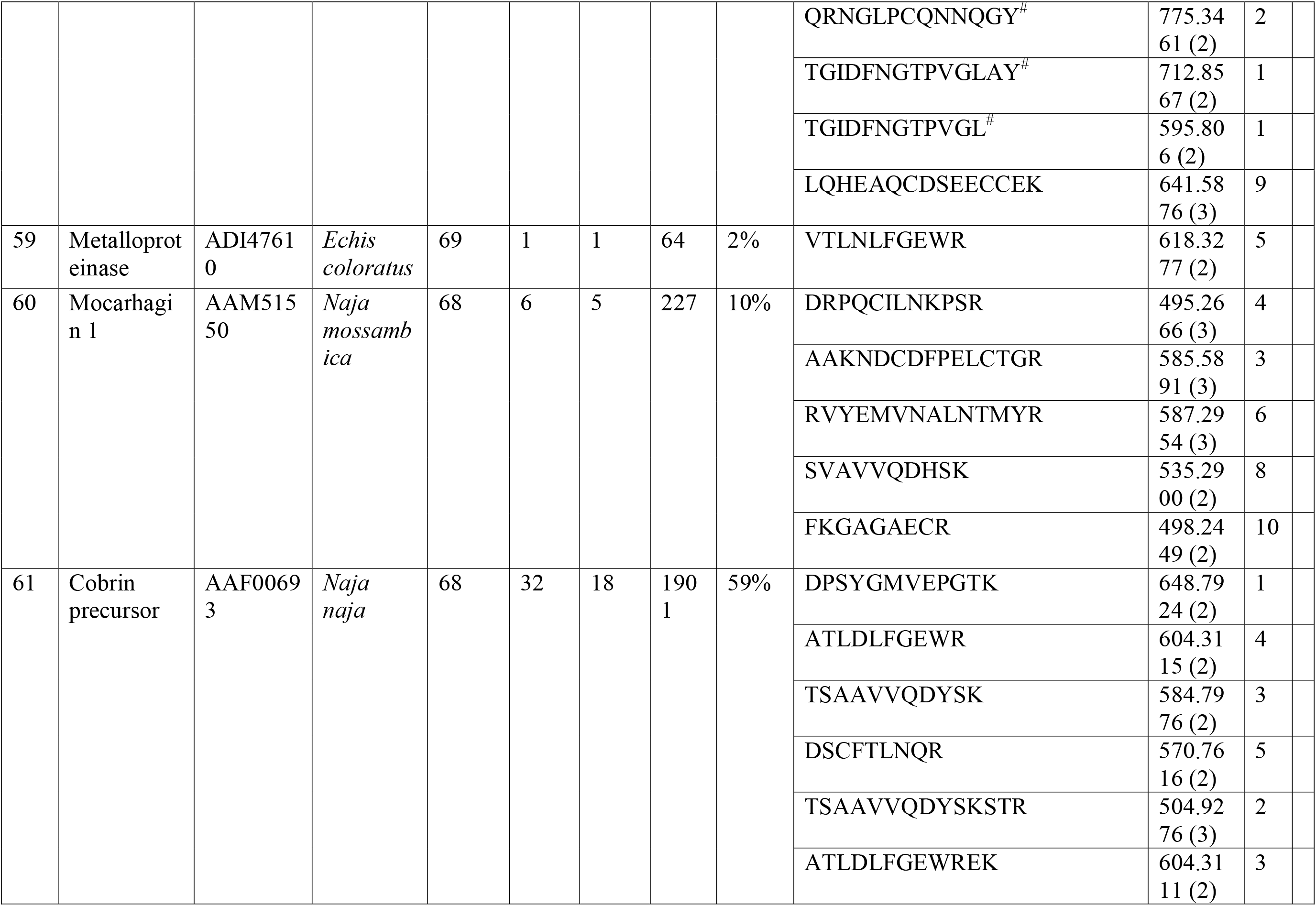
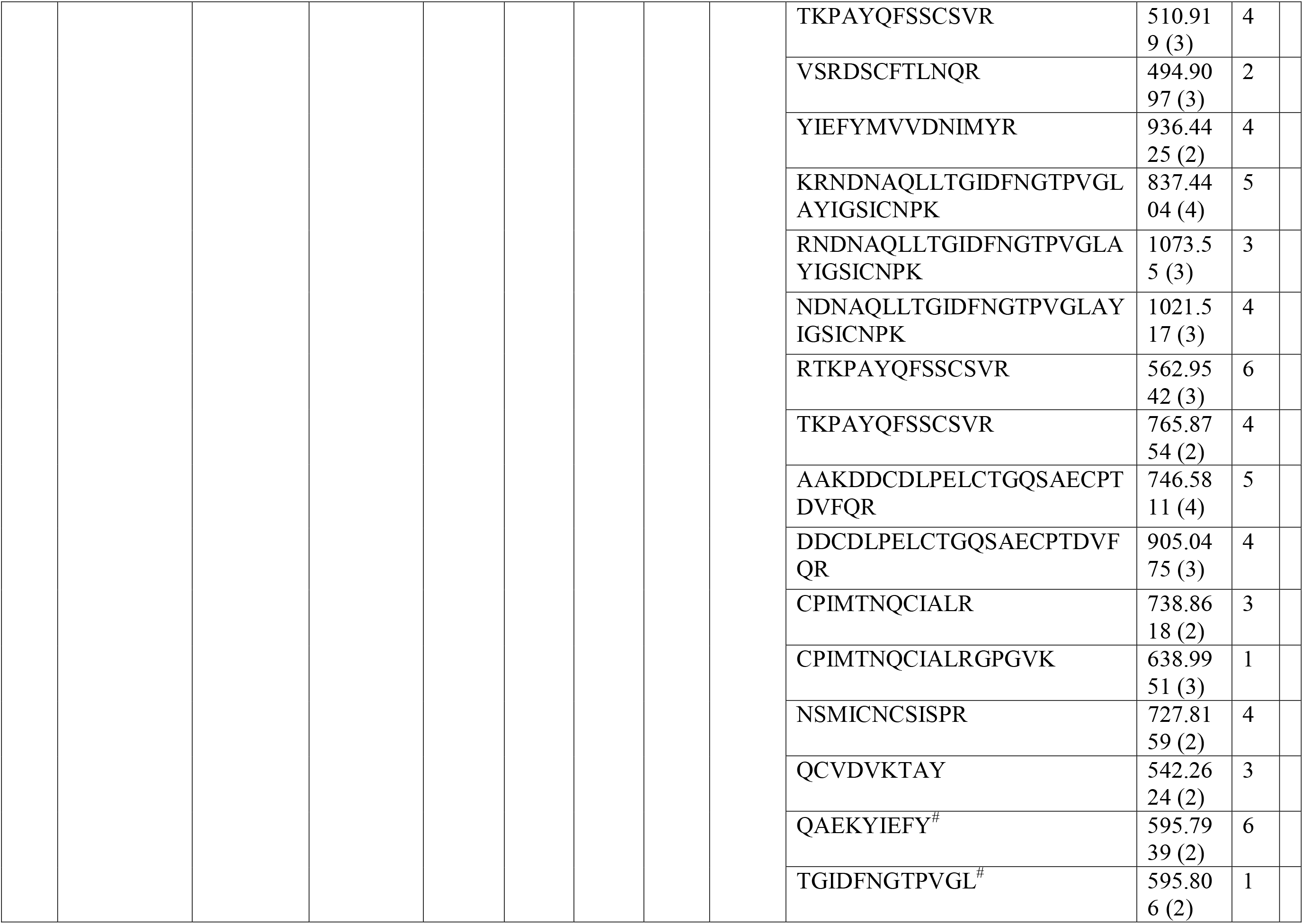
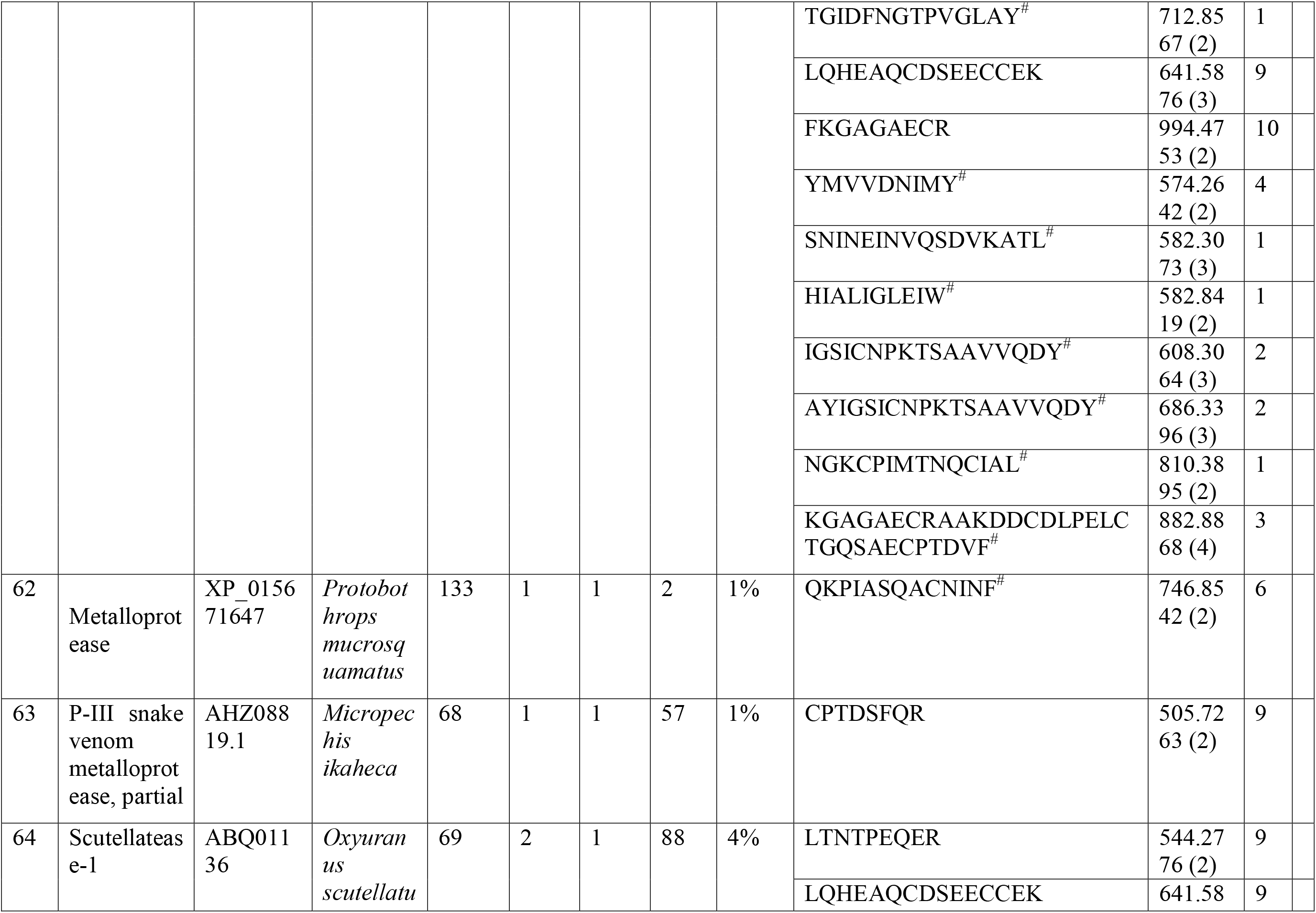
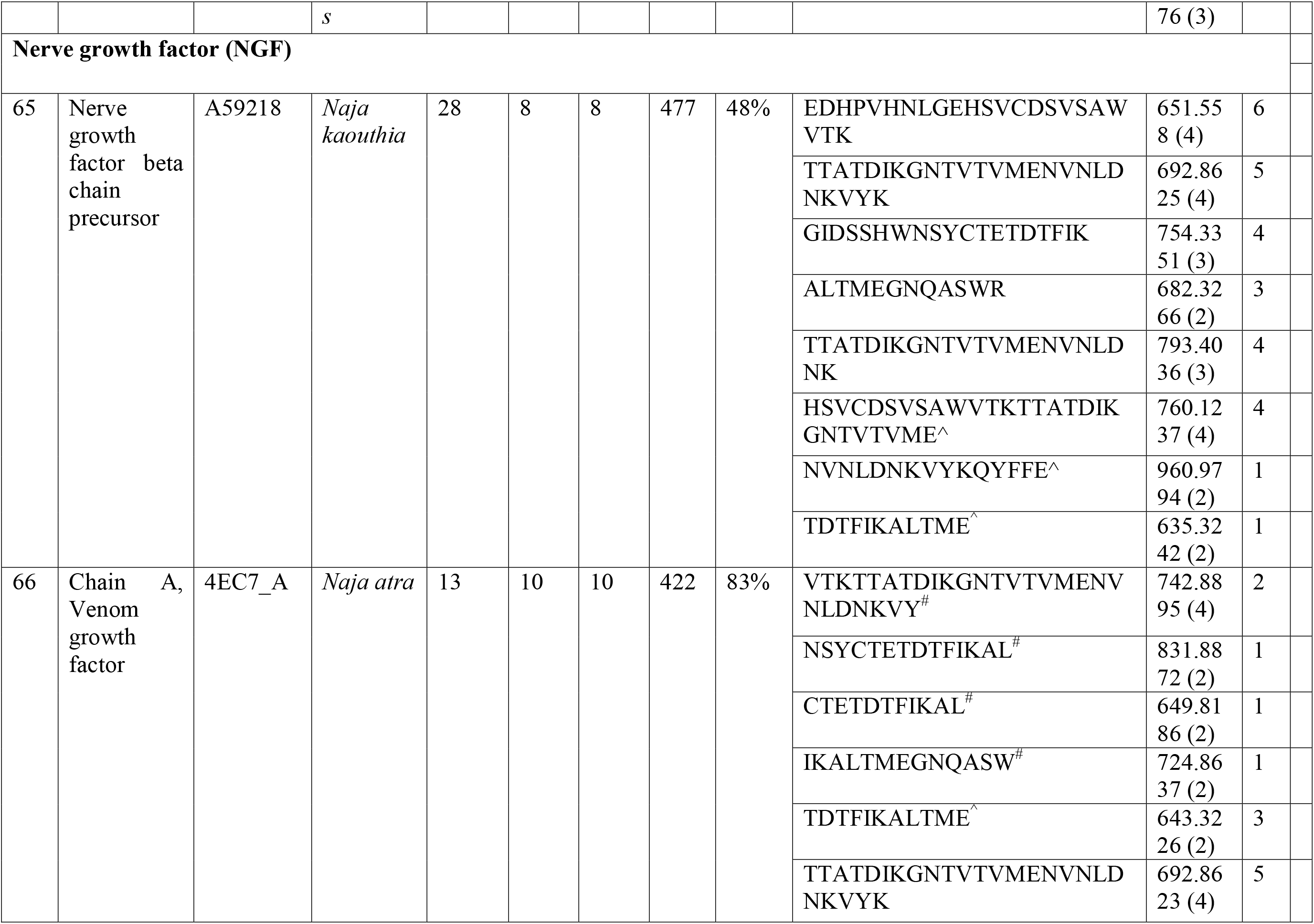
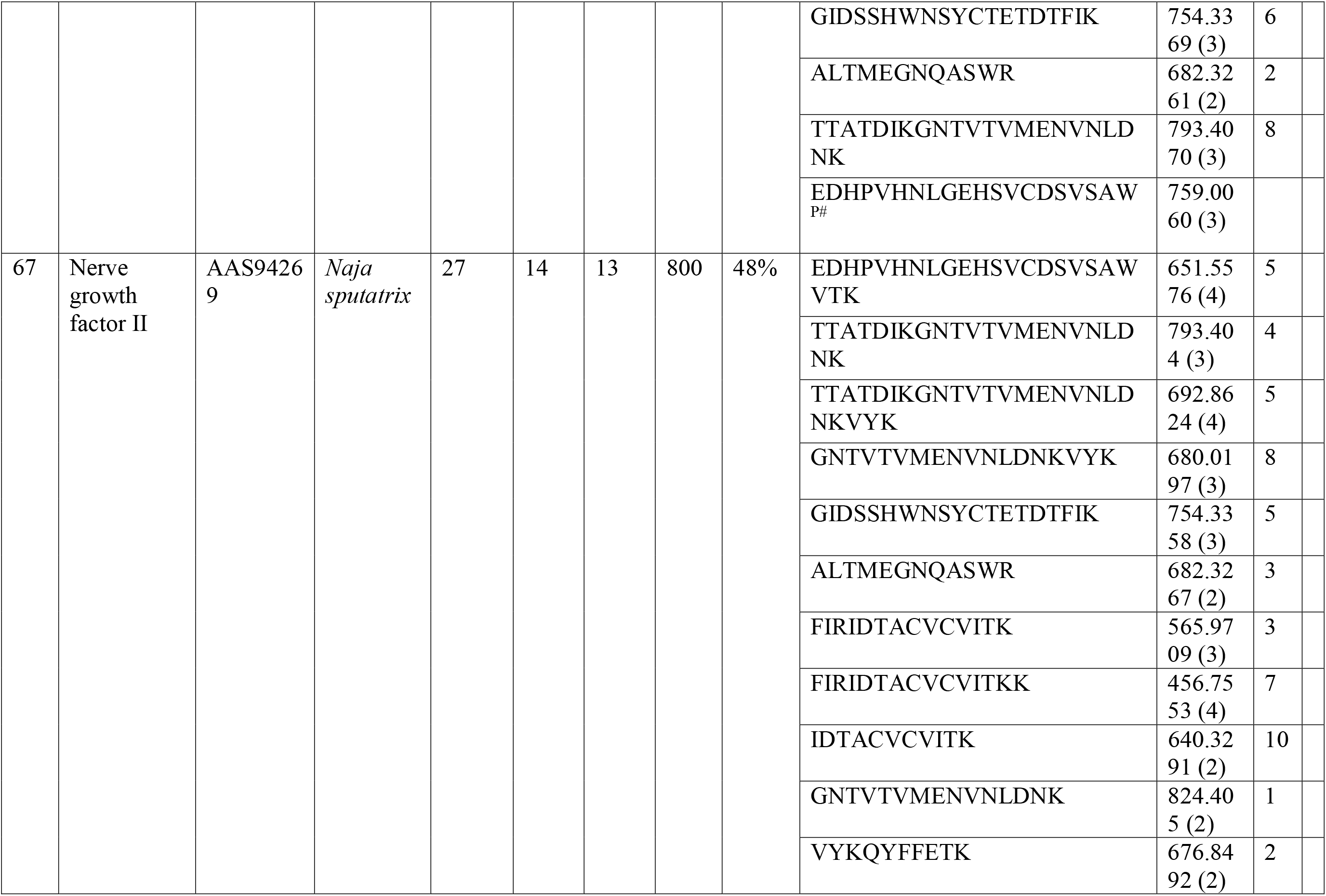
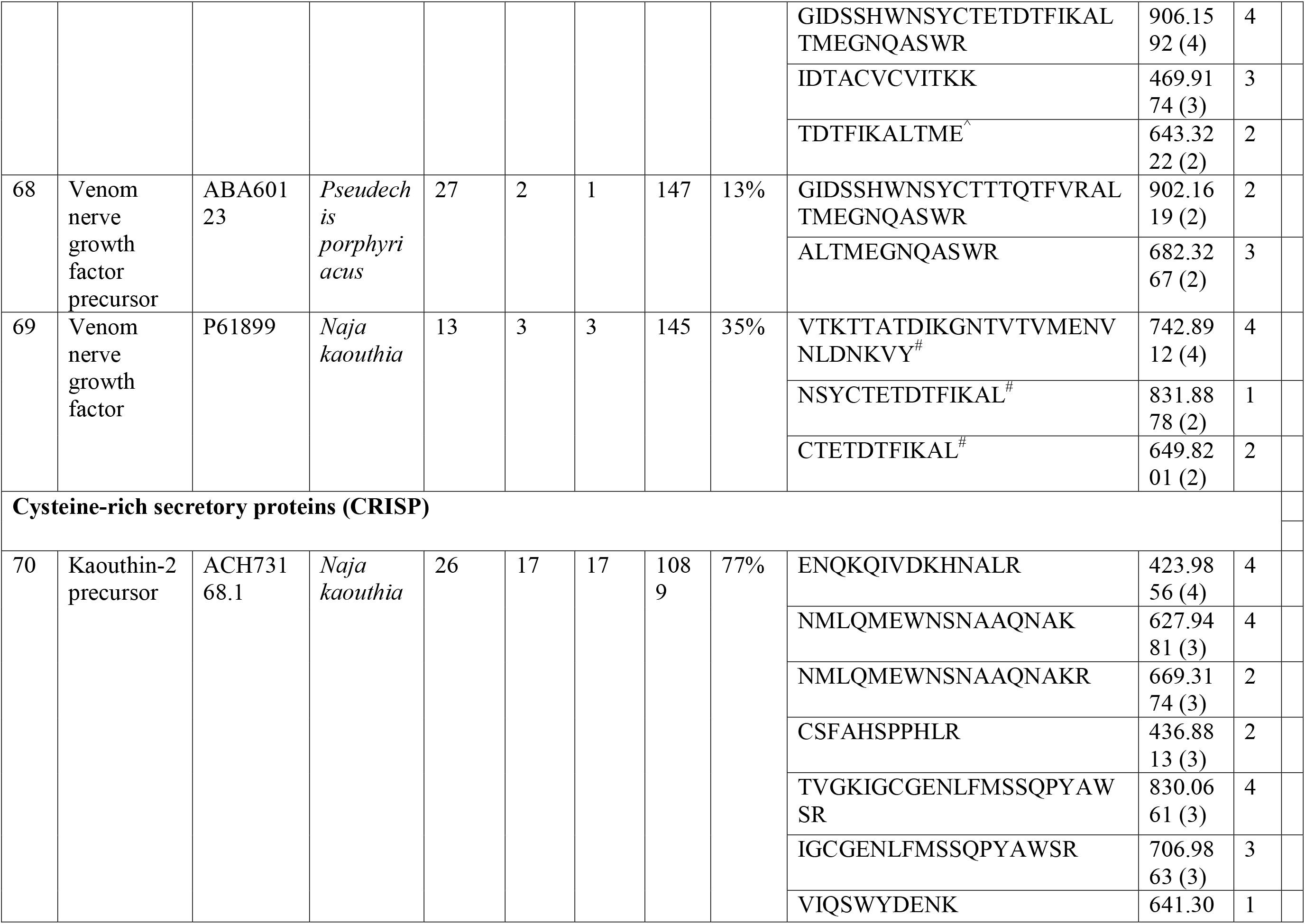
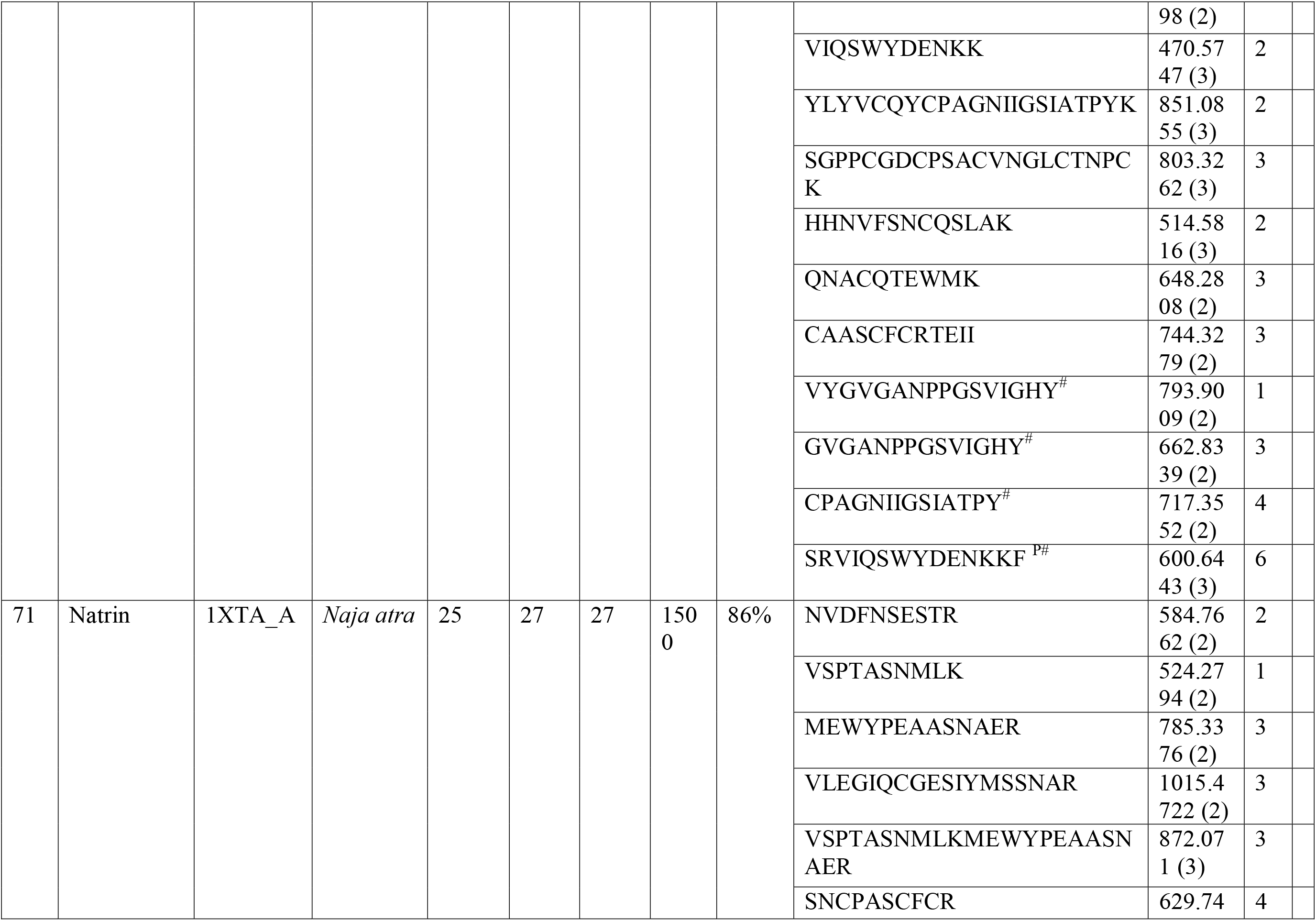
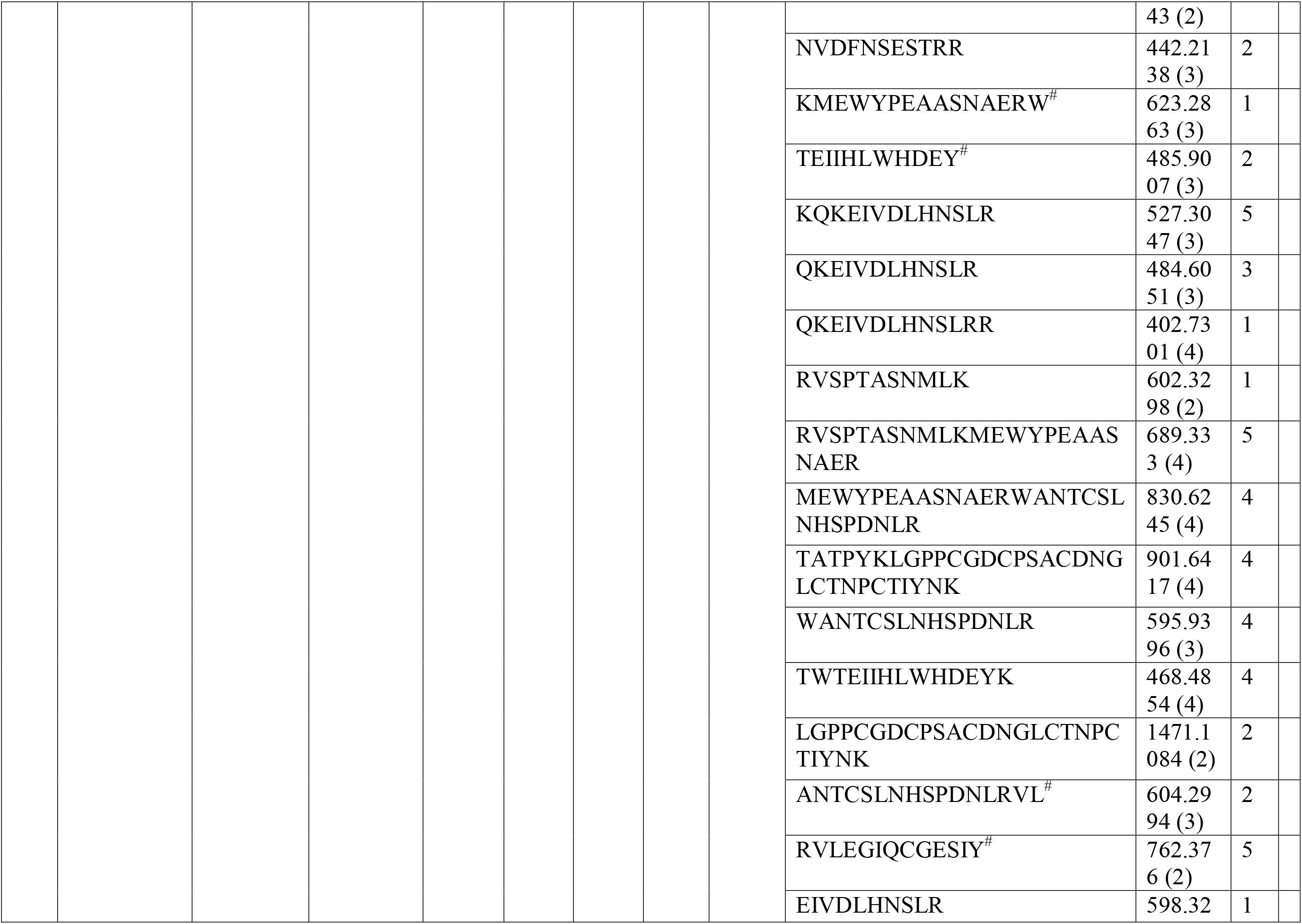
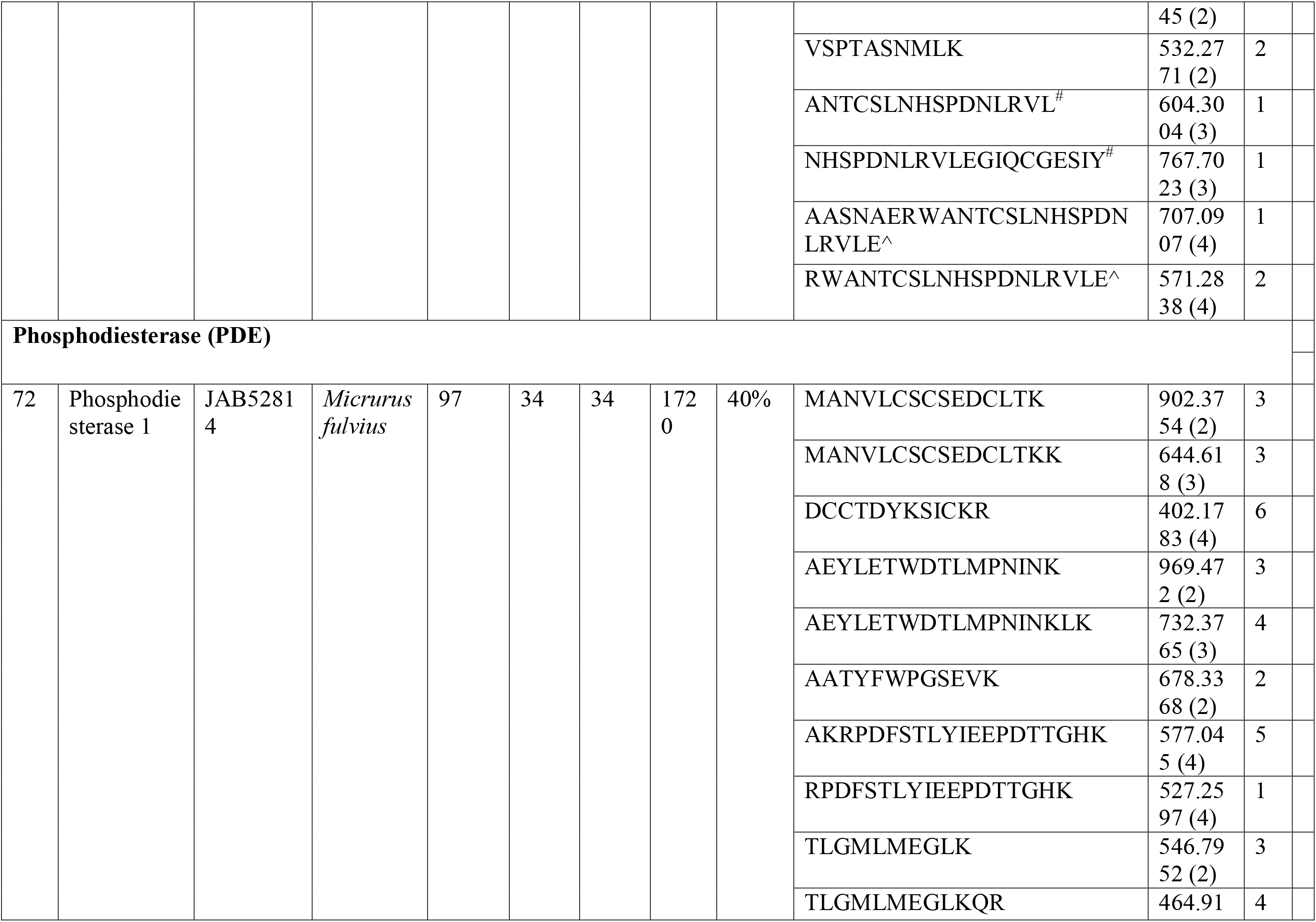
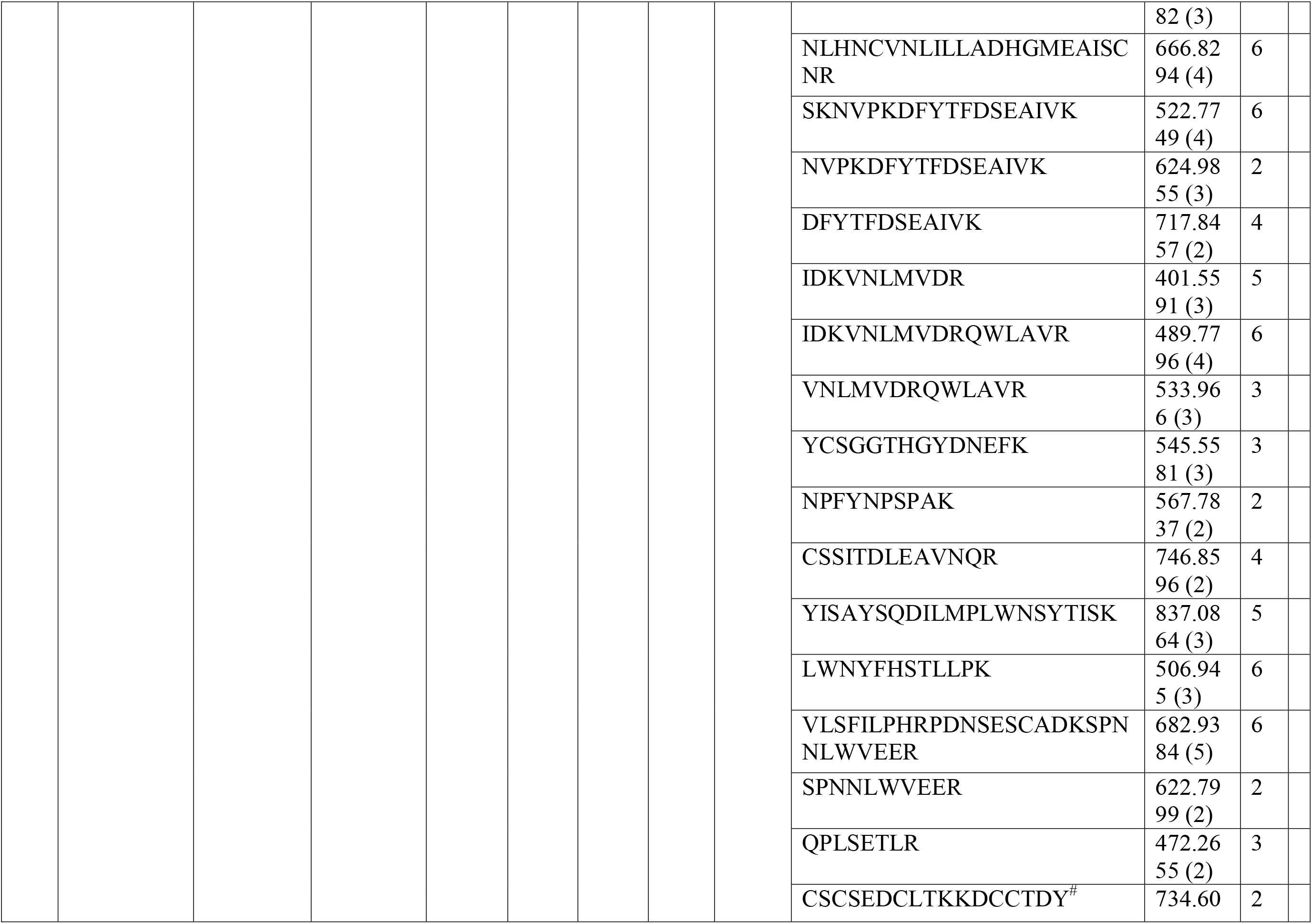
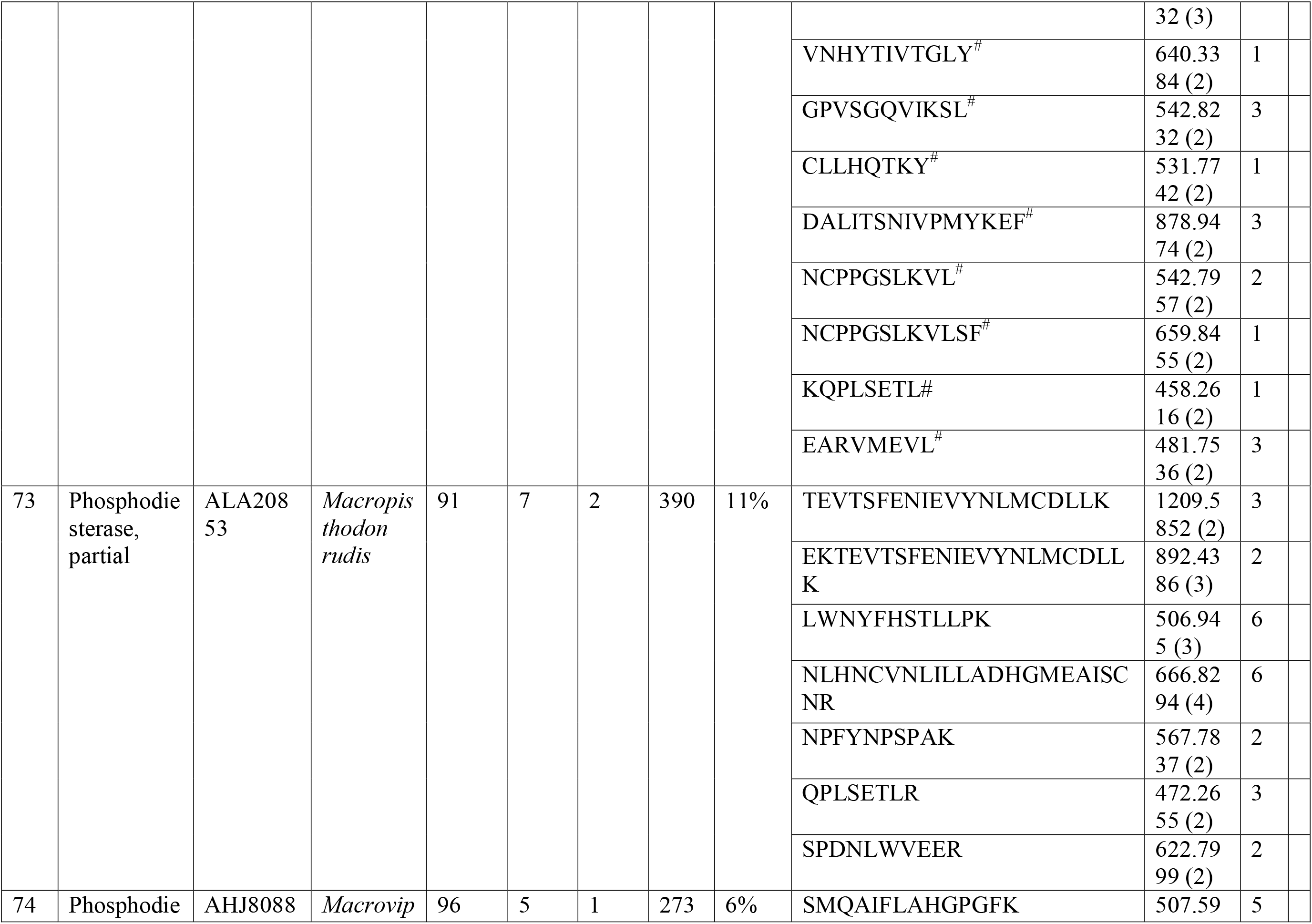
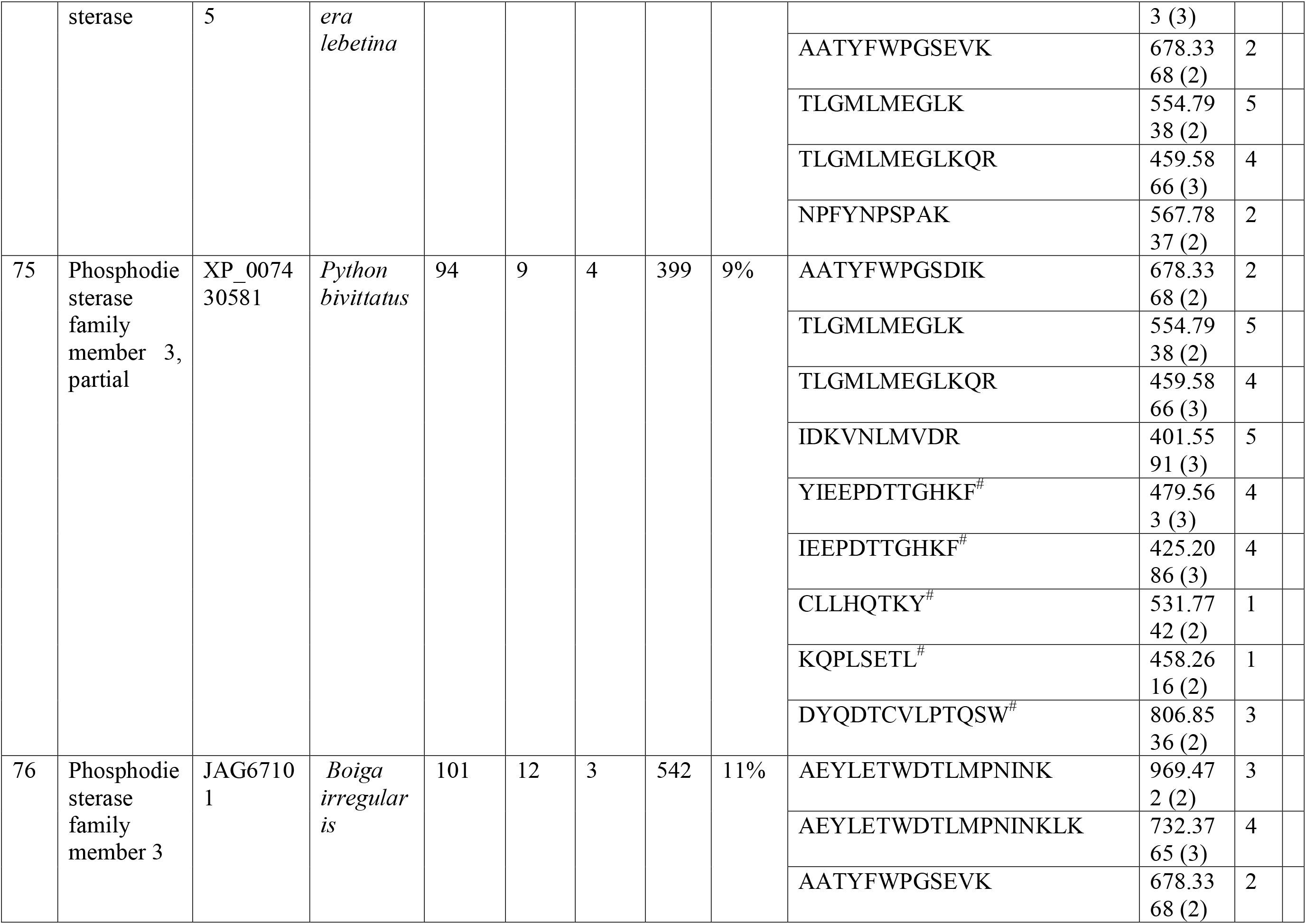
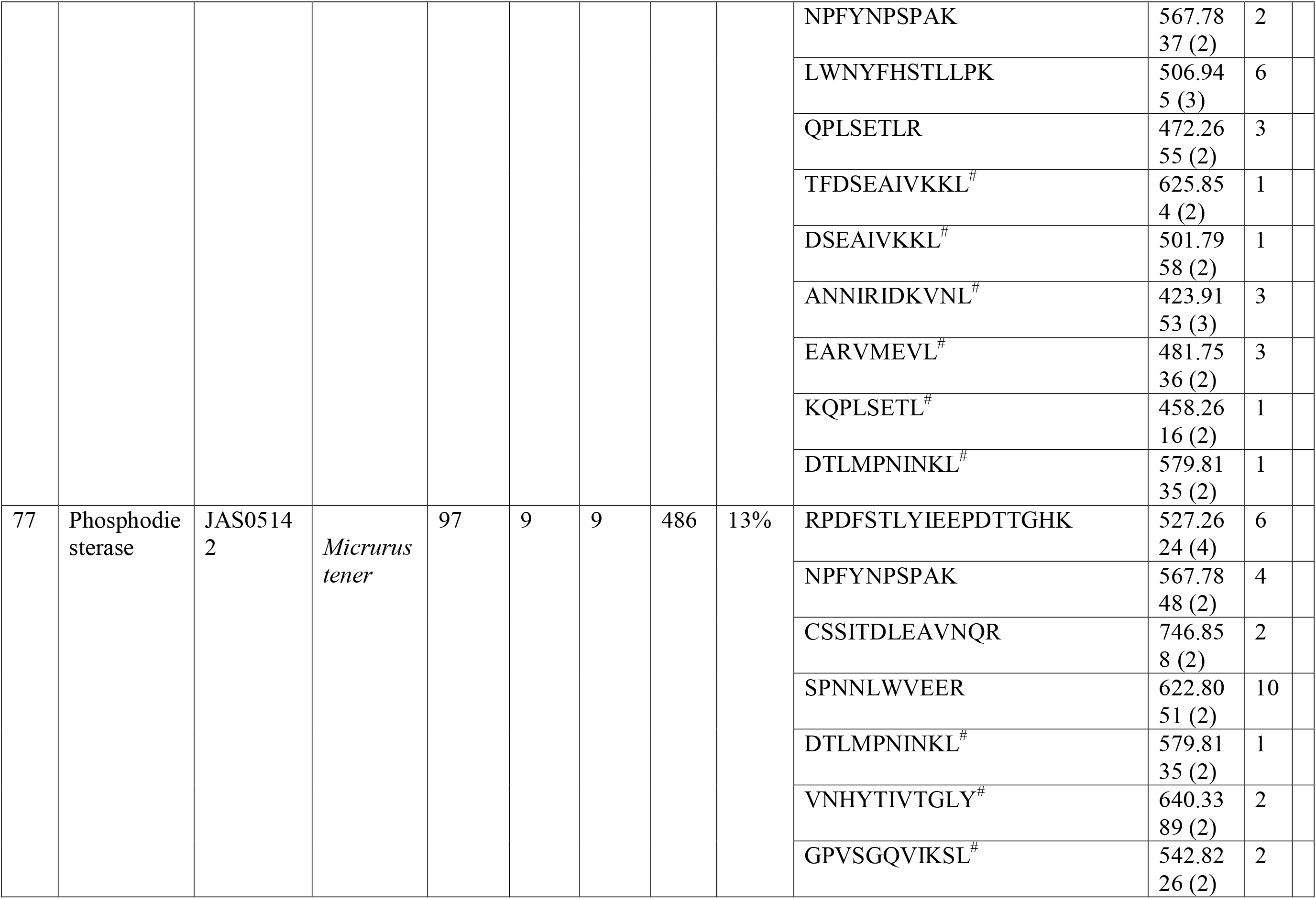
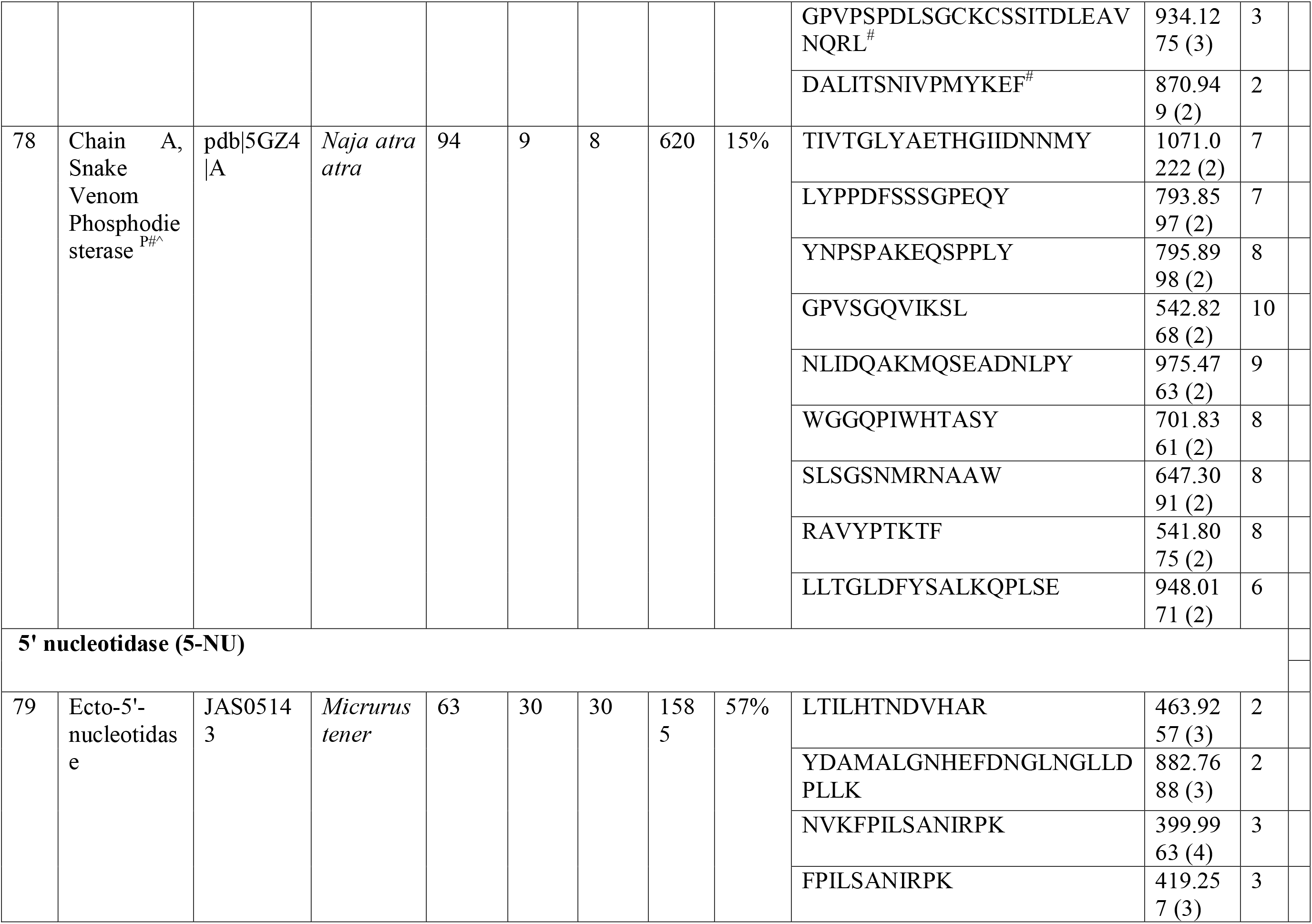
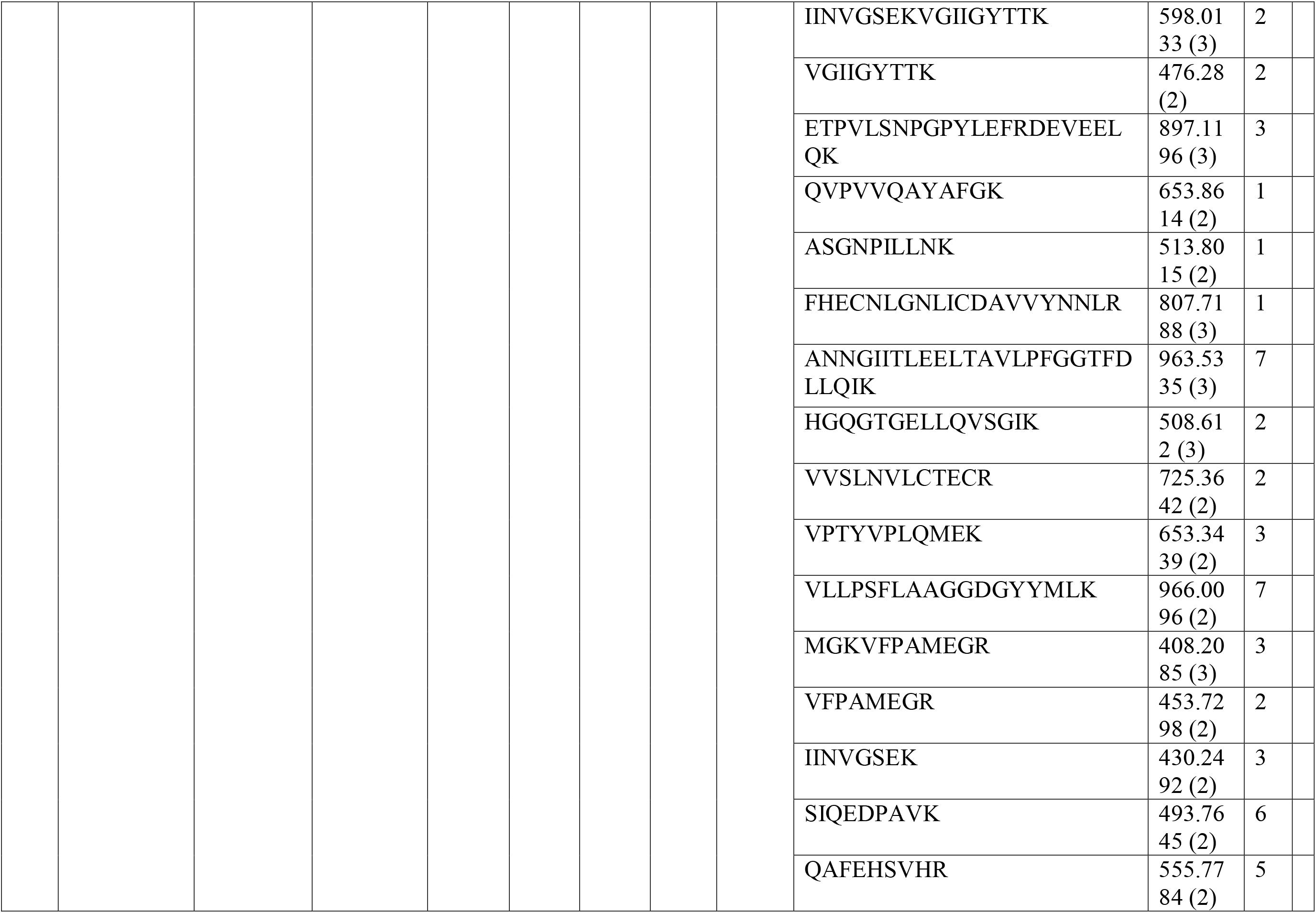
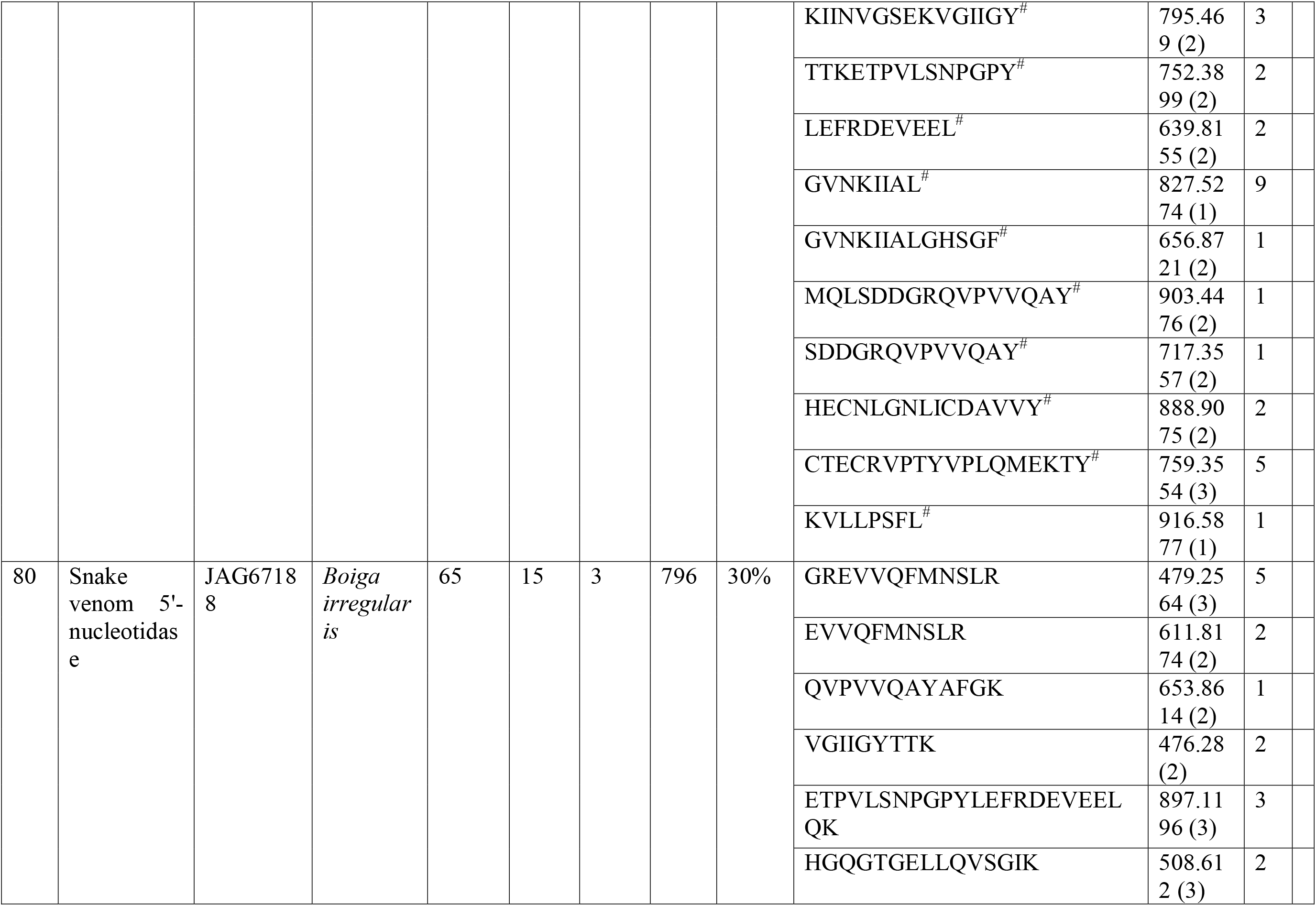
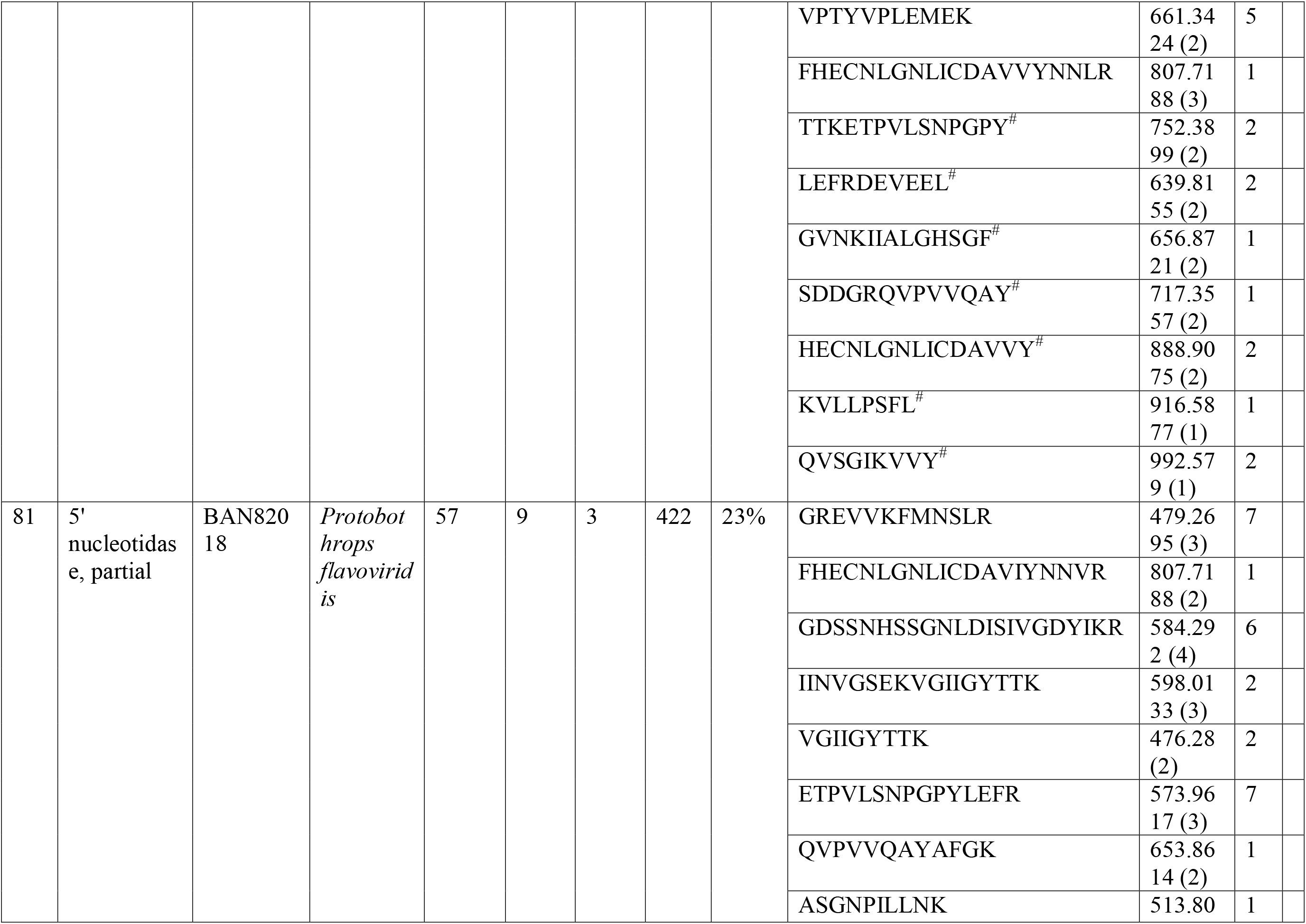
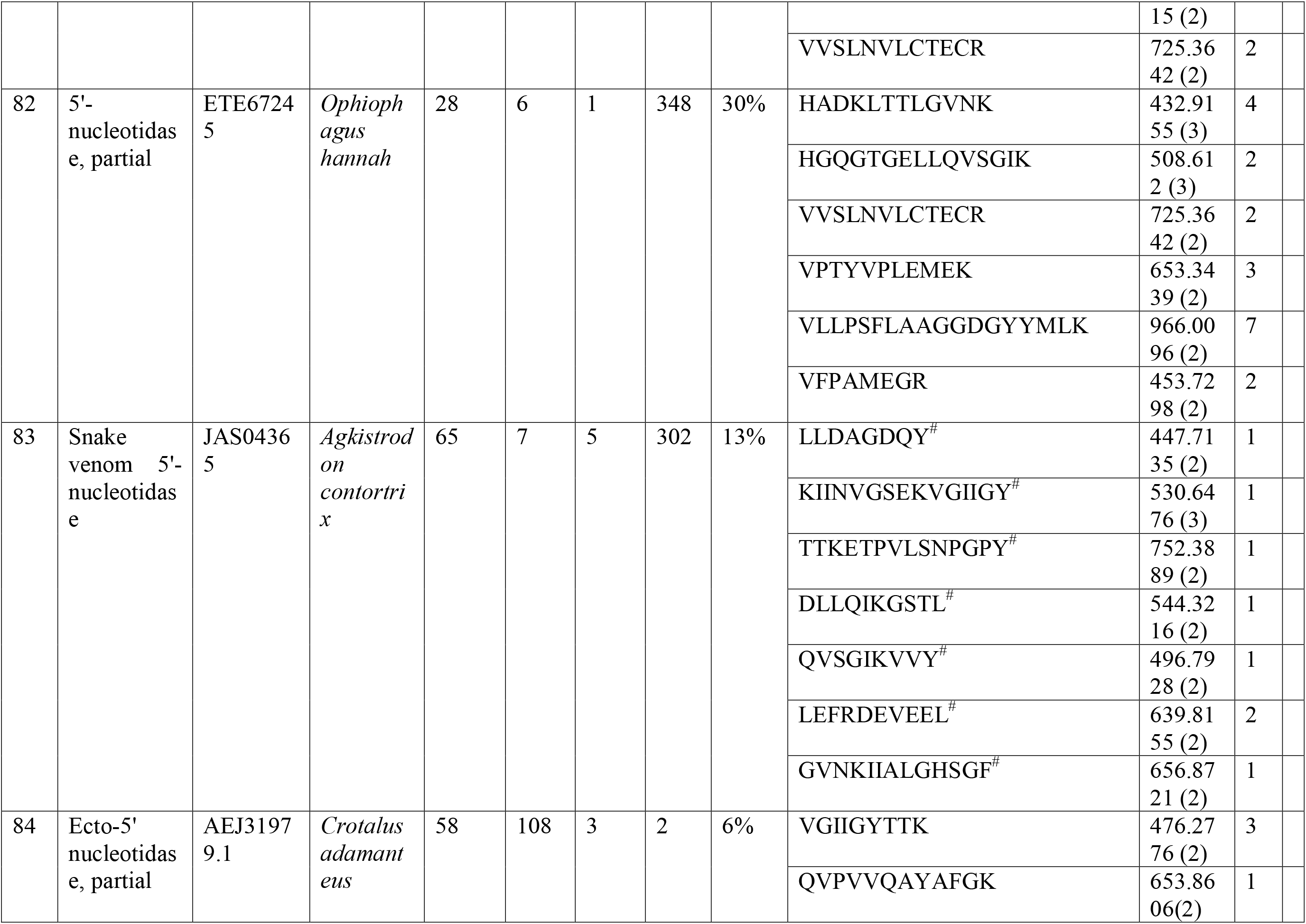
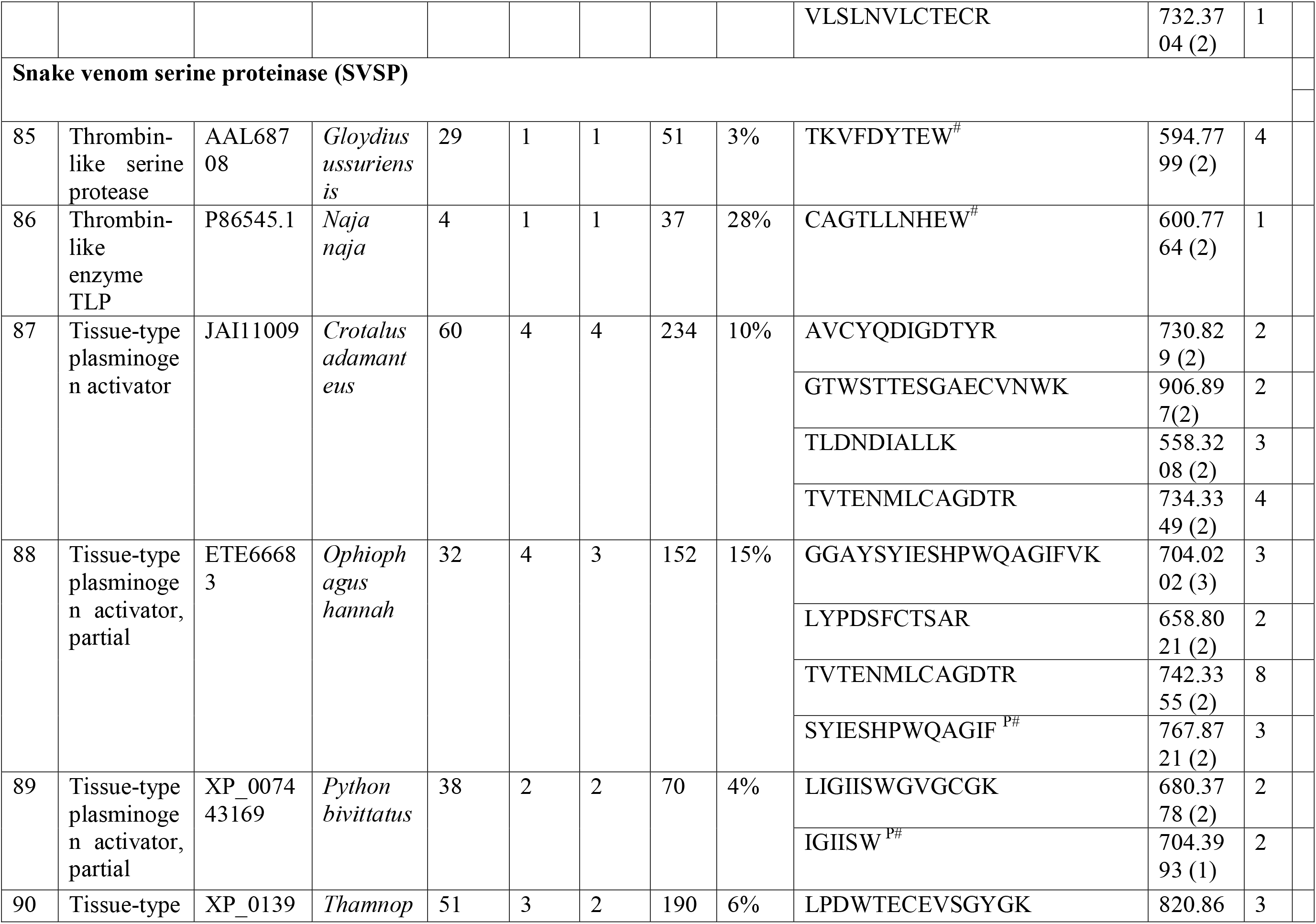
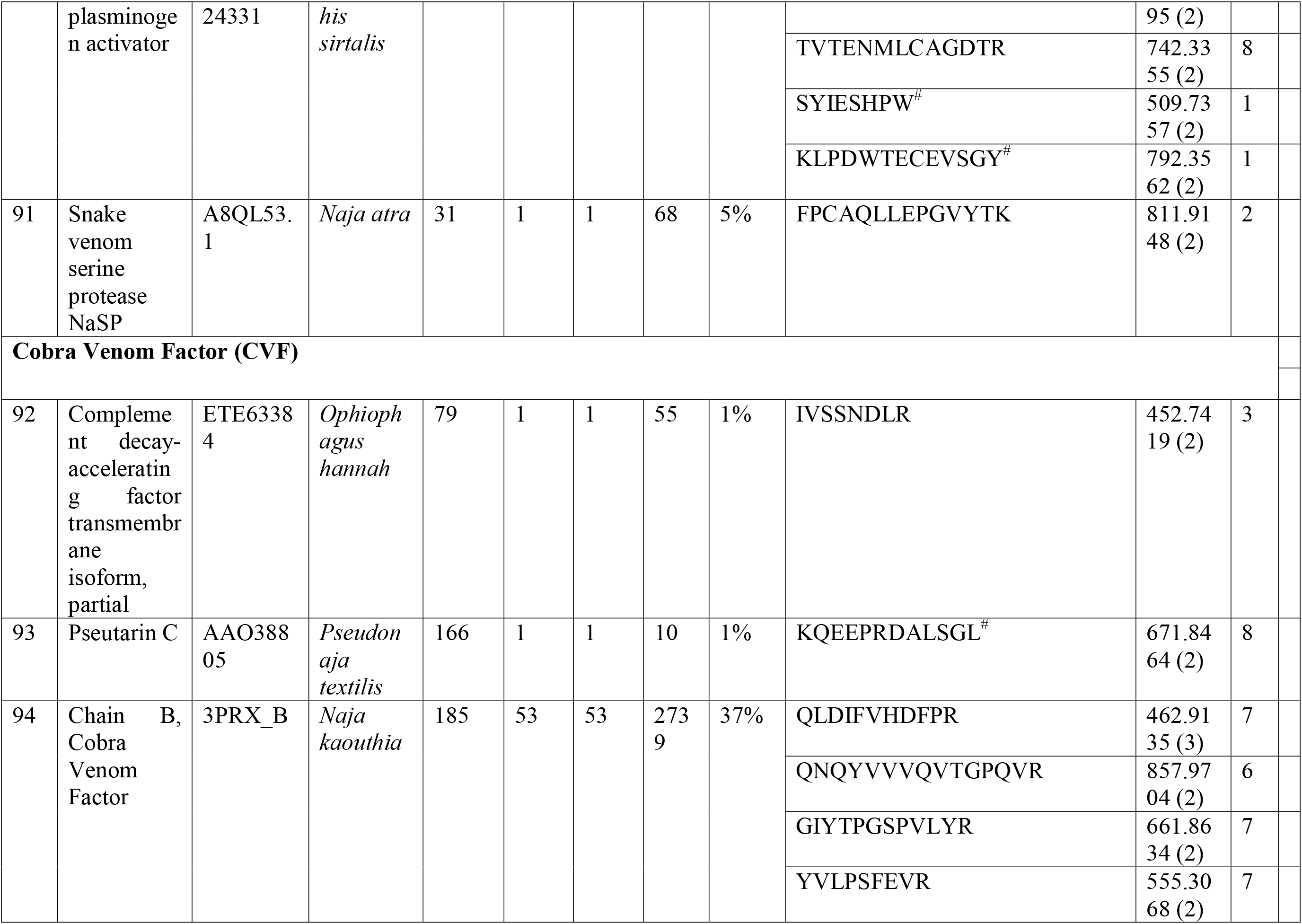
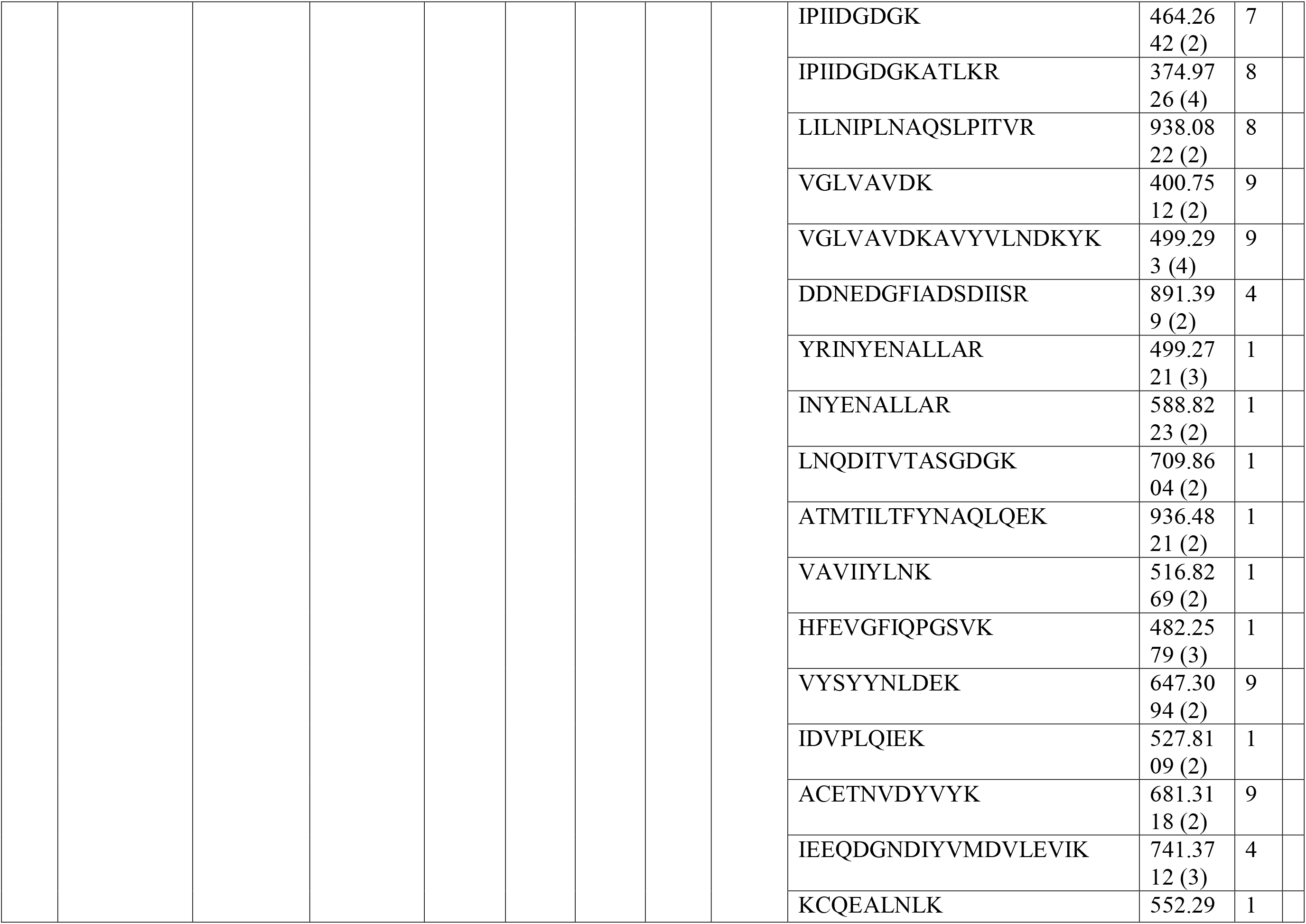
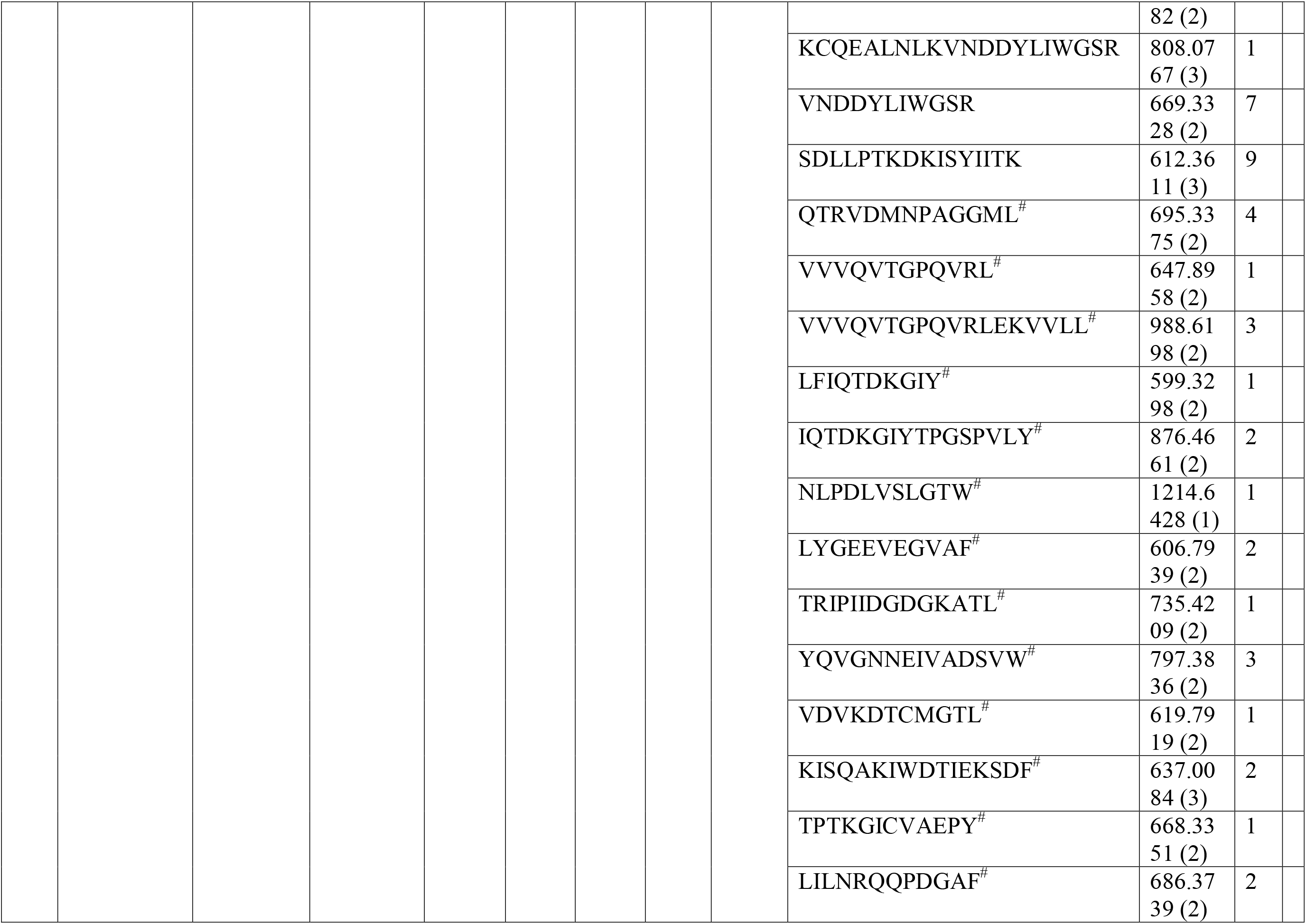
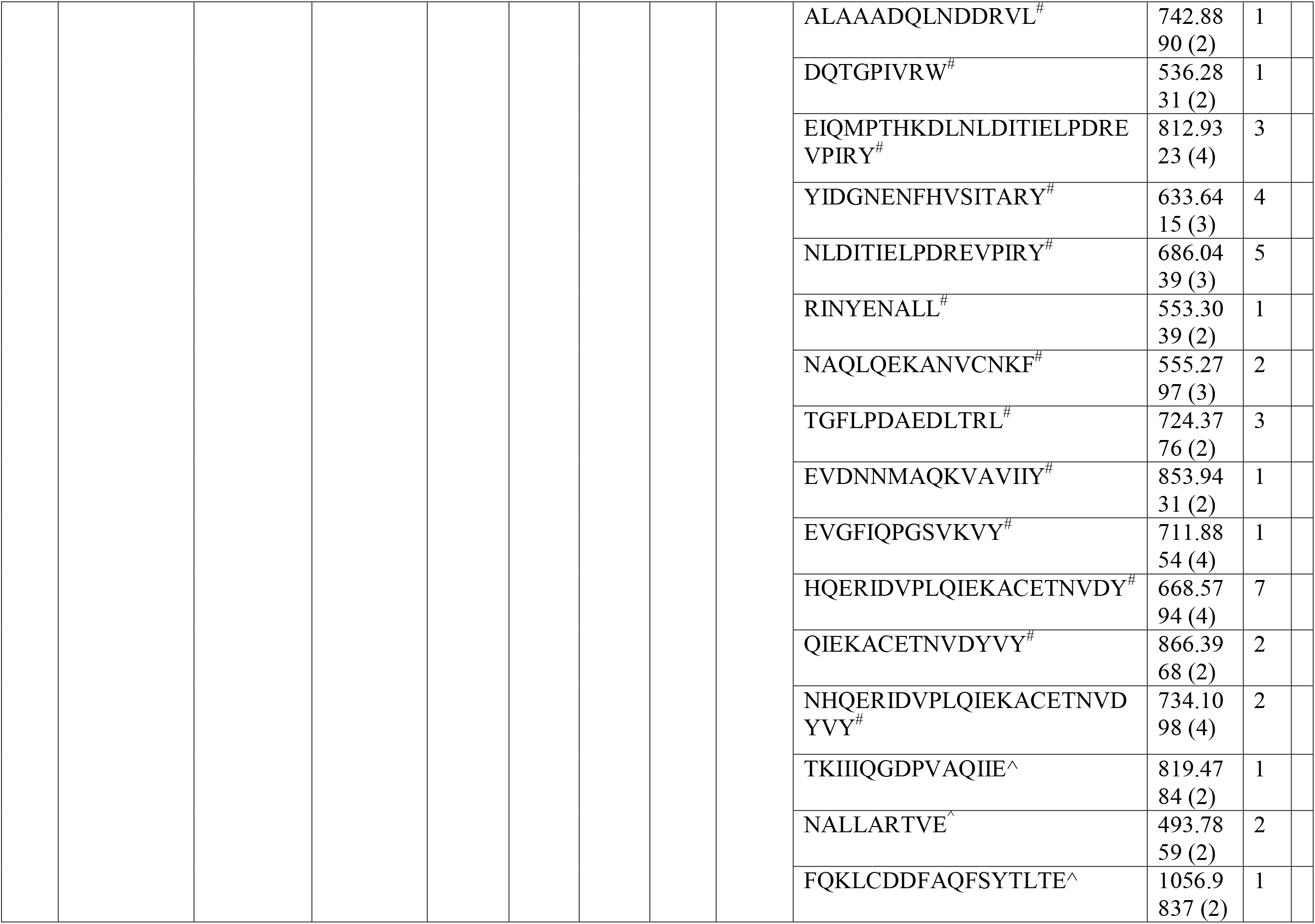
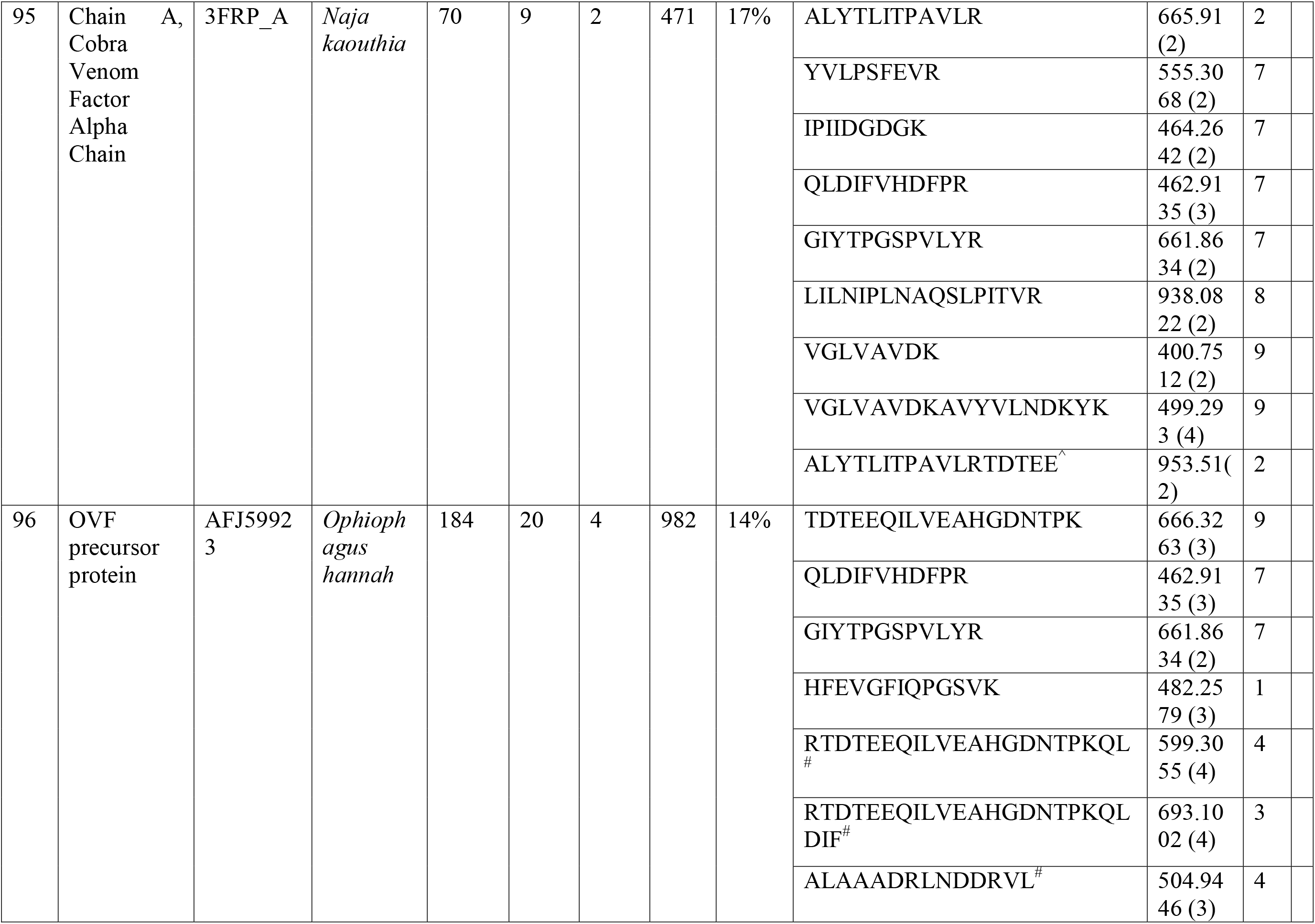
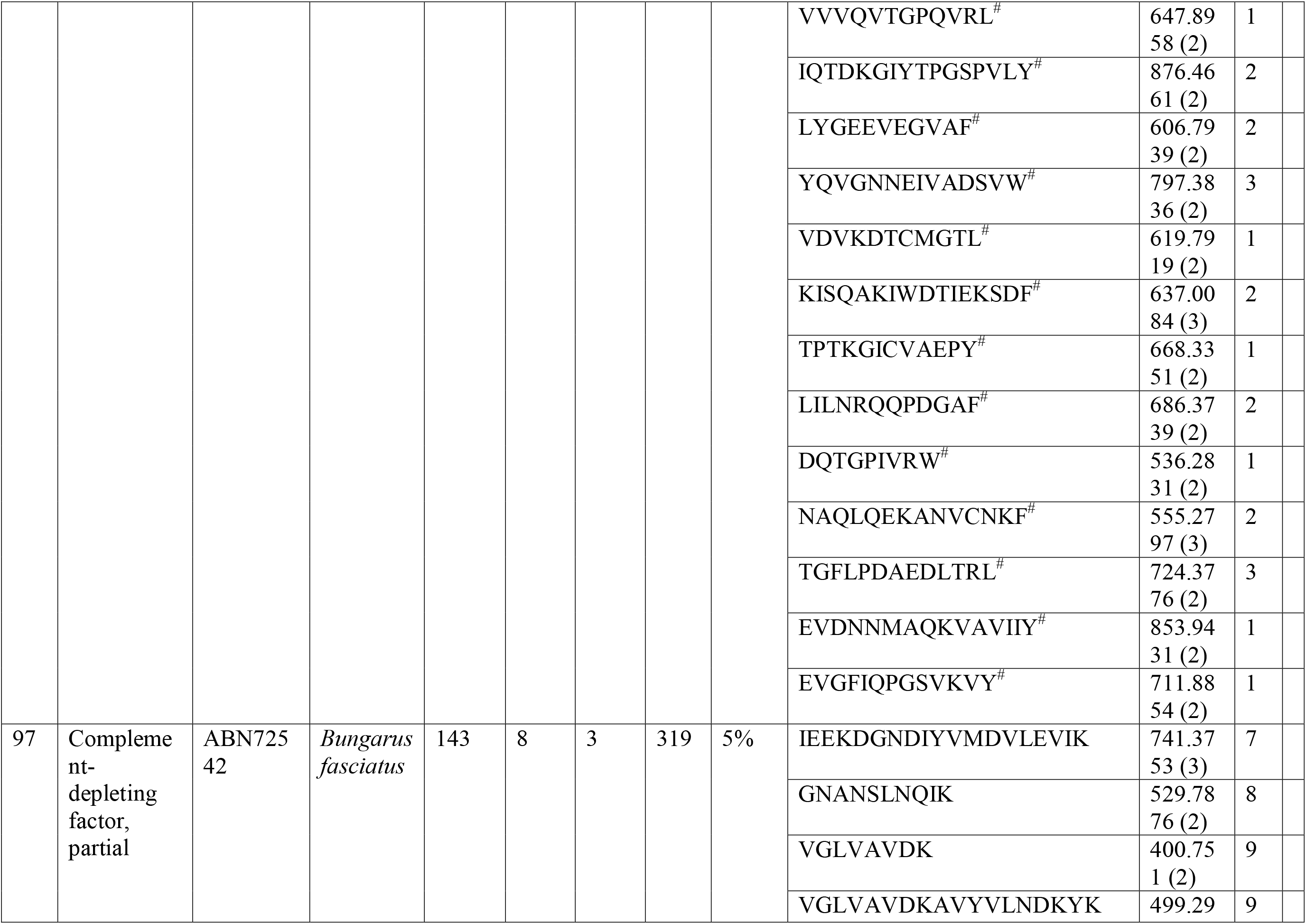
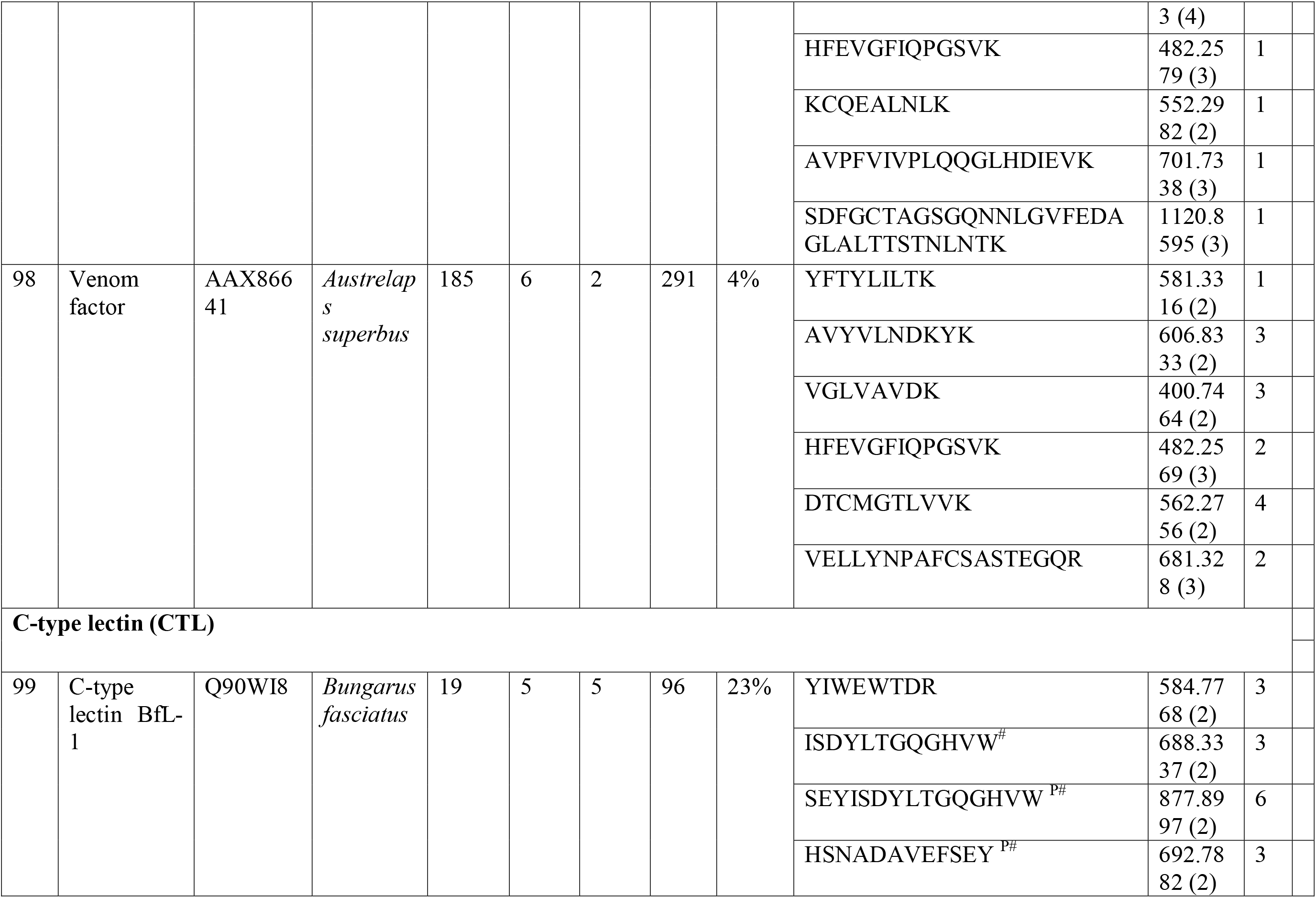
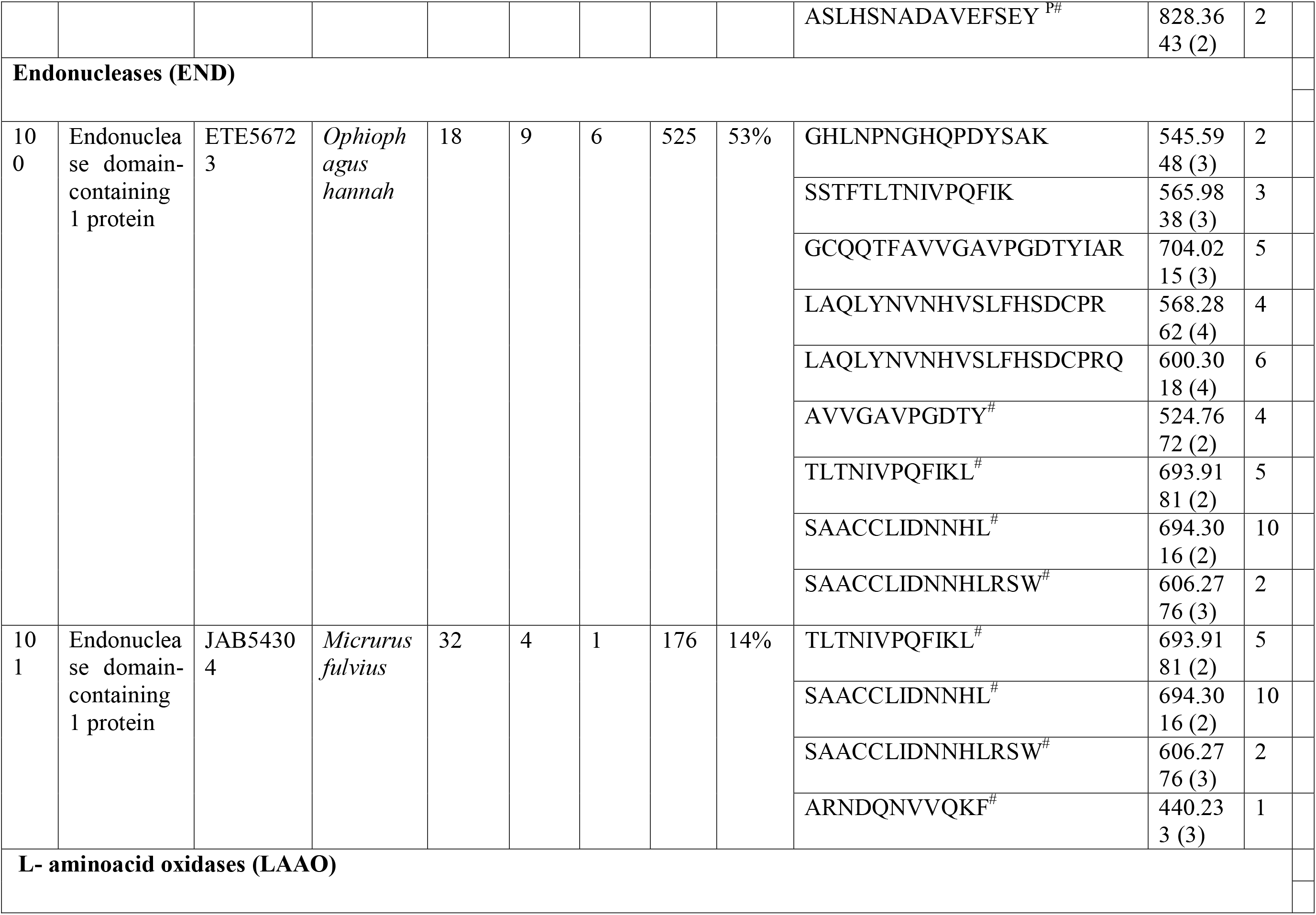
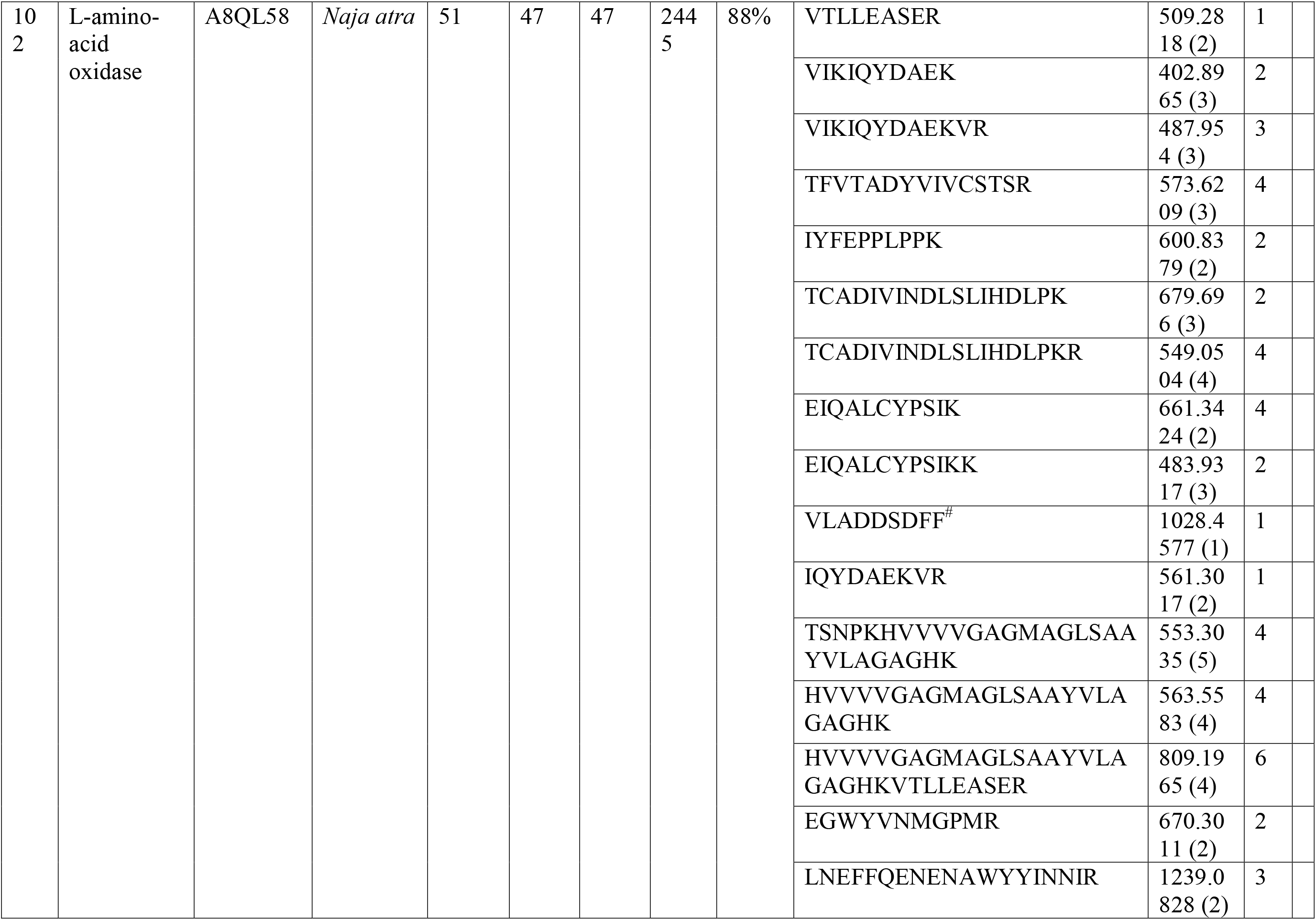
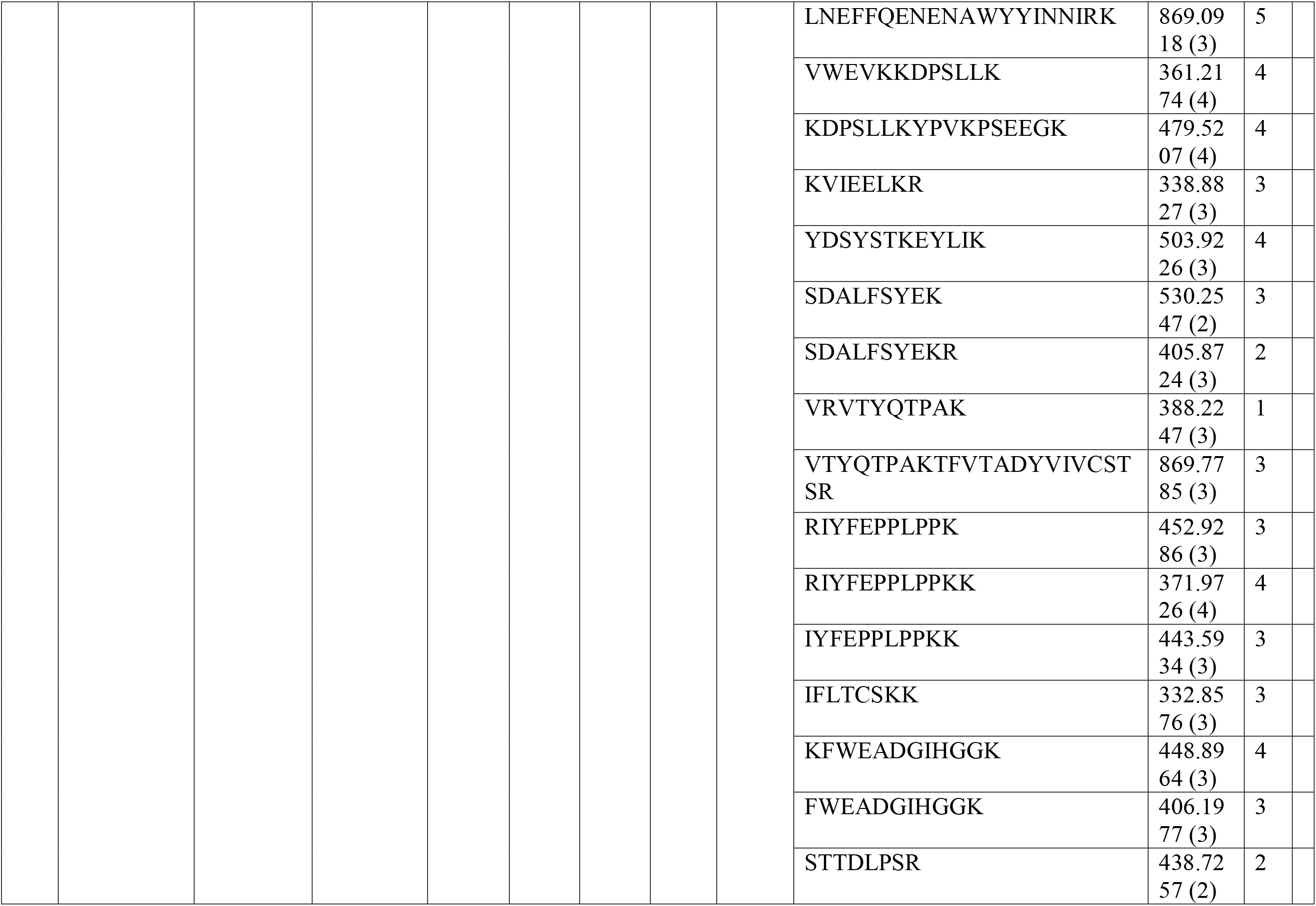
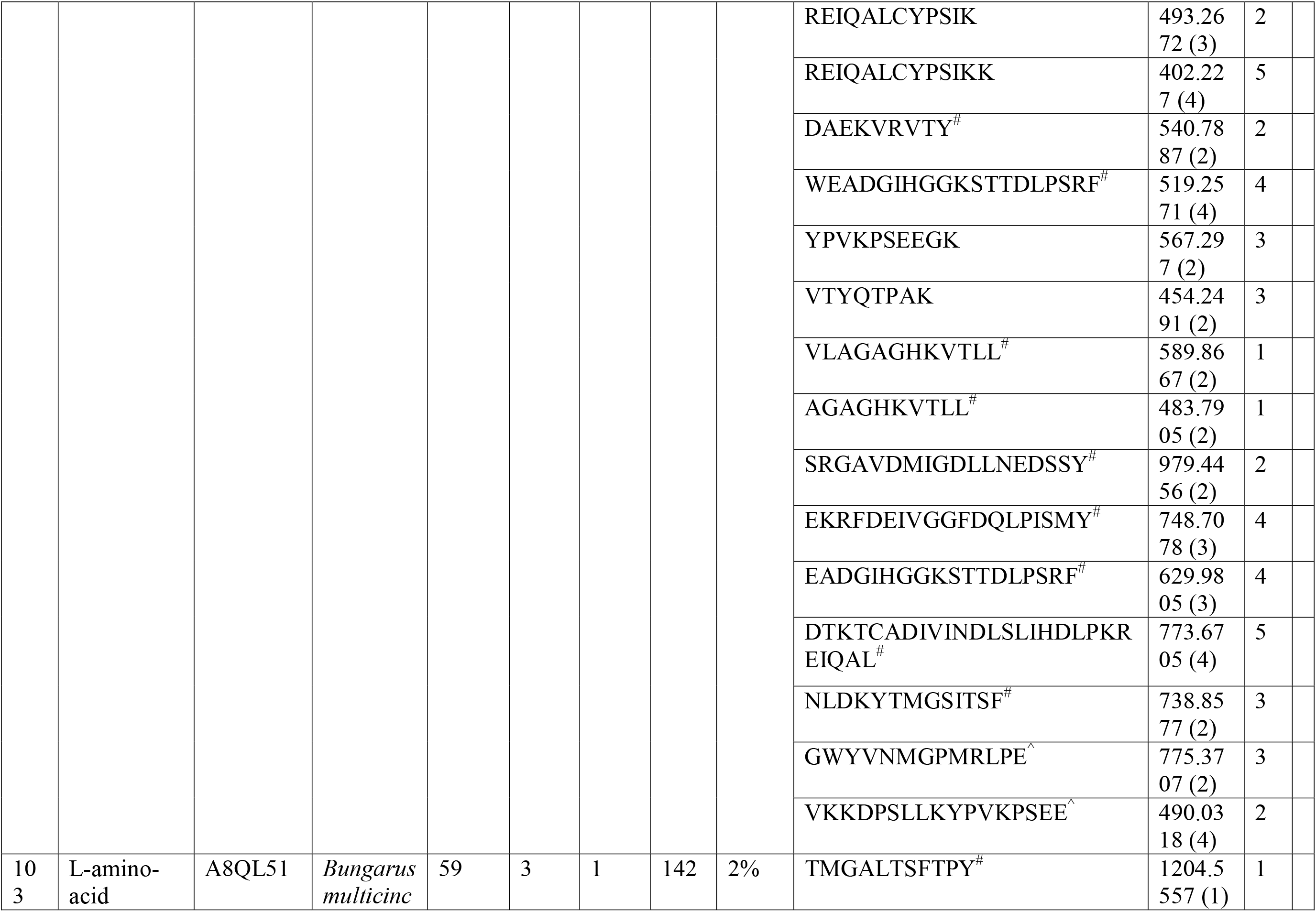
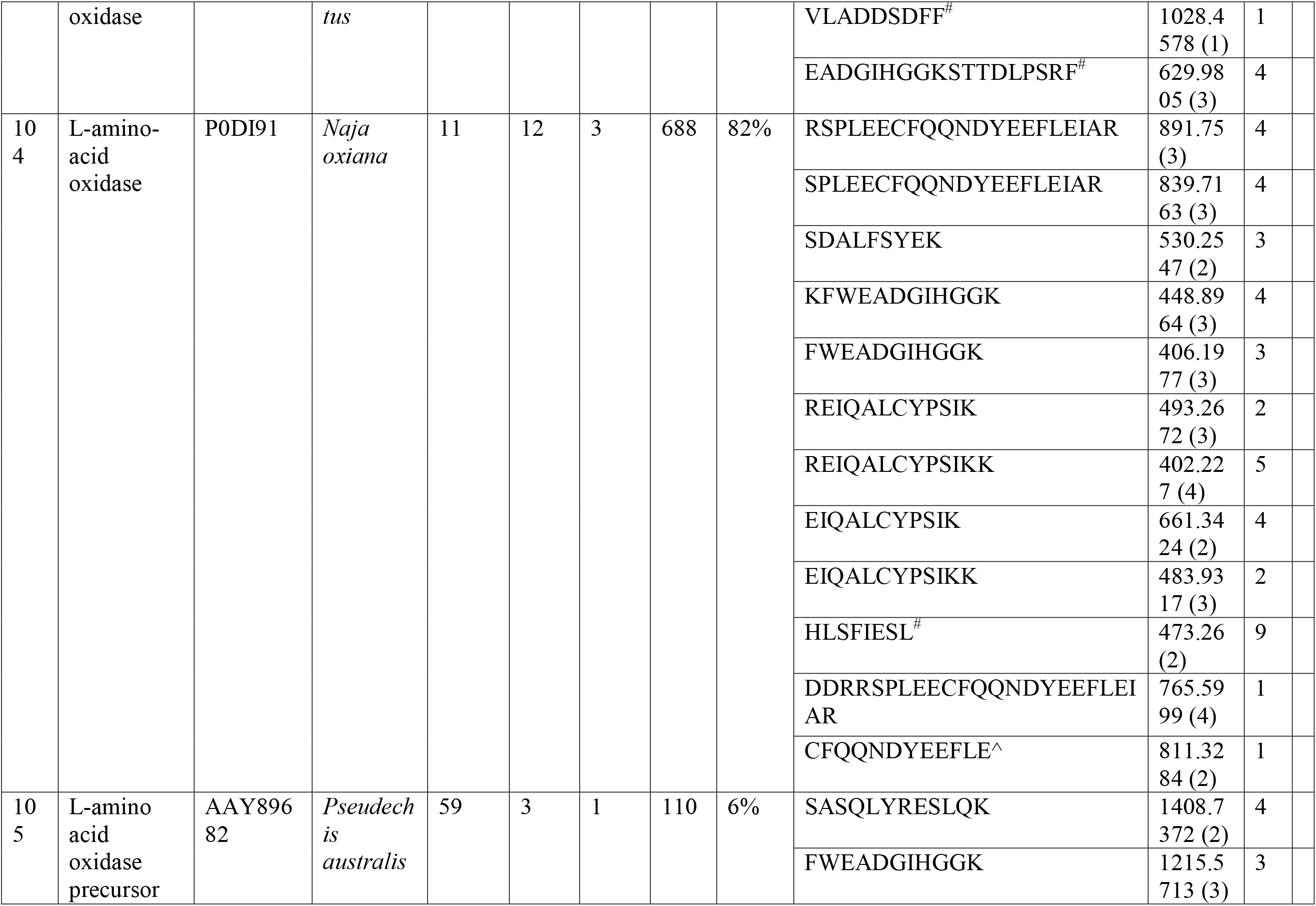
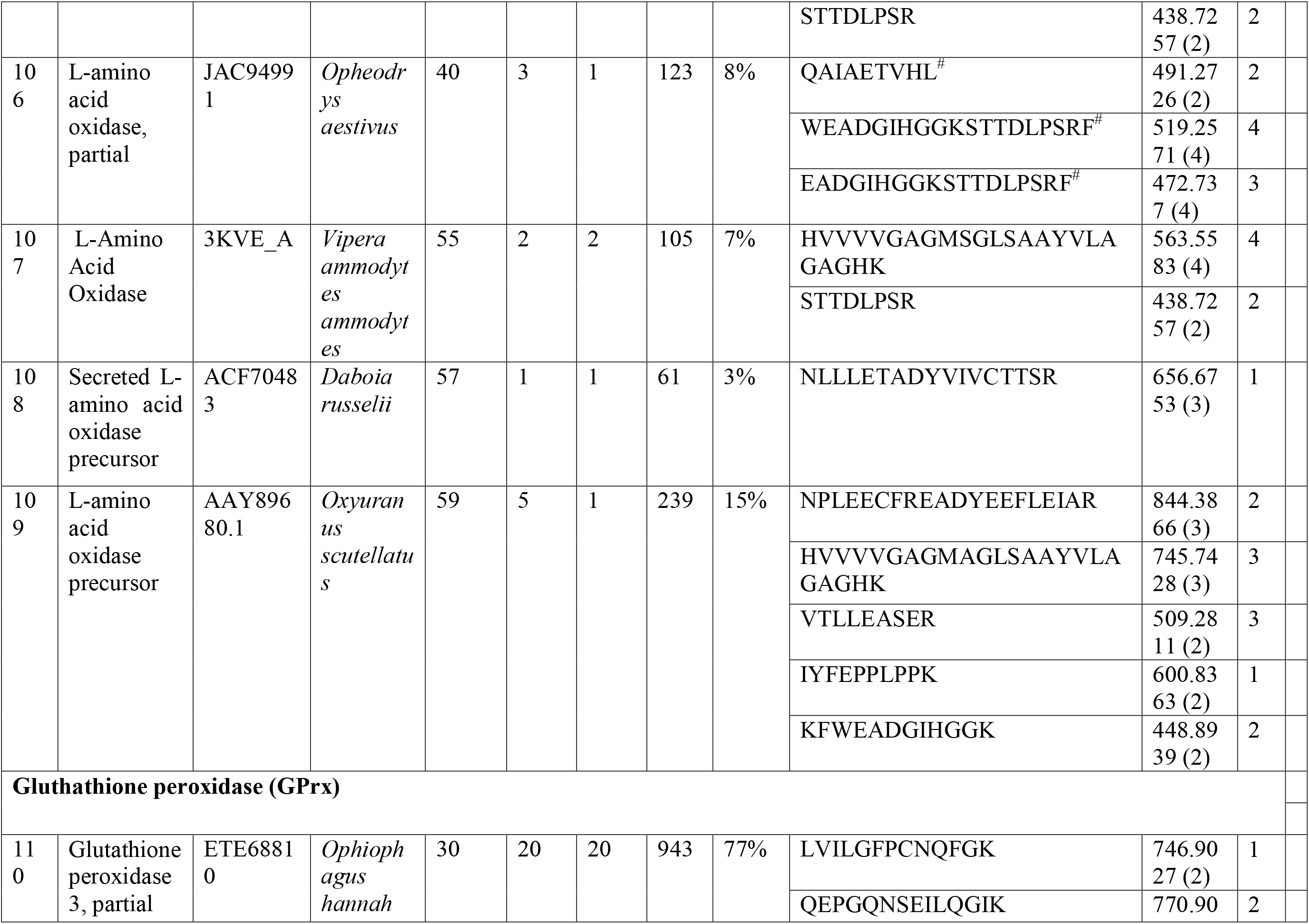
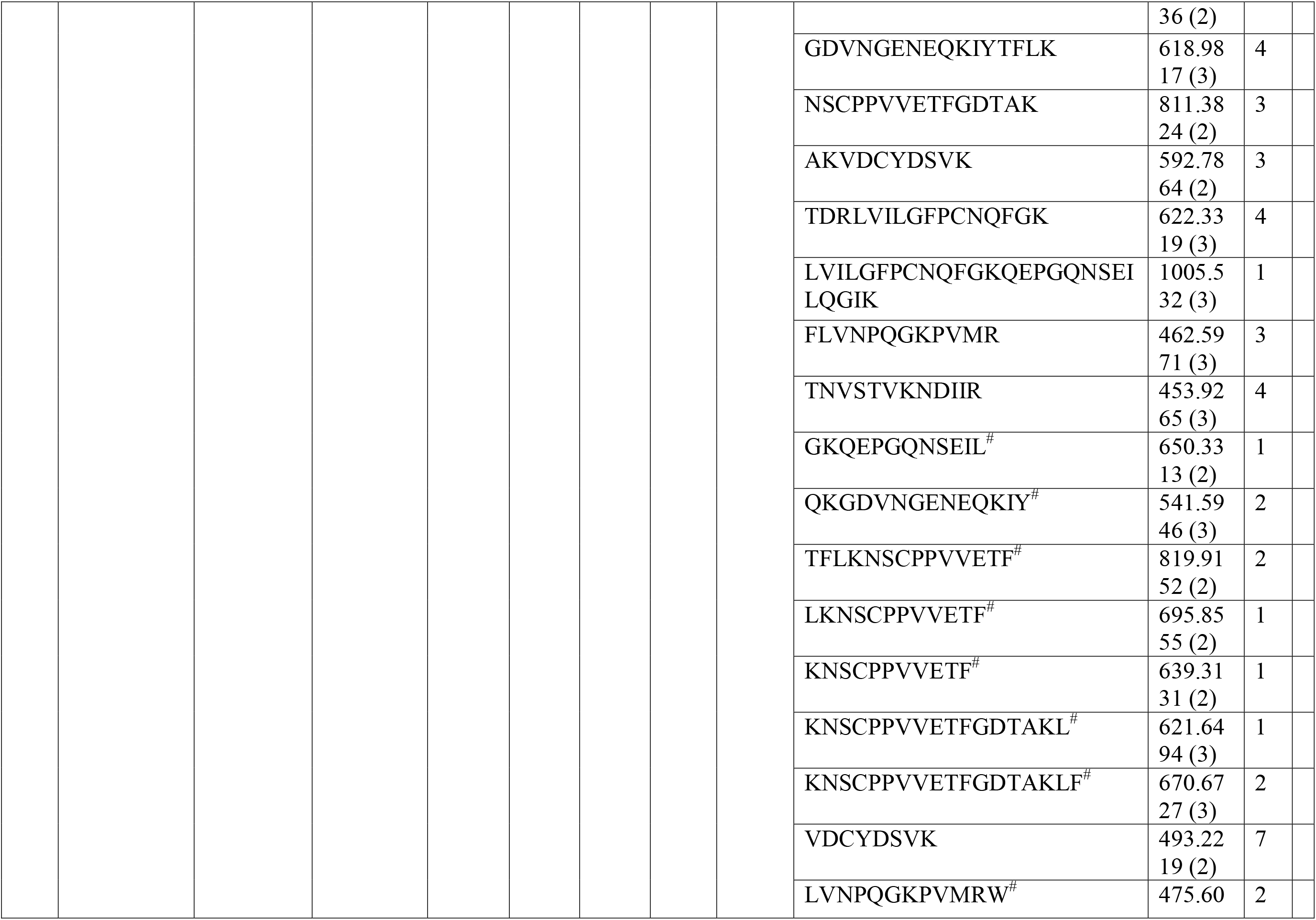
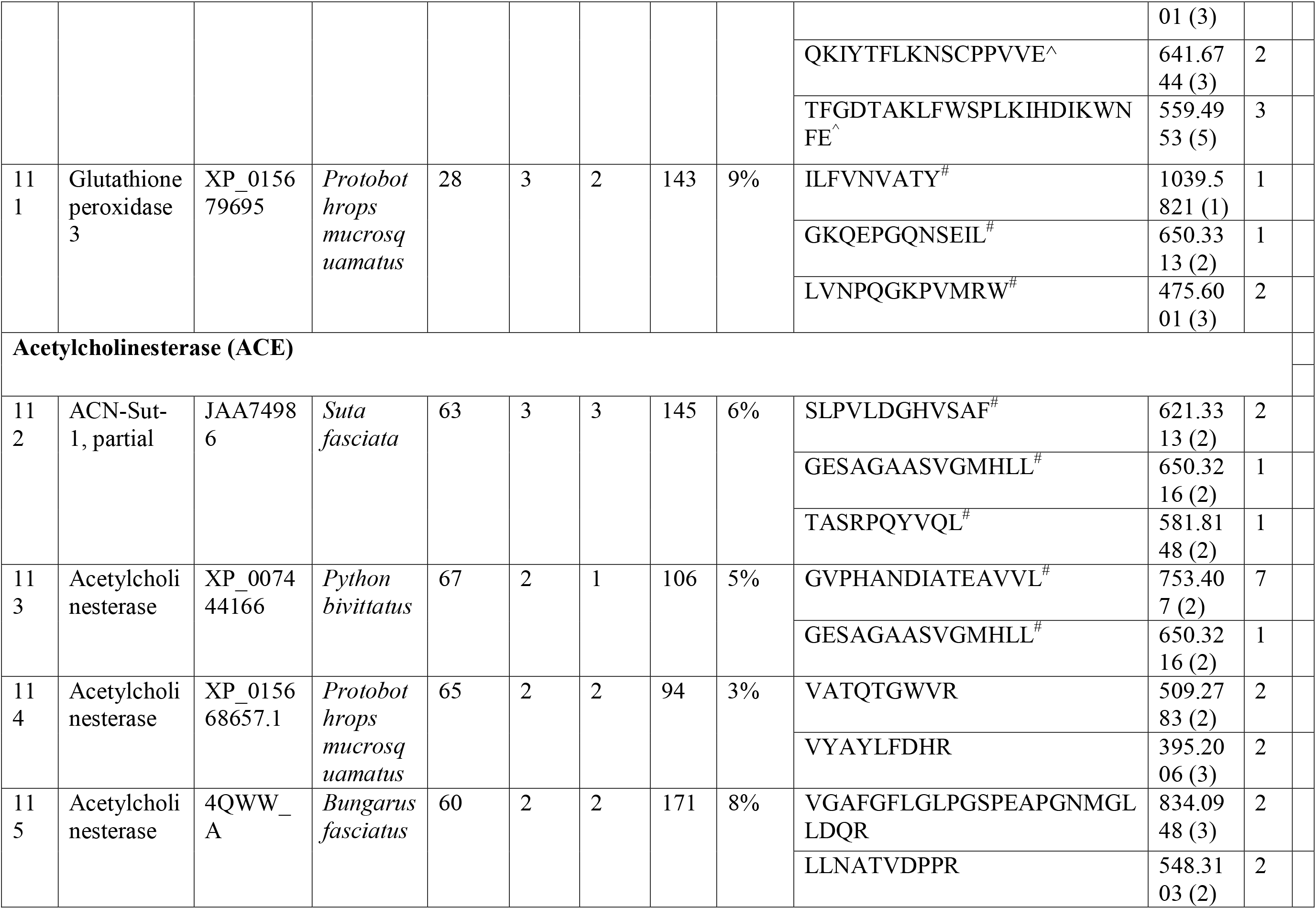
Proteins identified from crude NNwg venom after SDS-PAGE and HPLC fractionation. The proteins separated through SDS-PAGE and HPLC were subjected to in-gel and in-solution digestion using trypsin, chymotrypsin and V8 protease. The digests were then subjected to Q-TOF LC-MS/MS data collection and the obtained proteins were assigned to different venom protein families through Mascot and PEAKS analysis. #; peptides identified through chymotrypsin digestion, ^; peptides identified through V8 protease digestion, P#; peptides derived through PEAKS analysis after chymotrypsin digestion, P^; peptides derived through PEAKS analysis after V8 protease digestion. All the other peptides were obtained through trypsin digestion.

## Notes

### Competing Interest Statement

The authors have declared no competing interest.

