## Supplementary Table 1 for "Venomics and antivenomics of Indian spectacled cobra (*Naja naja*) from the Western Ghats"

**SUPPLEMENTARY FIGURES**

**Supplementary Figure 1. Peptides identified from *Naja naja* (NNwg) venom through *de novo* analysis**

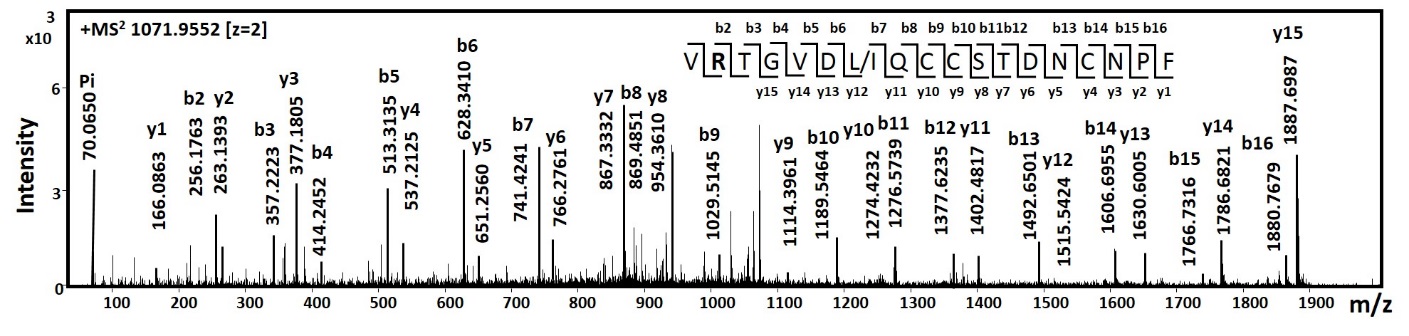

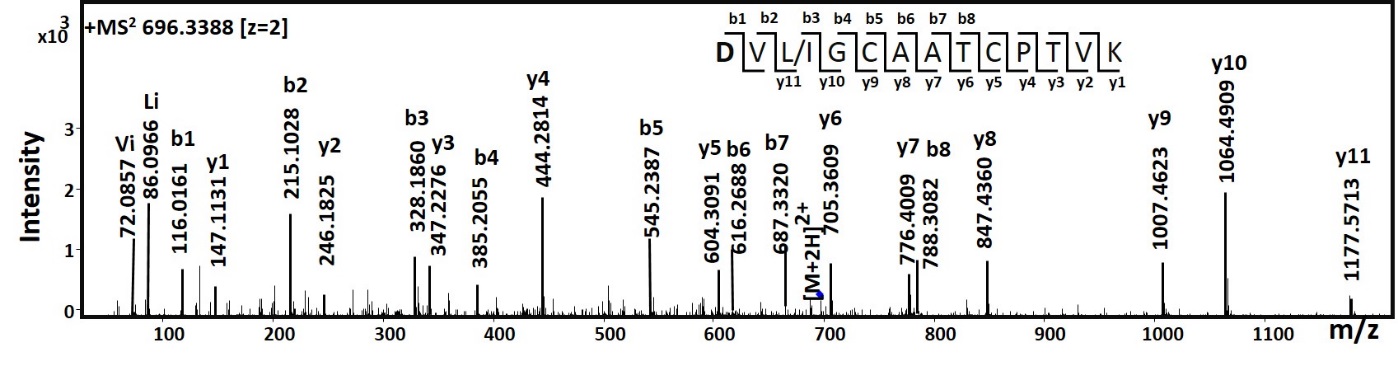

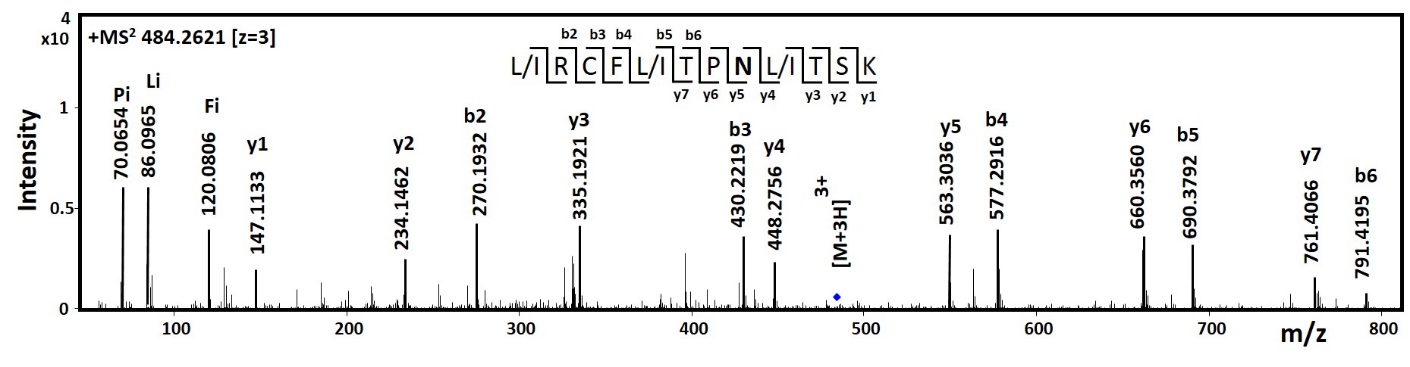

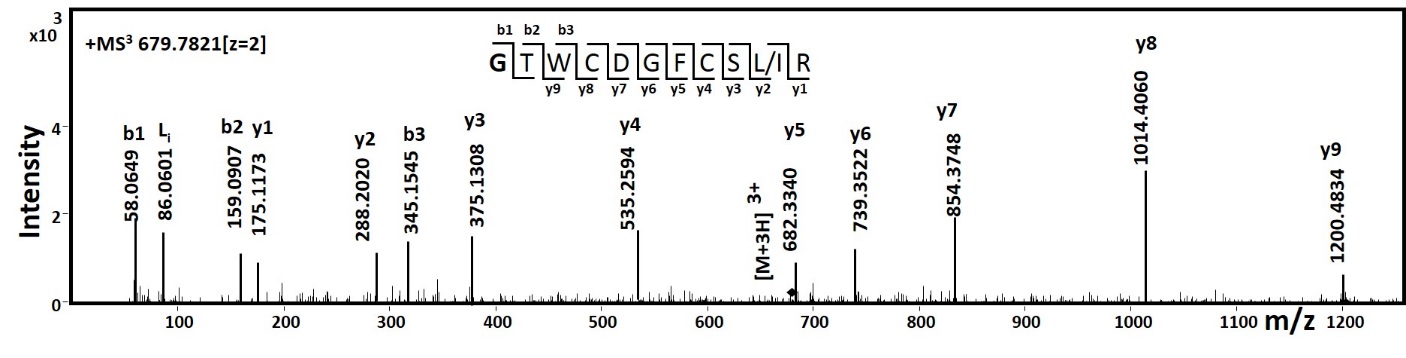

**1a (i)**

**1a (ii)**

**1a (iii)**

**1a (iv)**

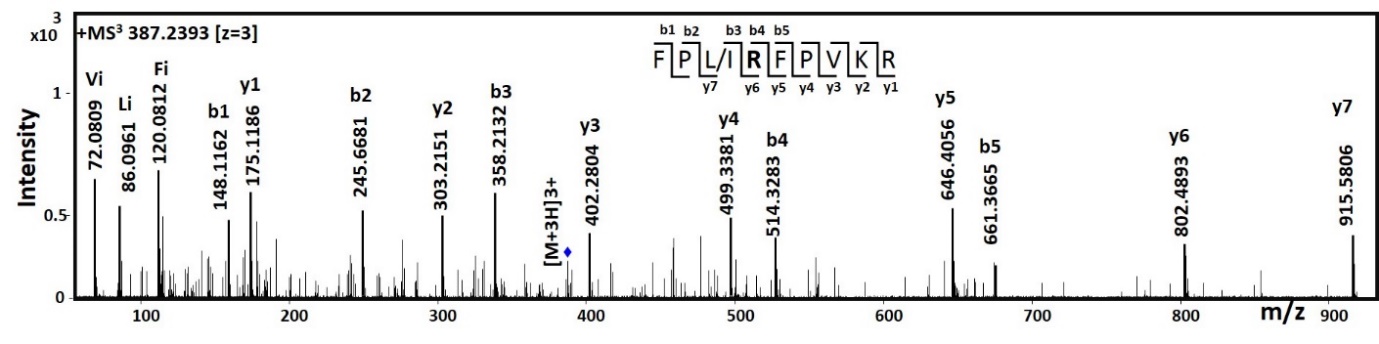

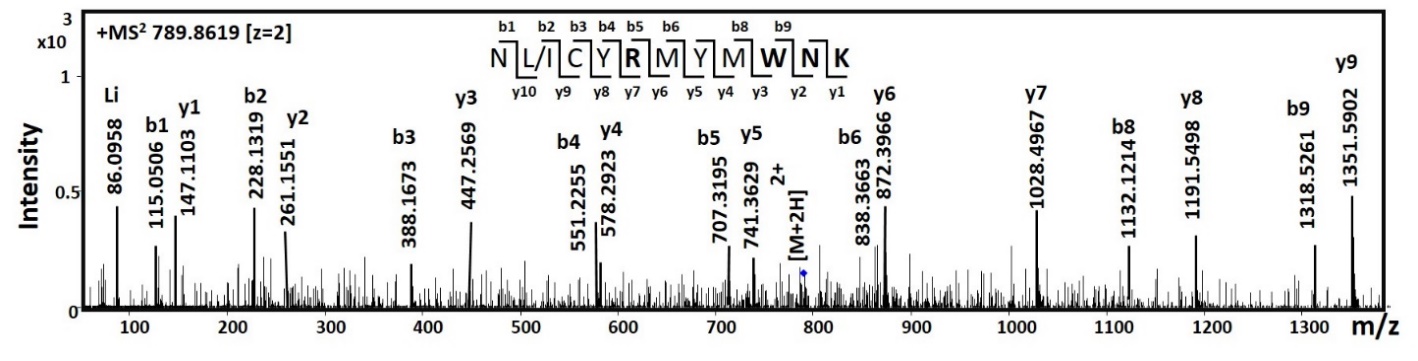

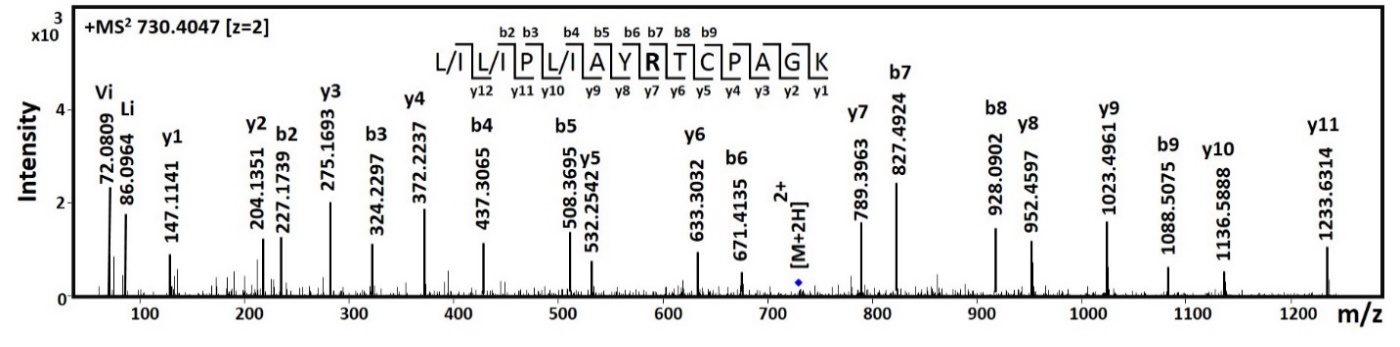

**1a (v)**

**1a (vi)**

**1a (vii)**

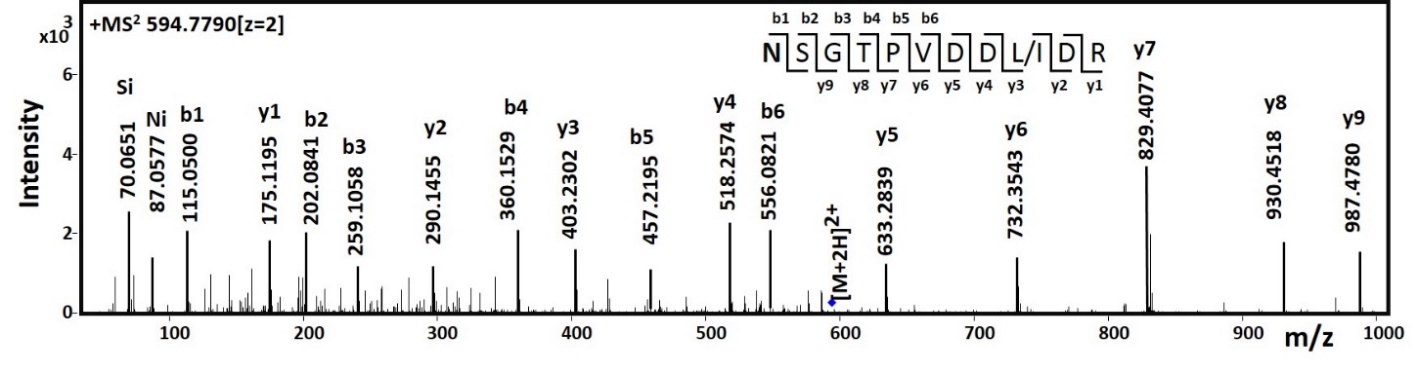

**1b**

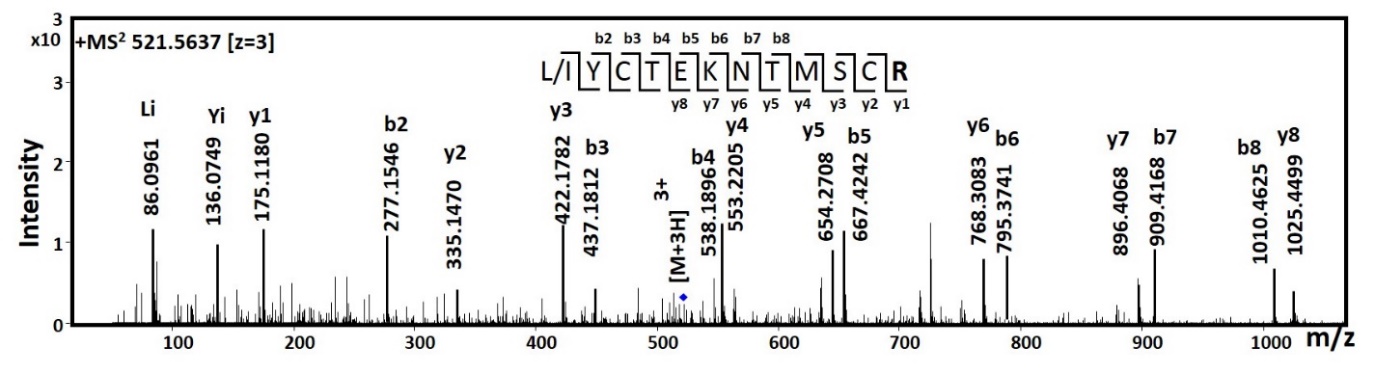

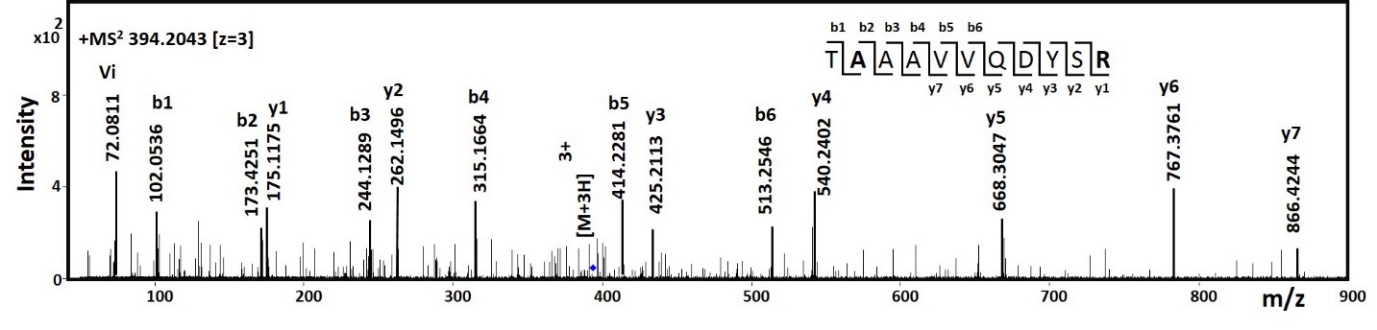

**1c (i)**

**1c (ii)**

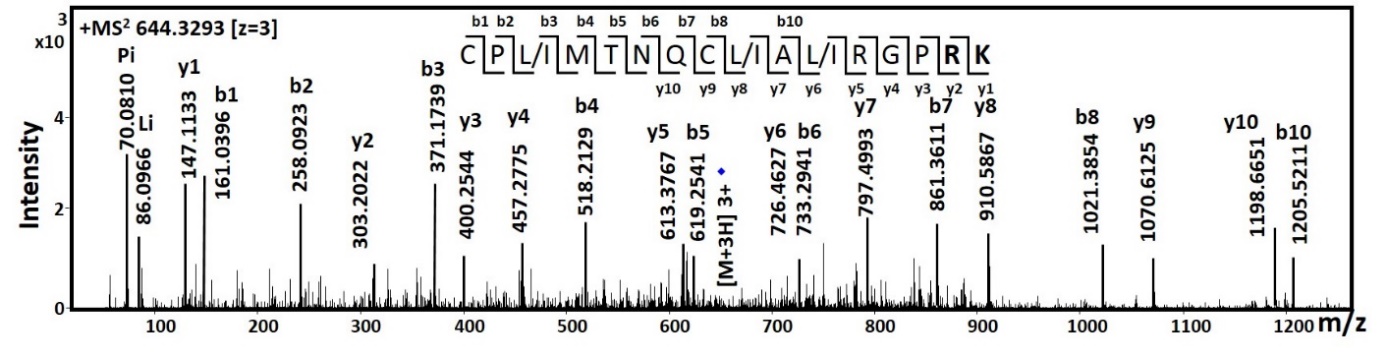

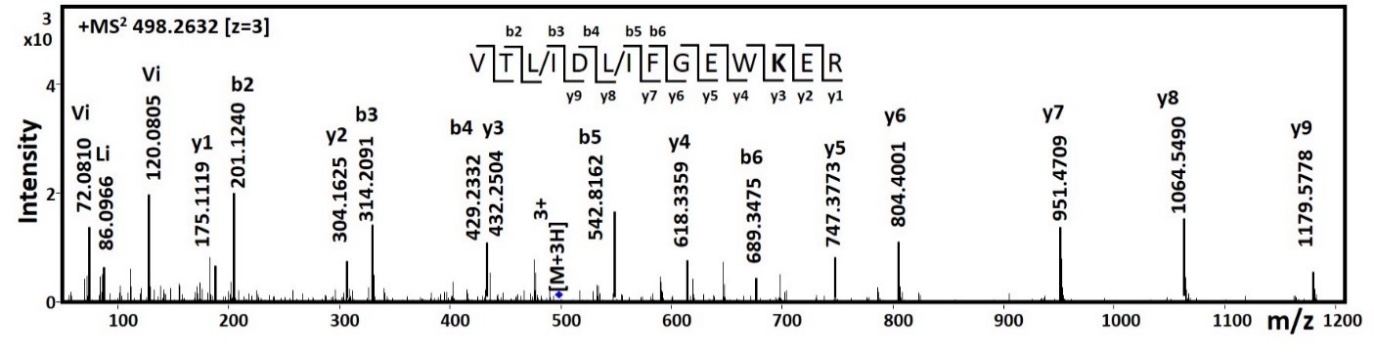

**1c (iii)**

**1c (iv)**

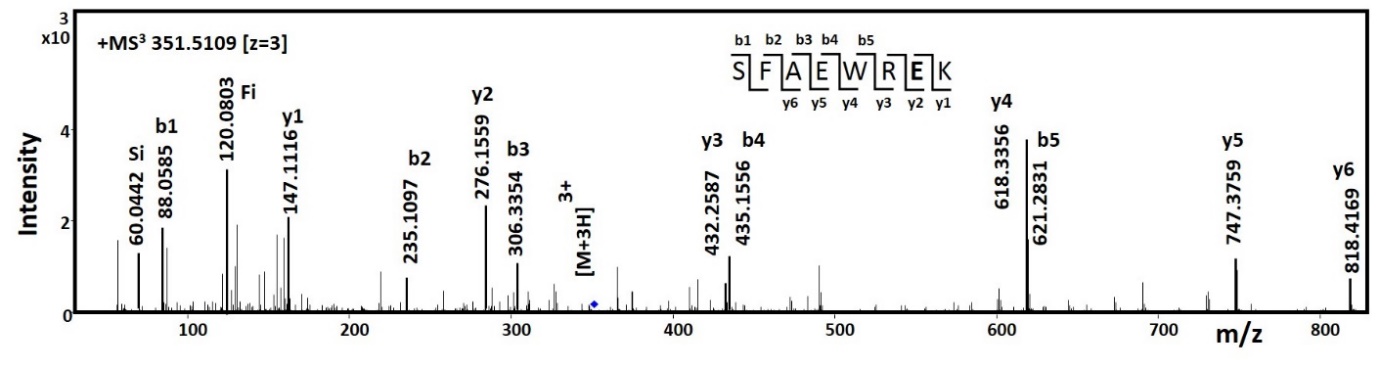

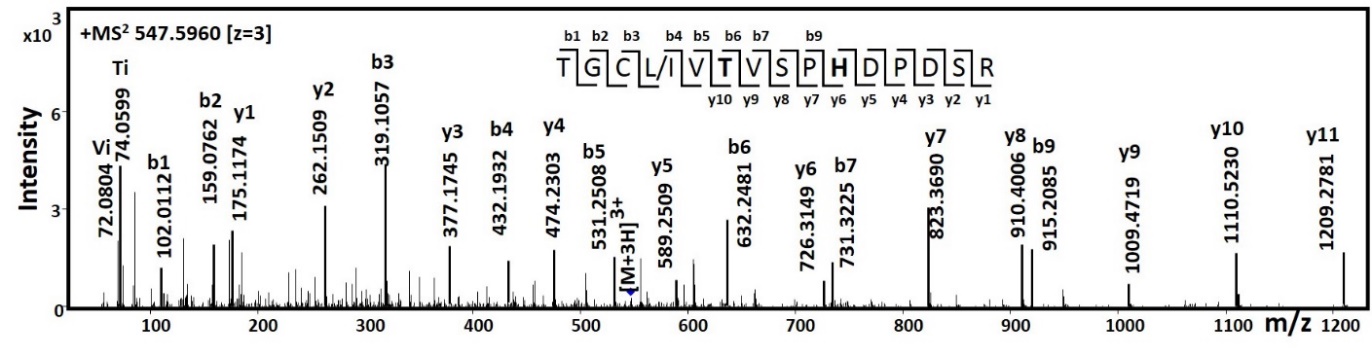

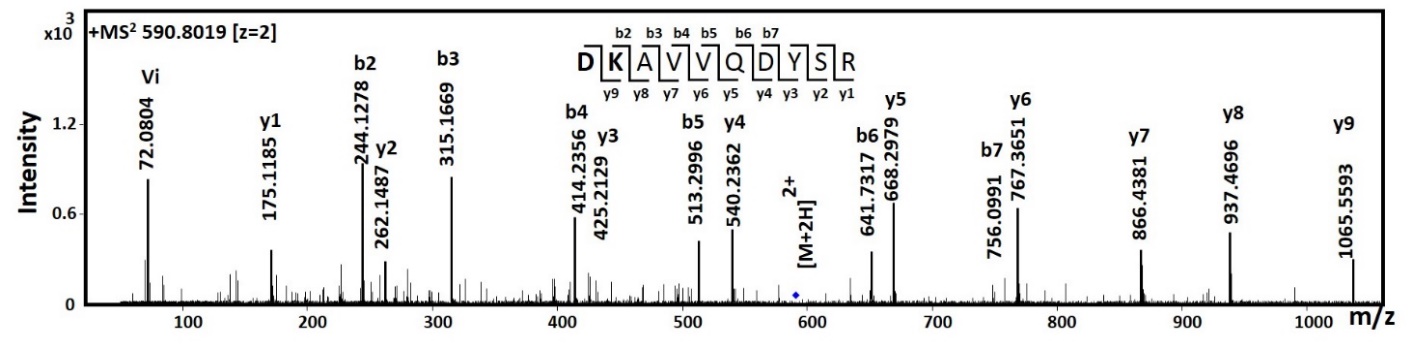

**1c (v)**

**1c (vi)**

**1c (vii)**

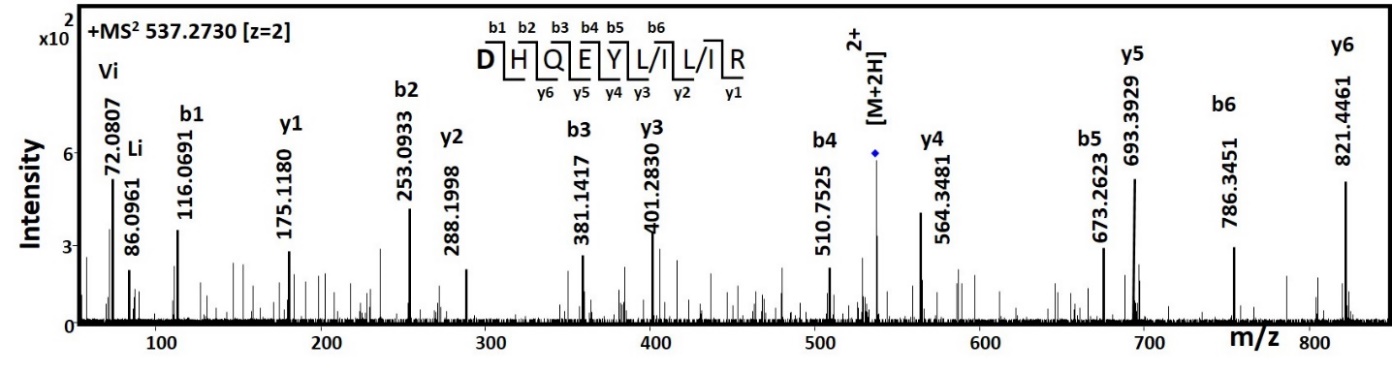

**1c (viii)**

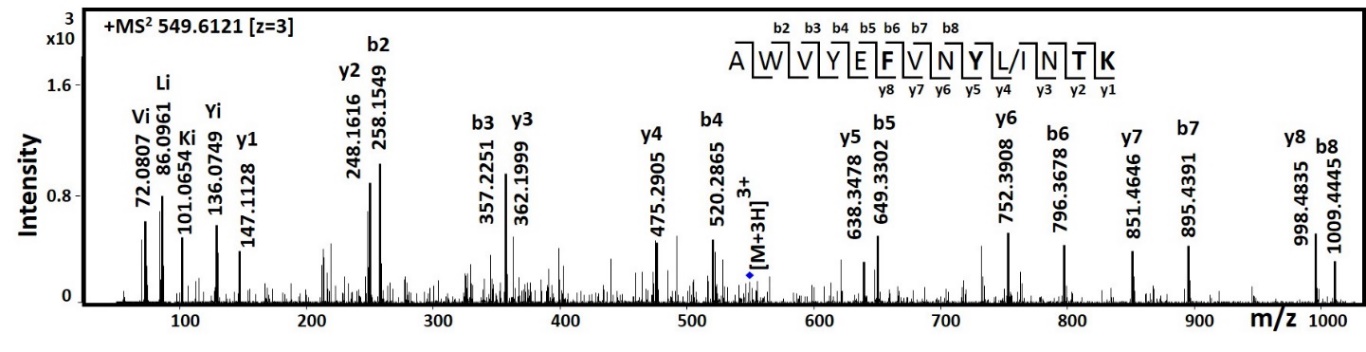

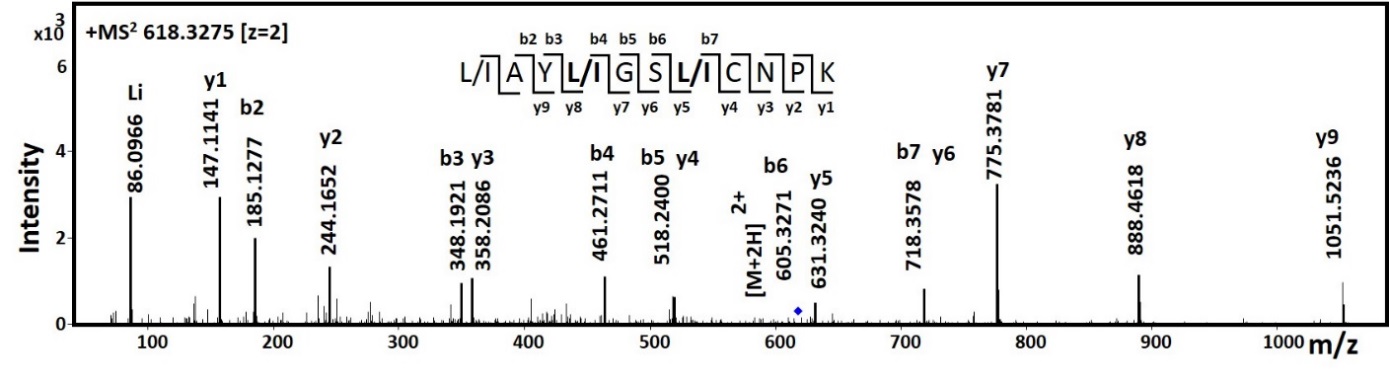

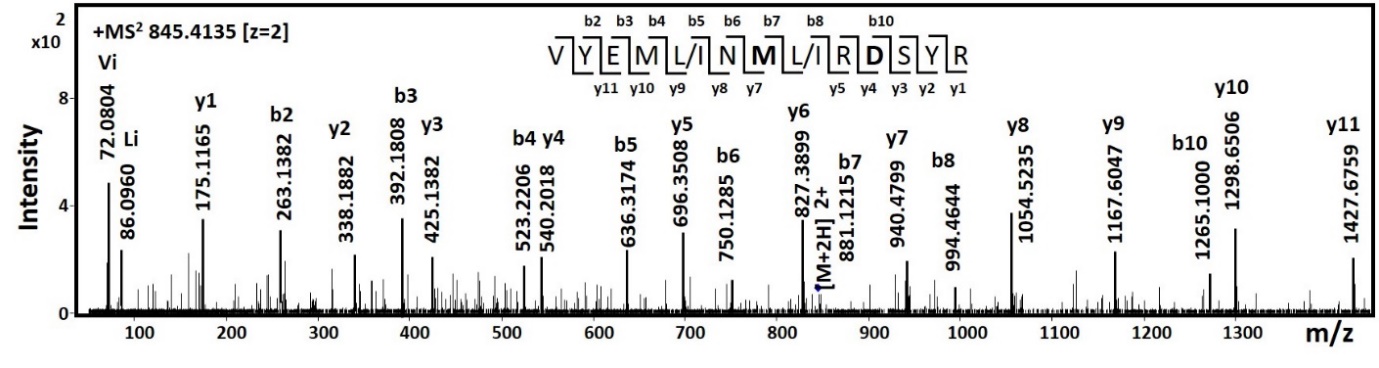

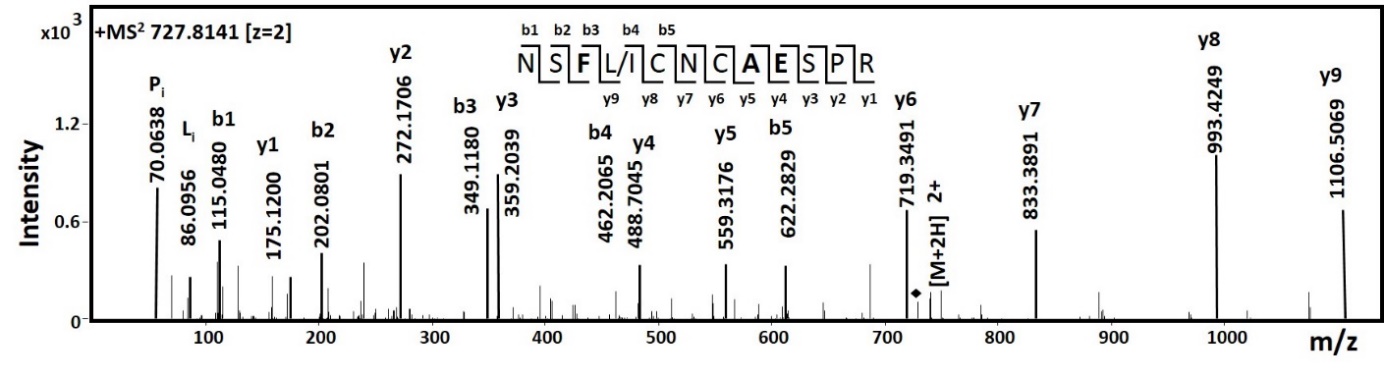

**1c (ix)**

**1c (x)**

**1c (xi)**

**1c (xii)**

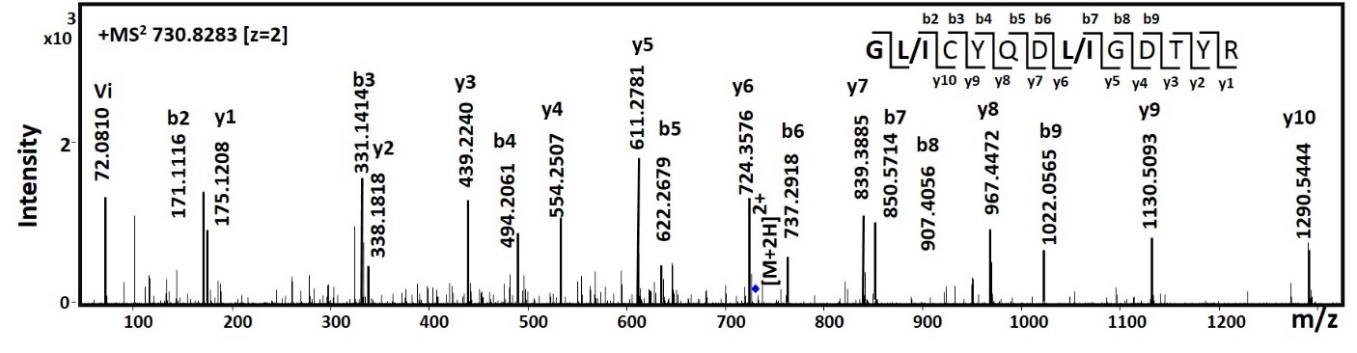

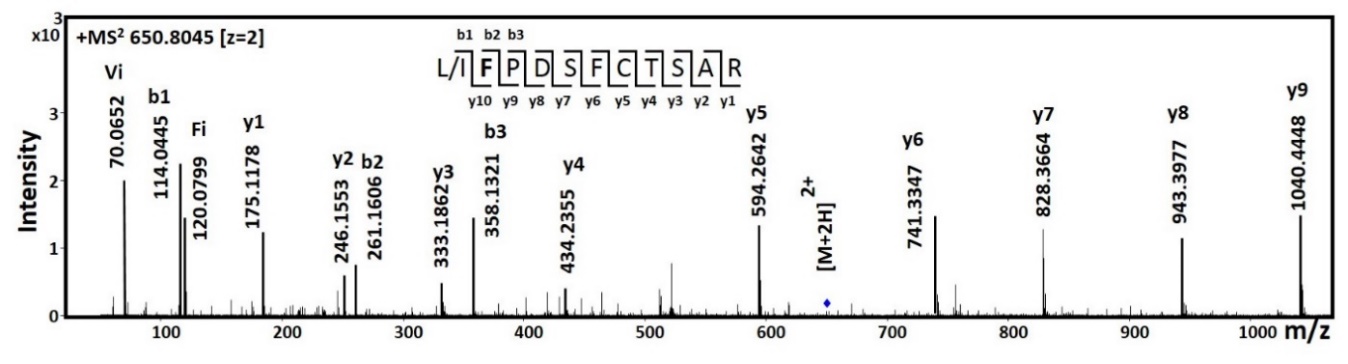

**1d (i)**

**1d (ii)**

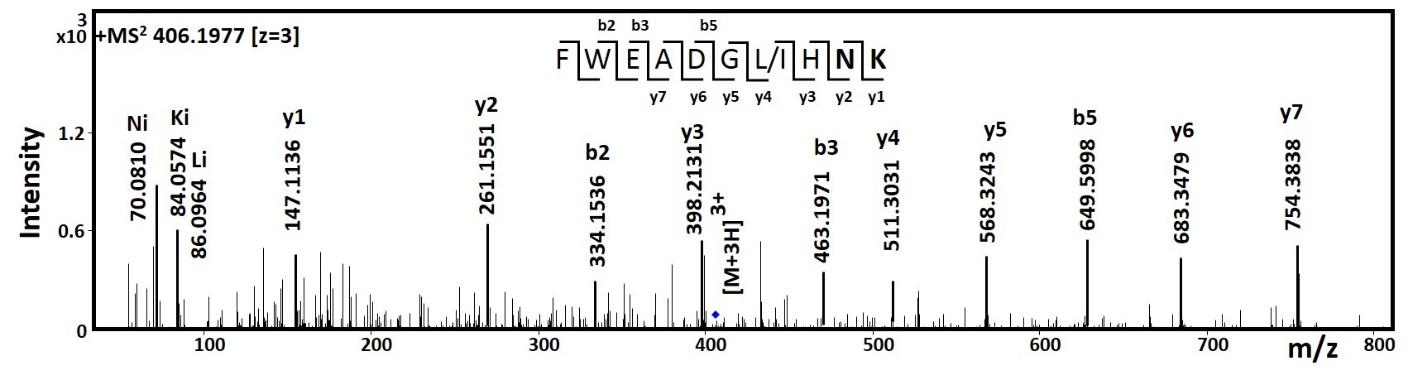

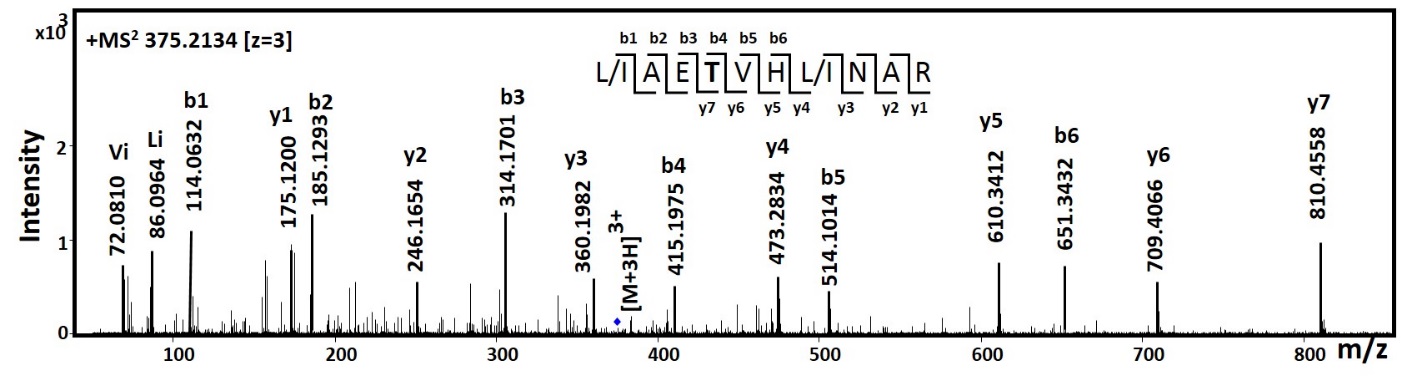

**1e (i)**

**1e (ii)**

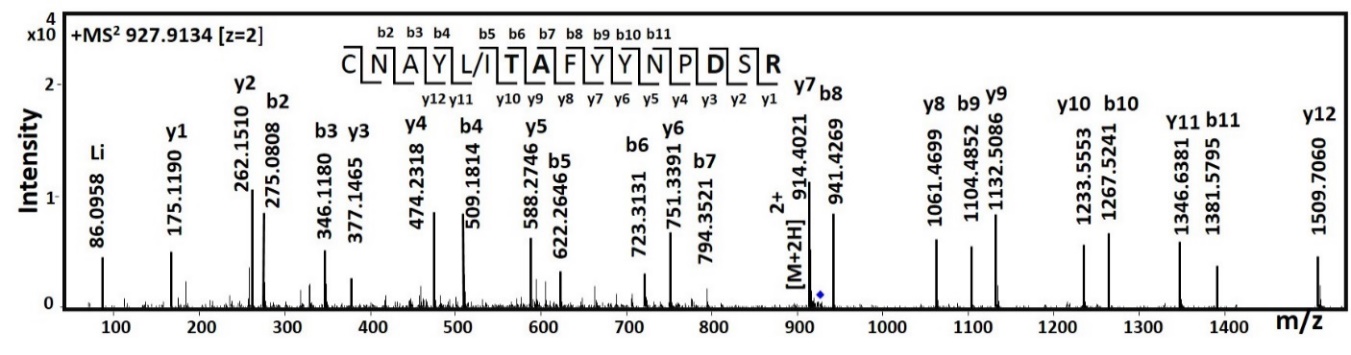

**1f**

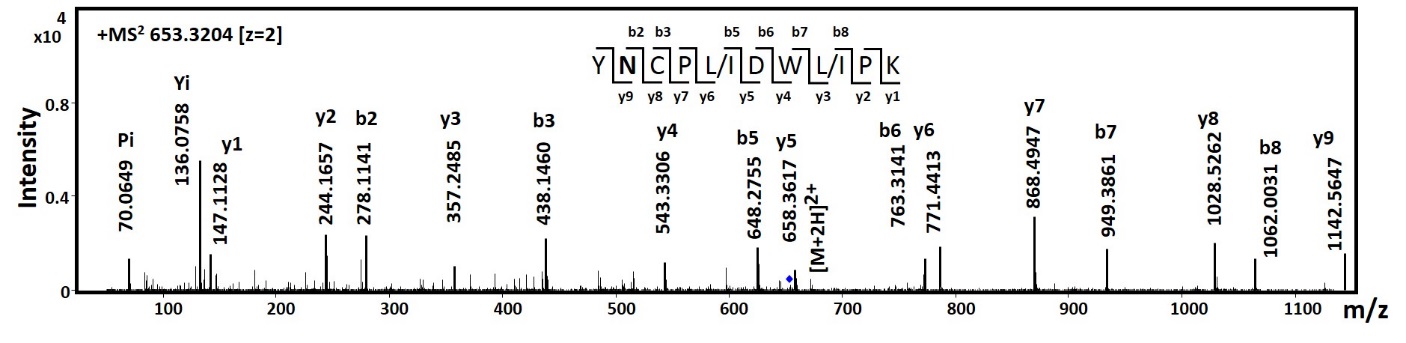

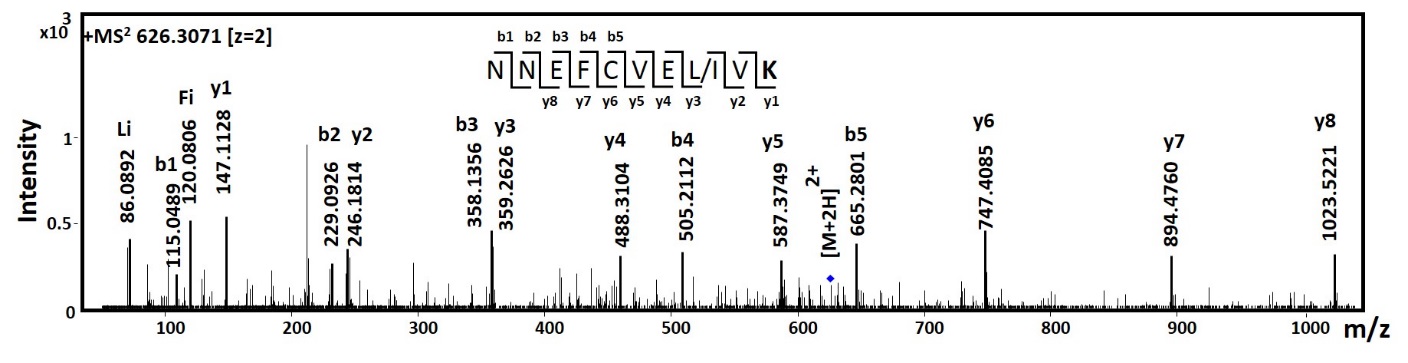

**1g (i)**

**1g (ii)**

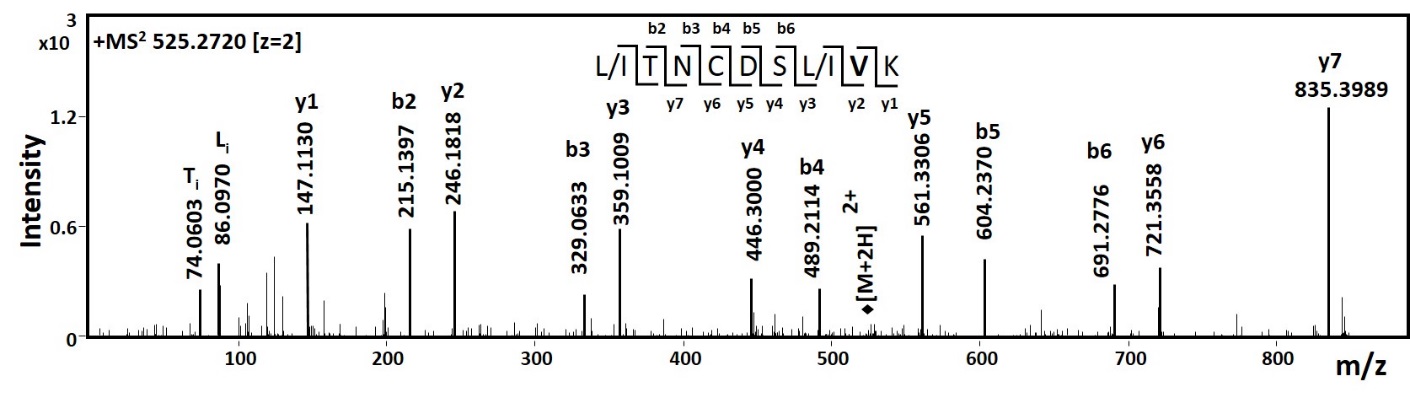

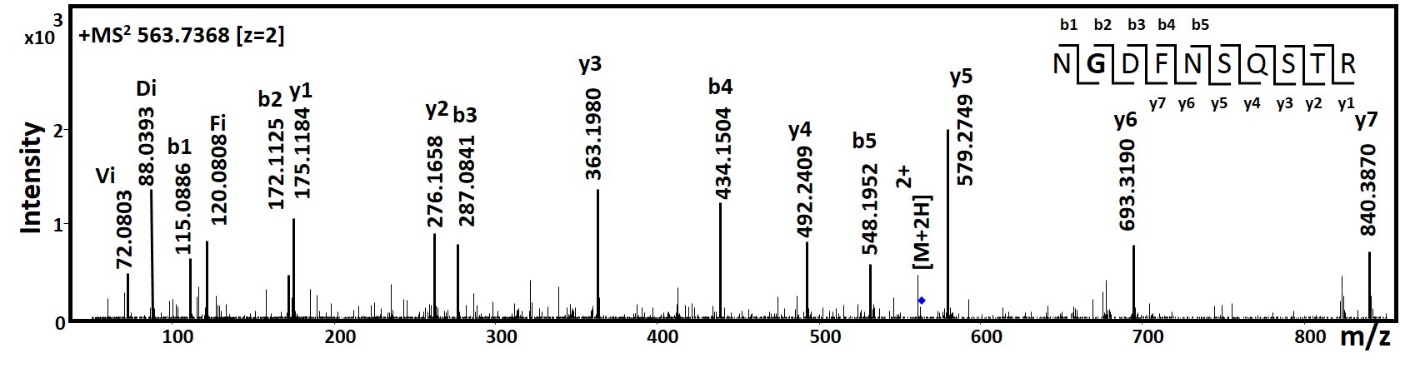

**1h (i)**

**1h (ii)**

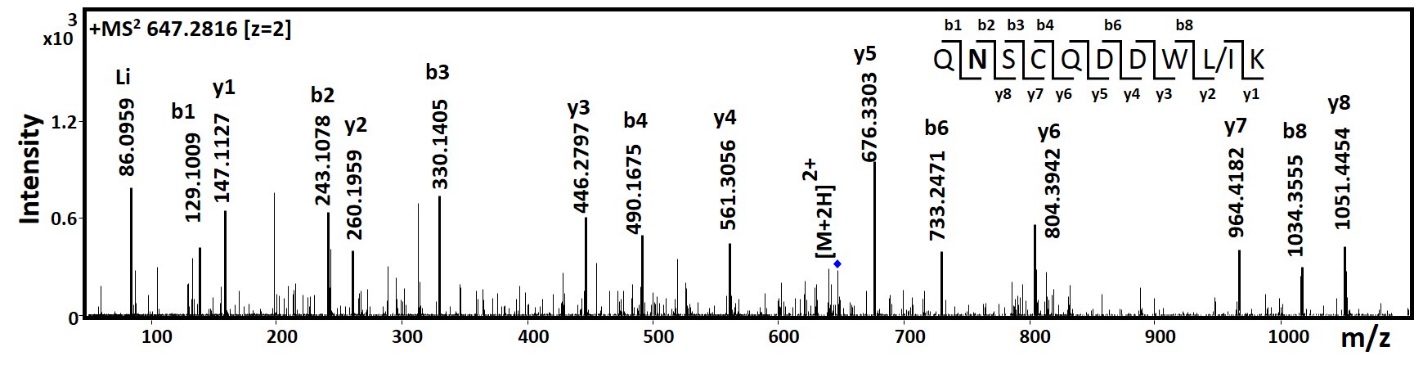

**1h (iii)**

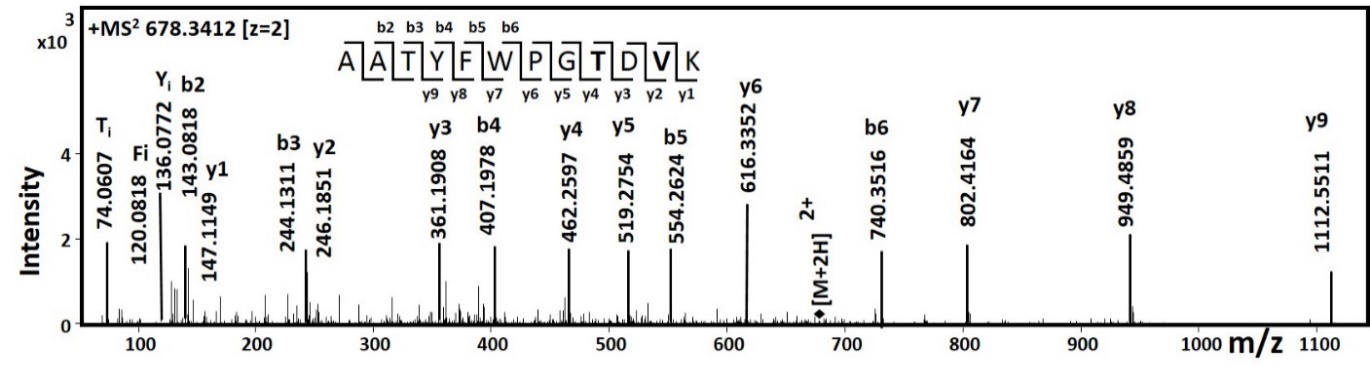

**1i**

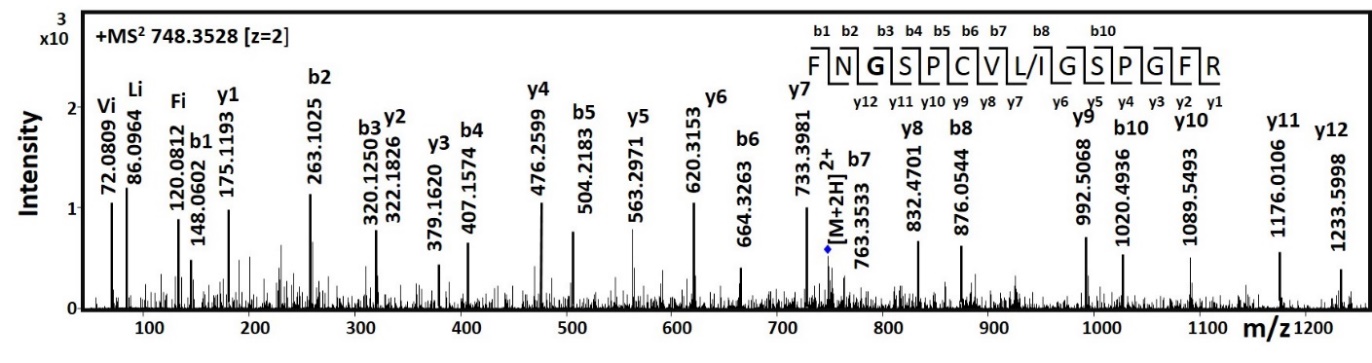

**1j**

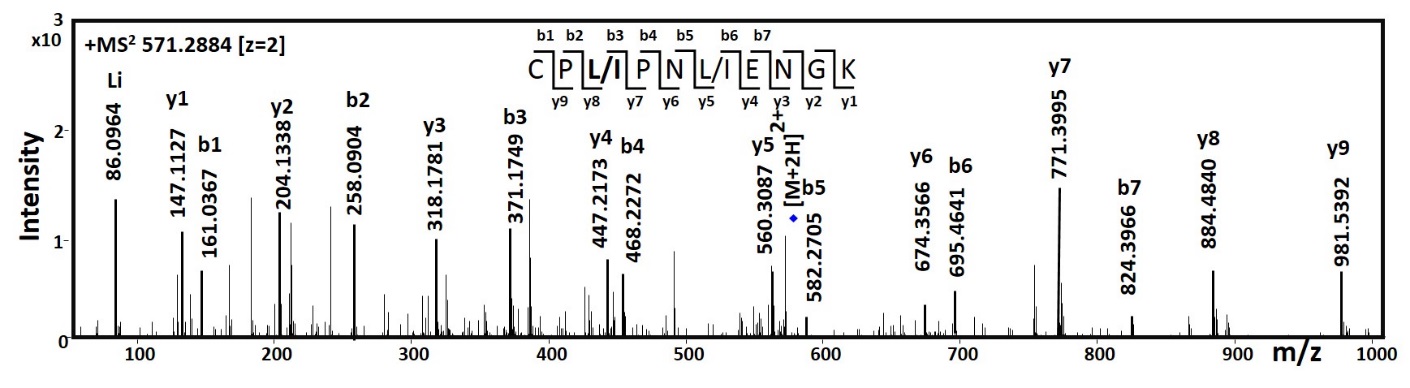

**1k (i)**

**1k (ii)**

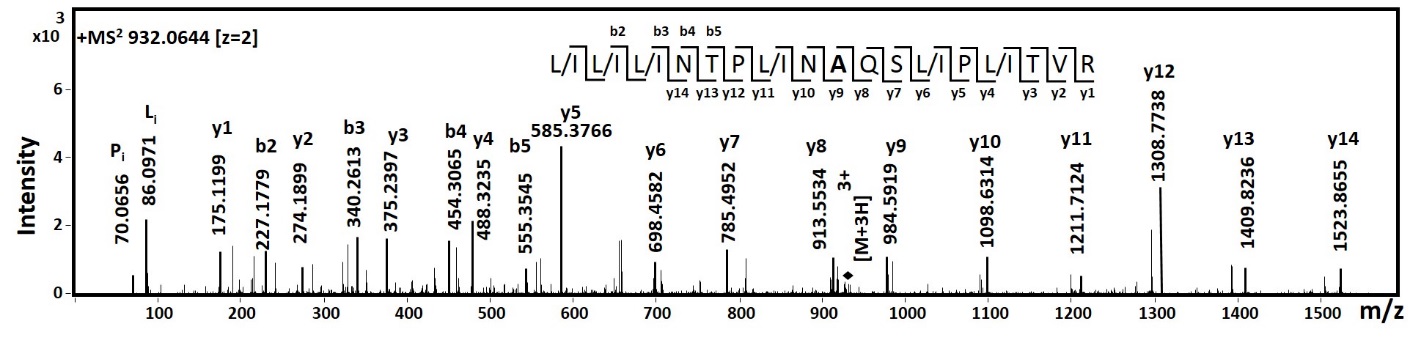

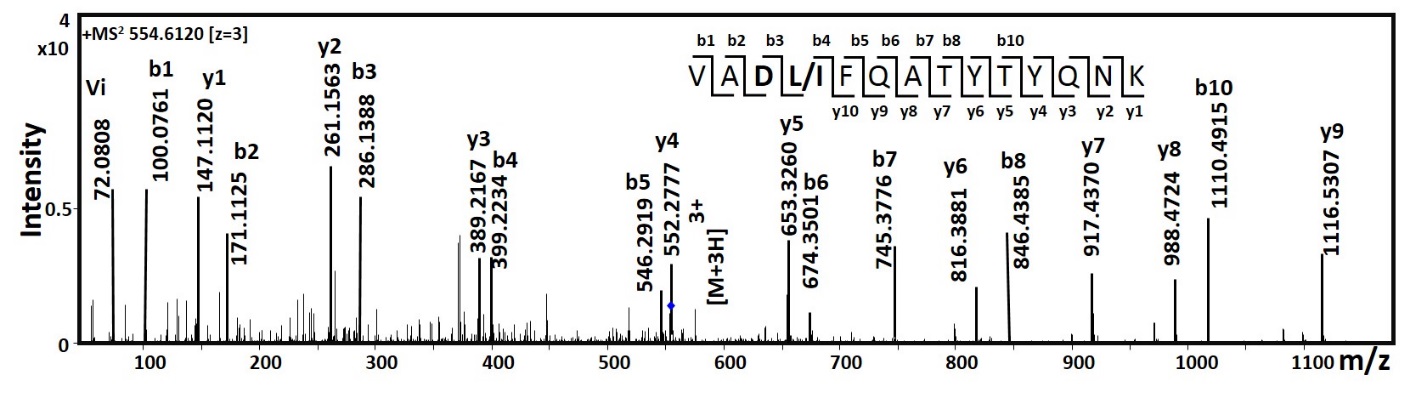

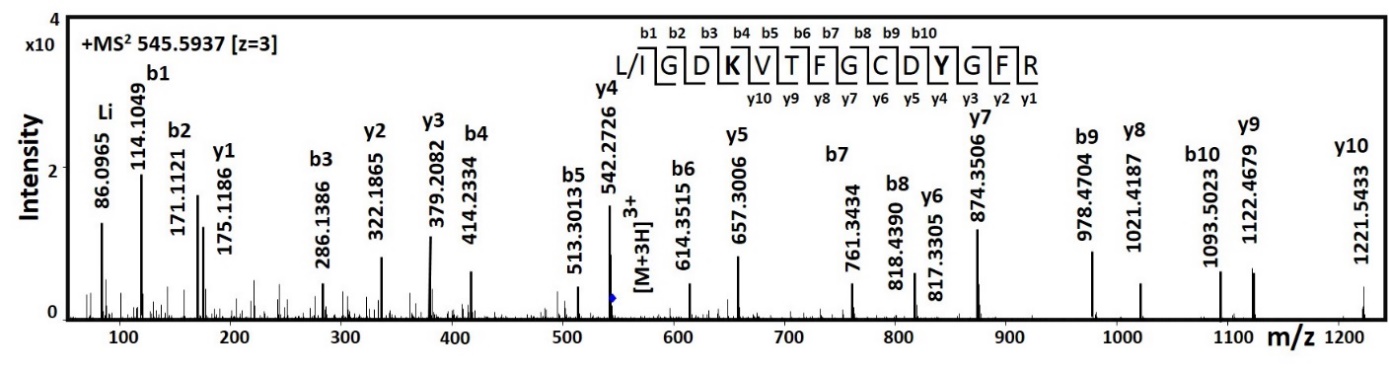

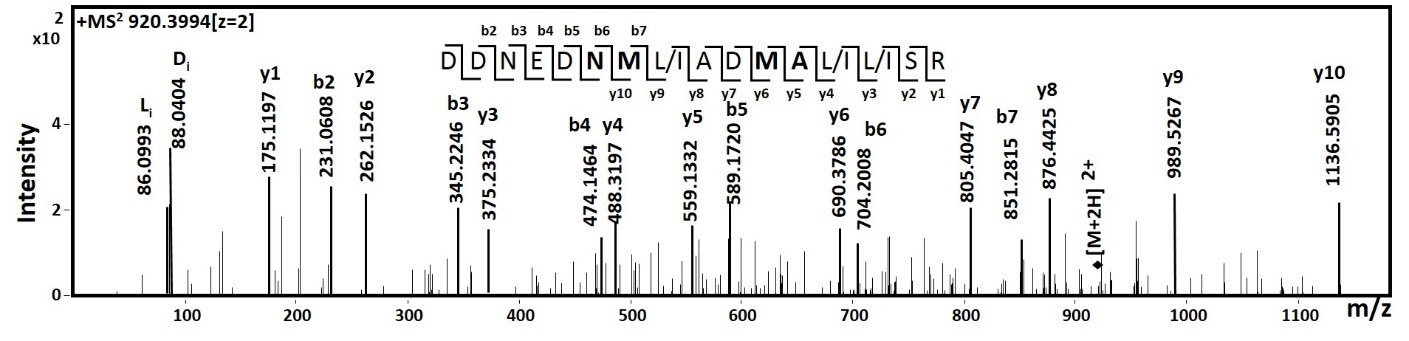

**1k (iii)**

**1k (iv)**

**1k (v)**

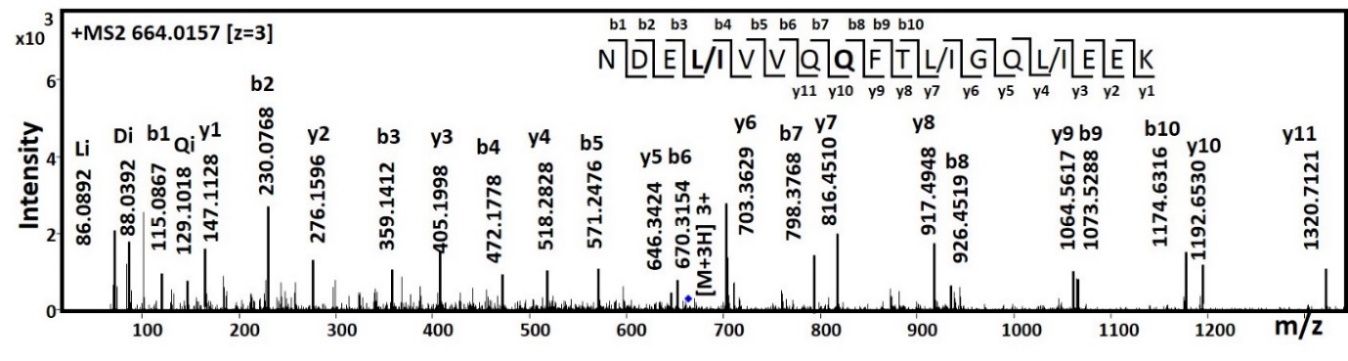

**1l**

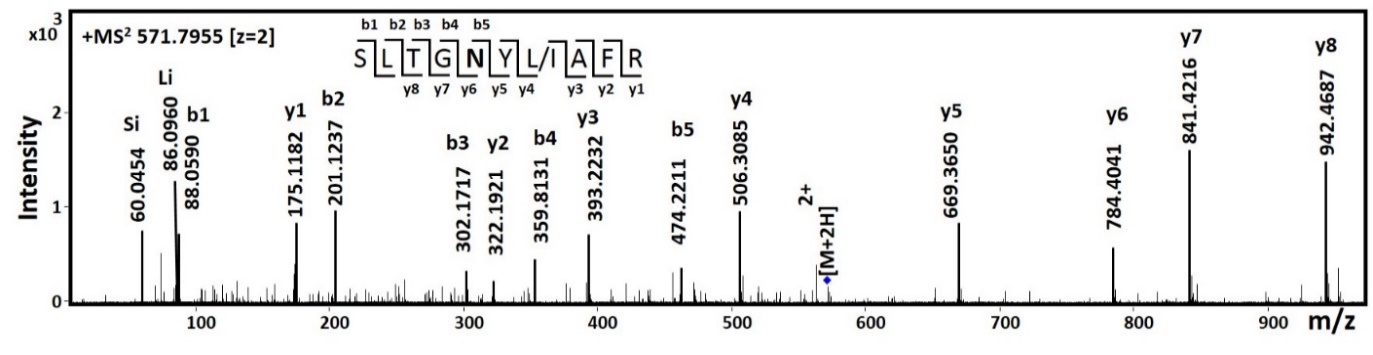

**1m**
